# When awareness outstrips performance: critical tests of subjective inflation under inattention

**DOI:** 10.1101/2025.07.03.661972

**Authors:** Karen J. Tian, Brian Maniscalco, Michael L. Epstein, Angela Shen, Olenka Graham Castaneda, Taiga Kurosawa, Jennifer A. Motzer, Emil Olsson, Emily E. Russell, Meghan E. Walsh, Juneau Wang, Tugral Bek Awrang Zeb, Richard Brown, Victor A. F. Lamme, Hakwan Lau, Biyu J. He, Jan W. Brascamp, Ned Block, David Chalmers, Megan A. K. Peters, Rachel N. Denison

**Affiliations:** Department of Psychological and Brain Sciences, Boston University; Department of Cognitive Sciences, University of California Irvine; Department of Humanities, LaGuardia Community College; The Graduate Center, City University of New York; Department of Psychology, University of Amsterdam; Department of Biomedical Engineering and Department of Intelligent Precision Healthcare Convergence, Sungkyunkwan University; RIKEN Center for Brain Science, Wako, Japan; New York University Grossman School of Medicine; Department of Psychology, Michigan State University; Department of Psychology, New York University; Department of Philosophy and Center for Neural Science, New York University

**Keywords:** spatial attention, awareness, subjective inflation, visual perception, consciousness

## Abstract

Visual experience can sometimes depart from visual performance, providing a powerful lens into the mechanisms generating conscious perception. In one proposed dissociation—subjective inflation—unattended locations in the periphery appear stronger than attended ones despite equated performance. Subjective inflation has played a central role in motivating theories of consciousness that reject the sufficiency of sensory signals for conscious perception. Yet the empirical basis for subjective inflation is limited. Here, in a large-scale adversarial collaboration, we conducted four simultaneously-replicated experiments testing the strength, character, and extent of subjective inflation under inattention. We used a new analytic approach to quantify inattentional inflation over full psychometric functions, beyond single matched-performance levels. We found robust inattentional inflation for contrast-dependent and texture-based perception, at and above the visual threshold. However at suprathreshold, we found inattentional inflation for the overall stimulus but not the specific feature relevant for performance. Finally, we establish the unifying principle that inattentional inflation occurs if and only if attention reduces performance thresholds more than visibility thresholds. Thus what we think we see may regularly exceed what we can visually discriminate, placing constraints on theories of conscious perception.

## Introduction

We generally think that when we attend to something, we will see it better. Indeed, attention enhances basic aspects of vision, including contrast sensitivity^1–5^ and spatial resolution^6–8^, to benefit performance on many tasks. Attention also enhances subjective appearance^9^, making attended items look stronger^10–15^, sharper^16,17^, and more perceptually organized^18^ than unattended ones. However, it has been reported that the effects of attention on objective and subjective aspects of perception do not proceed in lockstep^19,20^. Rather, at matched levels of task performance, unattended items can appear more visible than attended ones^21,22^.

Subjective inflation—the idea that our phenomenal experience can be stronger, sharper, more vivid, or otherwise “inflated” above what performance would suggest^23–27^—has been used to explain the apparent richness of the unattended periphery^28–30^, in spite of its poorer sensory processing^31^. Subjective inflation also suggests that unique mechanisms may underlie objective vs. subjective aspects of perception, and this dissociation has been taken by some as evidence supporting “higher-order” theories of conscious perception. This class of theories proposes that subjective experience arises from downstream metacognitive representations^30,32–37^, which can misrepresent the early-stage sensory processes governing visual performance. Against this view—and forming an enduring divide in theories of conscious perception^38,39^—first-order theories assert that sensory signals themselves are sufficient for subjective experience^40–42^.

Despite its theoretical implications, empirical evidence for subjective inflation is limited. While there are reports of subjective inflation for unattended (vs. attended locations)^21,22^, as well as for peripheral (vs. central) vision^23^ and crowded (vs. singleton) conditions^43^, these tests have been conducted only near detection thresholds and typically at only one or two matched levels of performance. Compelling evidence of subjective inflation specific to inattention (“inattentional inflation”) primarily comes from one study^21^. Others are partial, conceptual replications^28^. And yet some other studies have found only weak evidence^22^ or even counter-evidence^44^, leaving it unclear whether attention regularly dissociates objective and subjective aspects of perception in a way that could matter for everyday vision, where attention tends to be unevenly cast over clearly visible scenes.

The narrow scope of prior tests of subjective inflation was imposed in part by methodological constraints. One strategy to match performance across conditions of comparison is to physically titrate a single pair of stimuli. For example, in tests of inattentional inflation, the stimulus was made physically stronger in the unattended condition and weaker in the attended condition^21,22^ with the goal of equating stimulus processing across different levels of attention. However, this approach isolates measurements of inflation to a single performance level, one that may be suboptimal for revealing effects of inflation, and relies on statistical null effects to assume performance equivalence across conditions, such that what looked like inflation may have instead reflected small, non-significant performance differences^45,46^. It is unclear, then, whether the inconsistency in prior tests of inattentional inflation is a consequence of methodological limitations or the fragility of the phenomenon.

To justify subjective inflation as an explanation for the apparent richness of perception across the visual field and as a motivating pillar for higher-order theories of conscious perception^34–36^, it should withstand several key tests. First, subjective inflation should be tested beyond the visual detection of faint stimuli to understand whether inflation could underpin everyday, suprathreshold visual experience. Second, most demonstrations of inflation use grating stimuli^21,23,26^, leaving it unclear the extent to which inflation generalizes to the perception of visual properties beyond low-level contrast sensitivity. Third, studies of inflation have not asked participants about the visibility of the particular visual feature that is relevant to performance (the “task-relevant feature”), leaving open the possibility that previous findings of inflation were driven by objective and subjective reports accessing different stimulus information because the subjective task was relatively underspecified.

Here in four experiments (total n=120), conceived of in an adversarial collaboration ^47^ including first- and higher-order theorists and led by theory-neutral laboratories, we tested the robustness of inattentional inflation over a large range of stimulus and task conditions in a high-powered experimental design (see **Supplementary Note 1** for further details on the adversarial collaboration). We manipulated covert spatial attention using a central precue and measured its effects on the objective discrimination and subjective perception of peripheral stimuli. Across experiments, the stimuli were of different types (gratings and texture-defined figure-ground stimuli) and strengths, spanning threshold to suprathreshold regimes. We used gratings to facilitate comparison to previous tests of subjective inflation and figure-ground stimuli, which tap into the mid-level visual process of texture segmentation^6,48^, to test the phenomenon’s generalizability. To index subjective strength in suprathreshold regimes, in which detection measures would not be revealing, we created a comparative visibility task that asked participants to judge the apparent strength of a target relative to a learned reference. We used subjective measures that probed the visibility of the overall stimulus and of the task-relevant feature. Finally, to measure subjective inflation over full psychometric functions, we used a new analytic approach to quantify subjective reports over common ranges of performance using relative psychometric functions^46,49^, circumventing statistical and practical pitfalls in matching performance using single stimulus pairs.

In all experiments, we found strong and consistent inattentional inflation of the overall stimulus: withdrawing attention impaired the objective discrimination of visual features more than it reduced reports of apparent stimulus strength. Inflation of the stimulus not only extended beyond threshold regimes, but was more pronounced in suprathreshold regimes. However, when subjective reports specified the task-relevant feature, inflation was only found in threshold regimes. All effects of inattentional inflation were simultaneously replicated at two experimental sites and using two independent analytic pipelines. The results show that attention decouples objective and subjective reports in many, but not all, contexts. Peculiarly, when attention is withdrawn, we report seeing more than visual performance would suggest.

## Results

### Preregistration and simultaneous replication

All four experiments, including their methods, planned primary analyses, and theory predictions, were preregistered on Open Science Framework (preregistration document^50^: https://osf.io/p3erc; detailed documents: https://osf.io/yur93/). For simultaneous replication, all experiments and analyses were conducted in parallel at two experimental sites. We refrained from sharing data between sites and performing the critical analyses for theory predictions (i.e., quantifying subjective reports as a function of objective performance across varying levels of attention) until both sites completed data collection for an experiment. All main results were consistent across both sites and were confirmed using two independent analysis pipelines. For simplicity, we report the combined data across sites in the main text, with additional site-specific details provided in the Supplementary Material.

### Task protocol

In four experiments, human participants (n=30 per experiment with ∼3000 trials per participant) performed a spatial attentional cueing task **(Figure 1, Supplementary Figure 1)**. On each trial, participants viewed up to four peripheral targets, which independently varied across 7 strengths, defined separately for two stimulus types. Targets were either contrast-defined gratings in noise (Experiments 1 and 2) or texture-defined figure-ground ovals, with textures composed of line elements (Experiments 3 and 4). Targets were parametrically manipulated in strength—via grating contrast or texture line length—to span near-threshold (Experiments 1 and 3) to suprathreshold (Experiments 2 and 4) regimes **(Figure 1b)**. Stimulus strengths were titrated per participant to be near detection thresholds (see Methods, Thresholding) or fixed across participants to be at suprathreshold values (see Methods, Stimuli). To further increase the grating visibility in the suprathreshold experiments, the noise contrast was decreased from 50% to 20%. The 2x2 design of stimulus type and strength regime, along with presenting 7 levels of stimulus strength within each experiment, equipped us to test for subjective inflation across broad stimulus conditions.

**Figure 1.**
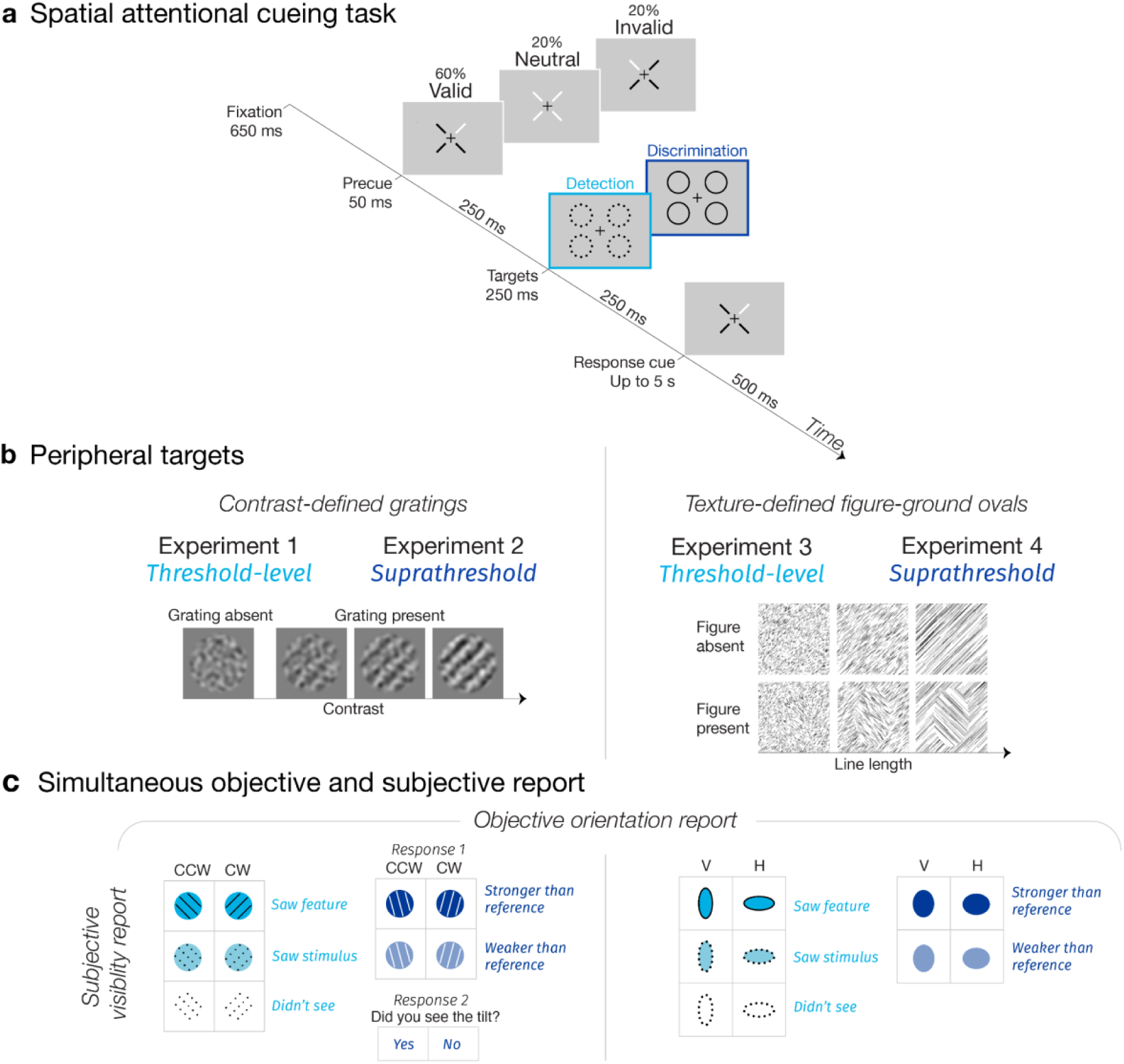
Spatial attentional cueing task. **a)** Trial timeline. A central precue before the targets directed attention covertly to one or all four target locations. A response cue after the targets indicated which peripheral quadrant to report, which most often matched the precued location (of the 80% of trials in which a single location was precued, the precue was 75% valid). **b)** Targets were either contrast-defined gratings (Experiments 1 and 2) or texture-defined figure-ground ovals (Experiments 3 and 4), which varied independently across 7 strengths in each quadrant. **c)** Participants made an objective orientation report and a subjective visibility report about the response-cued quadrant. The subjective visibility report was either to detect a threshold strength stimulus (Experiments 1 and 3) or to compare the visibility of a suprathreshold stimulus to a learned reference (Experiments 2 and 4). In three experiments (1-3), participants also specified whether or not they subjectively saw the task-relevant feature: the orientation of the grating or oval.

Before the targets appeared, a central precue directed covert attention to one or all target locations while central fixation was monitored **(Figure 1a)**. After the targets disappeared, a response cue indicated which single peripheral location to report. On most trials (60%), the response cue and precue matched (“valid” condition), so participants had incentive to direct attention to the precued location. On some trials (20%), the precue misdirected attention to a location the participant did not have to report (“invalid” condition). When the precue was spatially uninformative (20% of trials, “neutral” condition), participants were asked to distribute attention across all possible target locations.

Participants supplied simultaneously 1) an objective orientation report and 2) a subjective visibility report about the response-cued quadrant **(Figure 1c)**. The objective report was to discriminate the target orientation (i.e., “Was the grating tilted counterclockwise or clockwise from vertical?” or “Was the oval figure oriented vertically or horizontally along its major axis?”). In the threshold experiments, the gratings were orthogonally oriented (±45°) and figure-ground ovals clearly elongated (5° by 3° aspect ratio), so that the difficulty of orientation discrimination was driven largely by target detectability. In the suprathreshold experiments, a stimulus feature (the grating tilt or oval aspect ratio) was titrated per participant to ensure orientation discrimination remained challenging even when the target was easily detectable.

The subjective report was either to detect a near-threshold target (“Did you see or not see a stimulus?”, Experiments 1 and 3) or to compare the strength of a suprathreshold target to a learned reference (“Was the stimulus stronger or weaker than the reference?”, Experiments 2 and 4). We instructed participants to report subjective strength based on how visibly the gratings appeared to stand out from the noise or how visibly the texture-defined figures appeared to “pop-out” from the background, with excerpts of the instructions provided in the Supplementary Methods. In three experiments (1-3), participants additionally specified whether or not they saw the feature relevant for the objective task: the orientation of the grating or oval. Even if a target was not physically presented or consciously detected, participants nonetheless made a forced-choice guess about the orientation, so that every trial provided a concurrent objective and subjective measure, using a single key press to preclude post-choice effects^51,52^, except in Experiment 2, in which participants reported the feature visibility judgment using a second key press.

### Objective and subjective reports increased with stimulus strength

We first assessed objective performance (**Figure 2a**) and subjective visibility reports (**Figure 2b, 3b**) separately as functions of stimulus strength. In all experiments, objective performance was the proportion correct in identifying the target orientation—p(correct discrimination). Chance performance was 50%. We operationalized subjective visibility as the proportion of trials participants reported seeing the target—p(“saw stimulus”)—which was presented half of the time in the detection experiments. In the suprathreshold experiments, we indexed subjective strength by asking participants to rate how strong the target appeared relative to a middle reference strength they had learned before the main experiment (see Methods, Reference training) and measured the proportion of trials they reported the test stimulus as stronger than the reference—p(“test stronger”). We similarly quantified the proportion of trials in which participants reported seeing the task-relevant feature—p(“saw feature”) **(Figure 3b)**.

**Figure 2.**
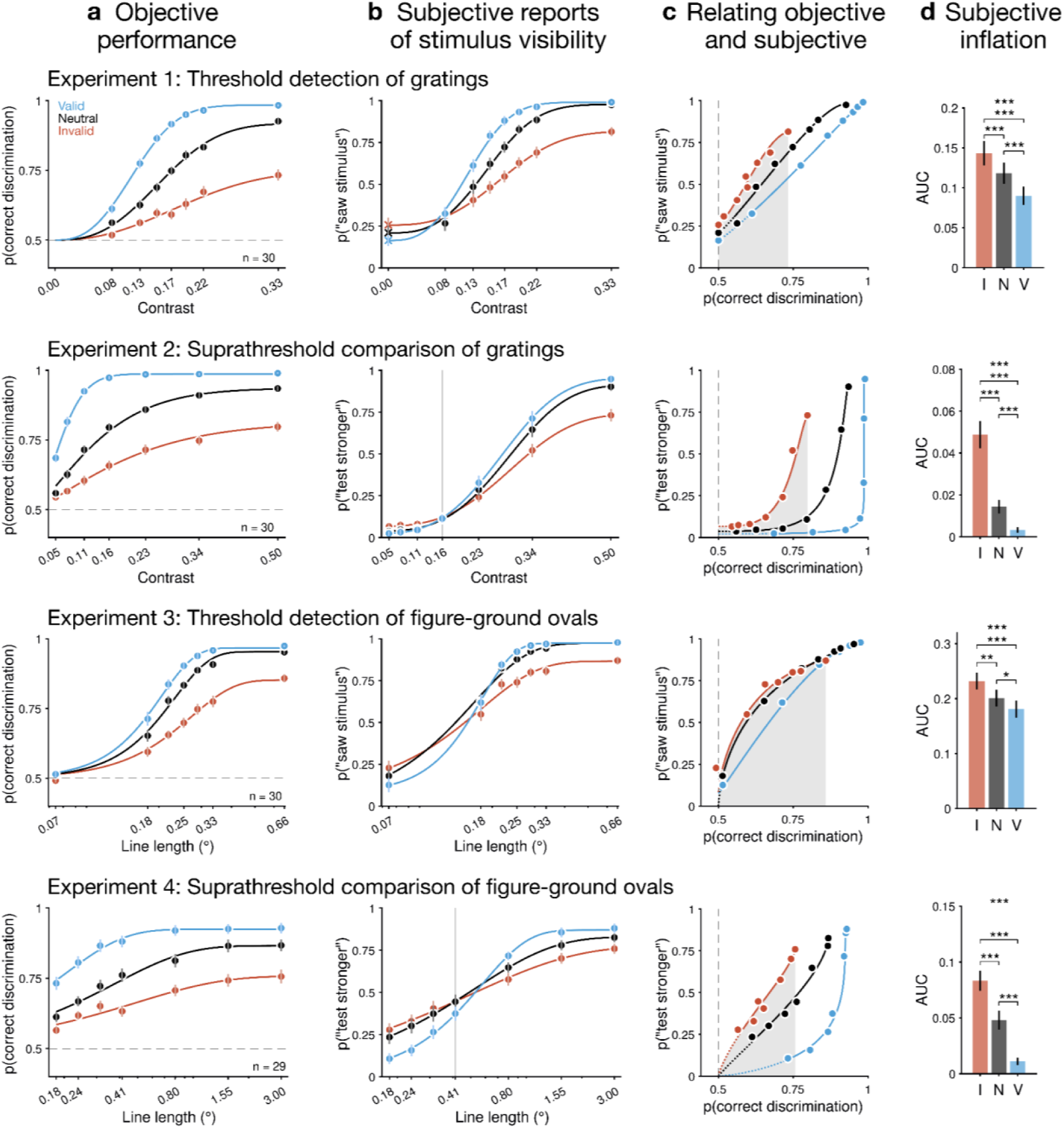
Inattentional inflation of stimulus visibility. In four experiments (one per row), **a)** objective performance increased with stimulus strength and with attention (blue = valid, gray = neutral, red = invalid). Contrast values indicate the grating contrast alone, not including the noise contrast, which was higher (50%) in Experiment 1 and lower (20%) in Experiment 2 to allow the gratings to be more visible. **b)** Subjective reports of stimulus visibility increased with stimulus strength, and attention increased the sensitivity to stimulus strength. Apart from when the grating contrast was zero in Experiment 1 (marked with x’s), the data show responses to when the target was present. Target absent data were also collected in Experiment 3, plotted in Supplementary Figure 4. Solid vertical lines mark the reference strength in the suprathreshold experiments. **c)** Subjective reports increased with objective performance. As a function of performance, subjective reports were higher for unattended than attended stimuli (subjective inflation). Gray-shaded regions mark the shared range of performance across attention conditions. For a range of matched performance, **d**) the area under the relative psychometric function^46^ (AUC) was greater for unattended than attended stimuli (repeated measures ANOVA per experiment, all p<0.001). Group means are fit with Weibull functions (a,b) or their corresponding relative psychometric function (c) for visualization. The portion of the fit spanning the range of performance observed between the lowest and highest stimulus strengths is shown as a solid line; the portion extrapolated to chance performance is dotted. \**p*<0.05, \*\**p*<0.01, \*\*\**p*<0.001. Data (total n=119; Experiments 1-3 each n=30, Experiment 4 n=29) are presented as mean values ±1 SEM.

**Figure 3.**
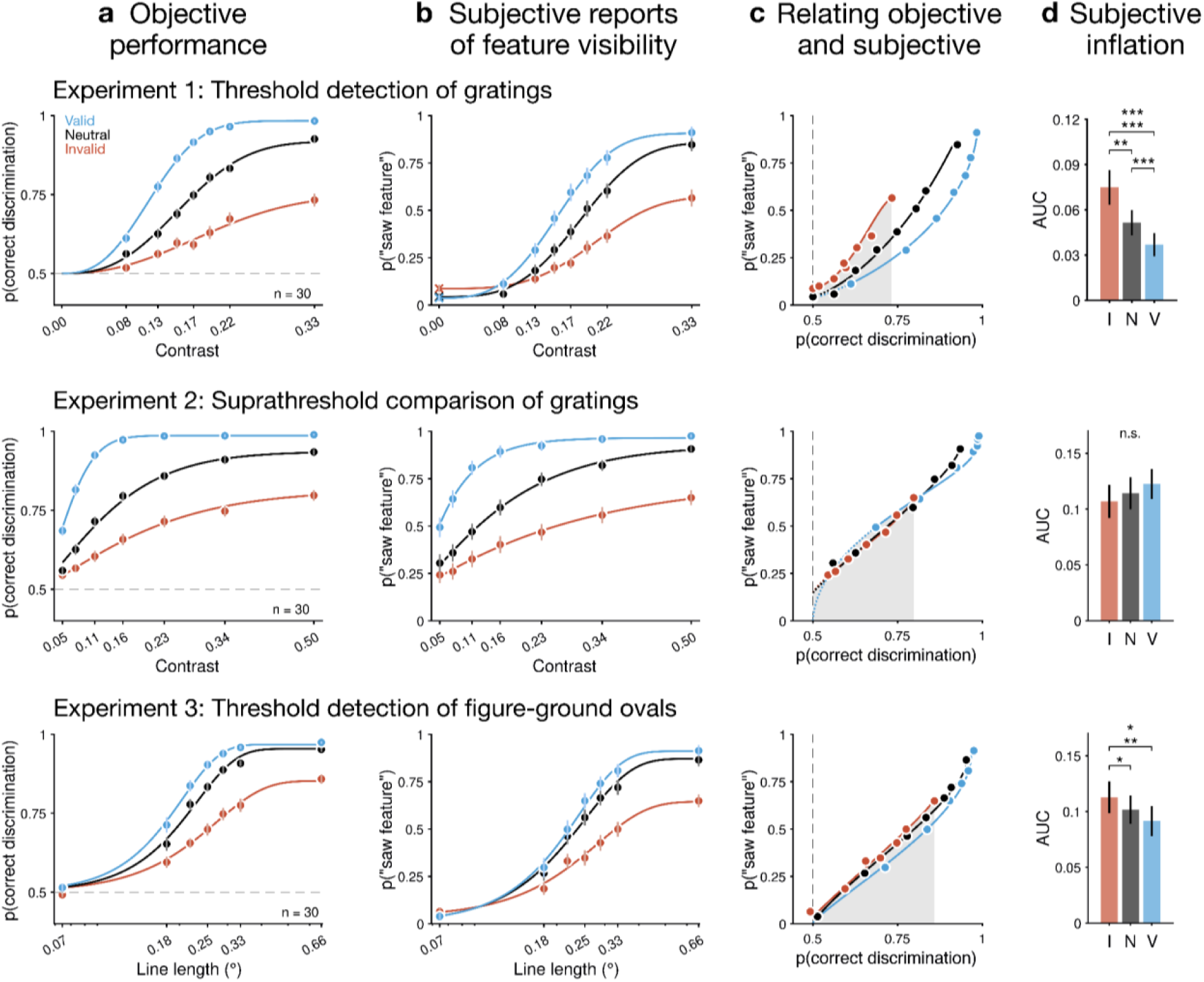
Inattentional inflation of task-relevant feature visibility. In three experiments (one per row), **a)** objective performance increased with stimulus strength and with attention (blue = valid, gray = neutral, red = invalid; same data as Figure 2a). **b)** Subjective reports of seeing the task-relevant feature (i.e., the orientation of the grating or oval) increased with stimulus strength. **c)** Subjective reports as a function of objective performance. For a matched range of performance (shaded gray region), subjective reports of the task-relevant feature were higher under inattention in Experiments 1 and 3. **d)** The AUC was greater for unattended than attended stimuli in threshold (repeated measures ANOVA per Experiment 1 and 3, all *p*<0.011) but not suprathreshold regimes (Experiment 2, non-significant trend in the opposite direction, *p*=0.118), so inattention inflated the task-relevant feature but only in threshold regimes. Group means are fit with Weibull functions (a,b) or their corresponding relative psychometric function (c) for visualization. The portion of the fit spanning the range of performance observed between the lowest and highest stimulus strengths is shown as a solid line; the portion extrapolated to chance performance is dotted. \**p*<0.05, \*\**p*<0.01, \*\*\**p*<0.001. Data (total n=90; n=30 per experiment) are presented as mean values ±1 SEM.

Whereas the contrast dependence of grating discriminability^3,5,53^ and visibility^10–14^ is well established, the effect of texture line length on the perception of texture-defined figures has not been tested without accompanying luminance confounds^54^. Here we ensured all textures had equal luminance (see Methods, Luminance calibration). Therefore, we first sought to confirm the effects of our stimulus manipulations on objective and subjective reports. We found that all objective **(Figure 2a)** and subjective measures **(Figure 2b, 3b)** increased with stimulus strength for both grating and texture targets (all *p*<0.001, full ANOVA tables are presented as **Supplementary Tables 1-3**) and spanned a wide range of performance and visibility levels, confirming the efficacy of our stimulus manipulations. Planned pairwise comparisons revealed each successive increment in stimulus strength significantly increased objective and subjective reports across all experiments (all *p*<0.004 after Holm’s correction). Participants reported seeing the task-relevant feature significantly less often than they reported seeing the stimulus (Experiment 1 gratings: (*F*(1,28)=138.75, *p*<0.001, *η*^2^=0.39; mean difference=26% [25,28]); Experiment 3 textures: (*F*(1,28)=108.61, *p*<0.001, *η*^2^=0.39; mean difference=26% [24,28]). Thus, subjectively perceiving a stimulus did not guarantee subjectively perceiving its features, even for easily discriminable features (e.g., +45° vs. -45° tilted gratings).

### Attention improved objective performance

To characterize the impact of attention on the full psychometric function, we fit Weibull functions to each participant’s objective and subjective reports, separately, for each cue validity condition (see Methods, Psychometric function fits). Across all experiments, attention improved orientation discrimination performance (main effect of validity: *F*(2,222)=892.80, *p*<0.001, *η*^2^=0.52, ε=0.70, **Supplementary Table 1**), consistent with previous reports^3,12,20^. Averaged across stimulus strengths, performance was highest on valid (87% [84,90]), intermediate on neutral (77% [74,79]), and lowest on invalid (66% [63,68]) trials. Reaction times were fastest for valid (0.55 s [0.53,0.58]), intermediate for neutral (0.87 s [0.84,0.90]), and slowest for invalid (0.97 s [0.94,0.99]) trials (*F*(6,666)=14.56, *p*<0.001, *η*^2^=0.01, ε=0.35), ruling out speed-accuracy tradeoffs as driving performance improvements^55^ **(Supplementary Figure 2)**. The benefit of attention on performance was greater at higher stimulus strengths, as revealed by an interaction of stimulus strength and validity (*F*(12,1332)=53.93, *p*<0.001, *η*^2^=0.08, ε=0.75), consistent with multiplicative response gain^3^. These across-experiment effects of attention were also significant for each experiment individually. In the two detection experiments, attention improved performance even for stimuli reported as unseen^56^ **(Supplementary Figure 3)** (main effect of validity: *F*(2,112)=5.48, *p*=0.006, *η*^2^=0.03, ε=0.94, **Supplementary Table 4**), though we cannot be sure participants had no conscious experience of the stimulus for all such trials^57,58^.

### Attention increased the sensitivity of subjective reports to stimulus strength

Attention overall increased subjective reports of stimulus visibility **(Figure 2b)** (main effect of validity: *F*(2,222)=82.33, *p*<0.001, *η*^2^=0.05, ε=0.70), which was driven by Experiments 1-3. In Experiment 4, attention primarily increased the sensitivity of subjective reports to line length. Attention also increased reports of seeing the task-relevant feature **(Figure 3b)** (*F*(2,168)=327.53, *p*<0.001, *η*^2^=0.27, ε=0.63), with significant effects in each experiment that included this measure (all *p*<0.001, **Supplementary Tables 2-3**). For both subjective measures, attentional enhancements were more pronounced at higher stimulus strengths (interaction of stimulus strength and validity; stimulus visibility: *F*(12,1332)<67.52, *p*<0.001, *η*^2^=0.05, ε=0.53; task-relevant feature visibility: *F*(12,1008)=71.71, *p*<0.001, *η*^2^=0.06, ε=0.48).

In fact, for weak or absent stimuli, inattention tended to elicit stronger subjective reports **(Figure 2b)**. In the threshold detection tasks, inattention increased false alarms^21,22^: the probability of reporting seeing a stimulus when none was present. This pattern was observed for both grating **(Figure 2b**, row 1) (*F*(2,56)=21.88, *p*<0.001, *η*^2^=0.07, ε=0.58) and texture stimuli **(Supplementary Figure 4)** (*F*(2,56)=9.54, *p*<0.001, *η*^2^=0.03, ε=0.61). In the suprathreshold comparison tasks, the probability that the test stimulus was reported as appearing stronger than the reference, in cases when the test was in reality weaker, was again higher under inattention, for both grating **(Figure 2b**, row 2) (*F*(2,56)=11.67, *p*=0.001, *η*^2^=0.04, ε=0.67) and texture stimuli **(Figure 2b**, row 4) (*F*(2,54)=30.43, *p*<0.001, *η*^2^=0.11, ε=0.75). Thus, attention increased the sensitivity of subjective reports of stimulus visibility to the true physical strength of the stimulus, resulting in relatively lower reports for absent or weaker stimuli and higher reports for stronger stimuli when they were attended vs. unattended.

The strength-dependent effects of attention on subjective measures may seem at odds with reports of attention increasing perceived strength across the entire psychometric function^11,13,14,59^; however, methodological differences can reconcile the two patterns. In studies that report wholesale increases in subjective measures with attention, participants judged which of two simultaneous stimuli appeared stronger, so that enhancement of the cued stimulus and suppression of the non-cued stimulus together contributed to the visibility judgment. Using this approach, a stimulus of a given physical strength was more often deemed stronger when cued than non-cued. In our experiments, the visibility judgment involved comparing a single stimulus to an internal criterion for detecting a stimulus (Experiments 1 and 3) or an internal criterion of the reference strength (Experiments 2 and 4).

### Apparent effects of attention on criterion setting may be better characterized as sensitivity effects

In a signal detection theory (SDT) framework^60,61^, attention consistently improves detection sensitivity^20,21,62^ (*d’*), the ability to determine whether a target is present vs. absent. Attentional effects on criterion (*c*), the propensity to report seeing a target, have been mixed^20,21,44^. Therefore we used SDT to assess sensitivity and criterion in the two detection experiments (Experiments 1 and 3).

In Experiment 1, there is one set of target-absent trials (when the grating contrast = 0) and multiple sets of target-present trials (when contrasts > 0). Because there is a single noise distribution, it is not possible for participants to set more than one criterion per attention condition. Conversely, in Experiment 3, each line length has its own set of target-absent and target-present trials, so separate criteria can, in principle, be set for each line length. Here, to facilitate comparisons across experiments and to previous literature, we chose to analyze all data by estimating different criteria *c*_YN_ for each stimulus strength level (where *c*_YN_ specifies the yes-no criterion, commonly referred to simply as *c*, estimated per stimulus strength. See **Supplementary Note 2** for a complete discussion of the modeling and interpretation of criteria in these different experimental designs).

Although attention has previously been found to yield higher *c*_YN_ values in near-threshold tasks by Rahnev et al.^21^, their computational model predicts that this effect should reverse at high enough stimulus strengths (**Supplementary Note 2**). Our findings confirm this prediction **(Figure 4a, Supplementary Figure 5b)** (interaction of validity and stimulus strength: *F*(12,672)=61.69, *p*<0.001, *η*^2^=0.04, ε=0.61). Attention increased *c*_YN_ at weaker stimulus strengths but its effect gradually reversed, such that attention decreased *c*_YN_ at higher stimulus strengths. At matched detection sensitivity, the effect of attention was consistently to increase *c*_YN_ **(Supplementary Figure 5c)**.

**Figure 4.**
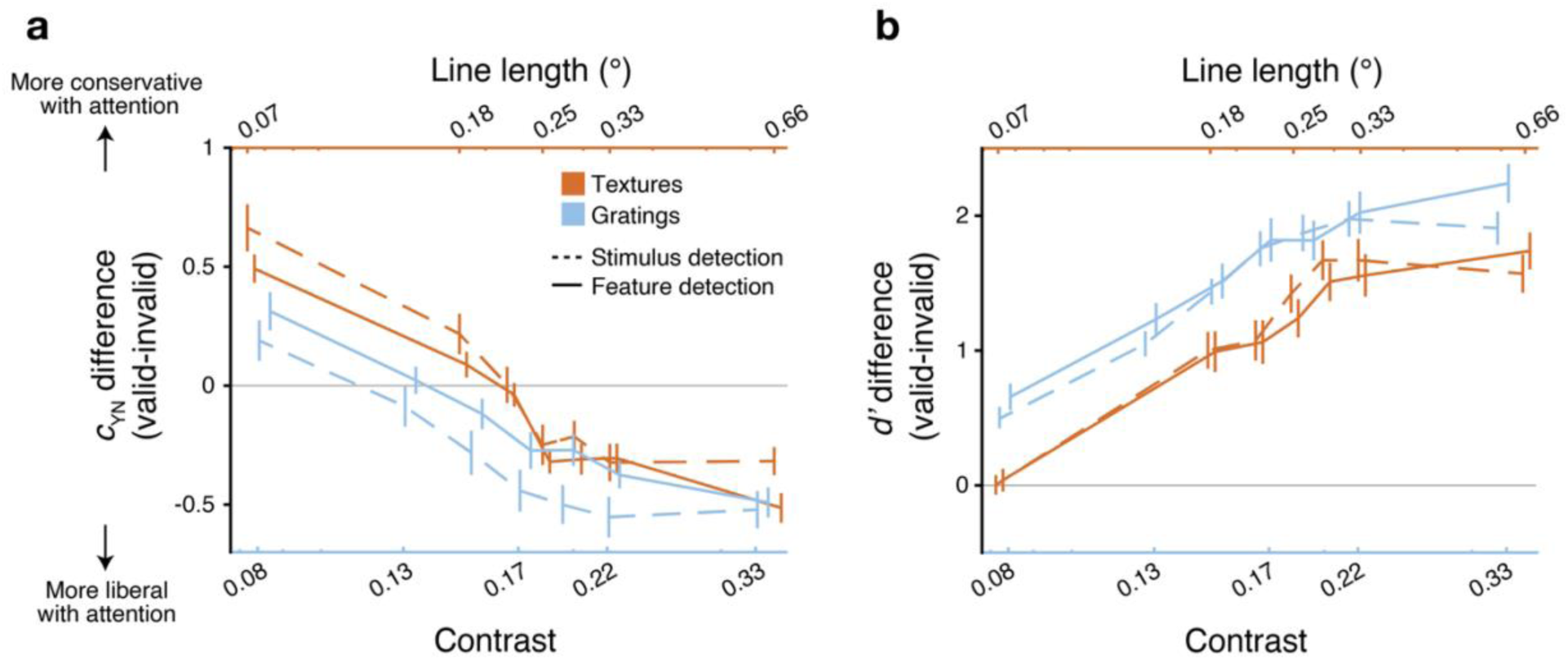
Attentional modulation of signal detection theory measures. **a)** Here we show the difference in the yes-no criterion *c*_YN_ between valid and invalid attention trials for detecting gratings (Experiment 1, blue) or texture-defined ovals (Experiment 3, orange) as a function of stimulus strength, which was controlled by either grating contrast (bottom *x*-axis) or texture line length (top *x*-axis). Attention made *c*_YN_ more conservative (positive difference) at lower stimulus strengths, which reversed (negative difference) at higher stimulus strengths (but see **Supplementary Note 2** for why these patterns may be best understood in terms of differences in sensitivity rather than criterion setting; in connection with this, note how the *c*_YN_ patterns mirror those of *d’* in panel b). **b)** Across both detection experiments, attention improved detection sensitivity (*d’*) and more so at stronger stimulus strengths. The effects of attention on SDT measures were consistent for detecting a stimulus as a whole (dashed lines) and for detecting a particular task-relevant stimulus feature, the grating or oval orientation (solid lines). Data (total n=60; Experiments 1 and 3 each n=30) are presented as mean values ±1 SEM.

Meanwhile, attention increased detection sensitivity across the entire psychometric function (main effect of validity: *F*(2,112)=296.54; *p*<0.001, *η*^2^=0.40, ε=0.74), and more so at higher stimulus strengths (interaction of stimulus strength and validity: *F*(12,672)=56.13, *p*<0.001, *η*^2^=0.10, ε=0.77) (**Figure 4b, Supplementary Figure 5a**). As detailed in **Supplementary Note 2**, due to Experiment 1’s task structure, there can only be one criterion per attention condition, and so *c*_YN_ values computed for each stimulus strength must reflect changes in sensitivity rather than criterion setting. A similar phenomenon likely occurs in Experiment 3, since the empirical relationship between *c*_YN_ and *d’* in that experiment closely mirrors the arithmetic relationship of these quantities in Experiment 1 (**Supplementary Figure 5c**). The measurement here of full psychometric functions thus suggests that the apparent effects of attention on criterion setting previously reported^21,22,44^ may be attributable to the effects of attention on sensitivity.

Across both detection experiments, the effect of attention on SDT measures for detecting the task-relevant feature behaved the same as its effect on detecting the stimulus as a whole: attention increased detection sensitivity across the entire psychometric function **(Figure 4b, Supplementary Figure 5a)** (main effect of validity: *F*(6,336)=397.97, *p*<0.001, *η*^2^=0.64, ε=0.72) and more so at higher stimulus strengths (interaction of validity and stimulus strength: *F*(12,672)=47.89, *p*<0.001, *η*^2^=0.10, ε=0.74), while making *c*_YN_ more conservative or liberal, depending on stimulus strength **(Figure 4a, Supplementary Figure 5b)** (interaction of validity and stimulus strength: *F*(12,672)=58.27, *p*<0.001, *η*^2^=0.04, ε=0.66). The effects of attention on the SDT measures were significant for each detection experiment individually and for both subjective reports types (all *p*<0.001, **Supplementary Tables 5-8**).

To confirm that these effects of attention on SDT measures did not depend on any unmet assumptions of equal variance of the internal noise and signal distributions, we also calculated the unequal variance measures: *d_a_* and *c_a_* (see Methods, Signal detection theory). The effects of attention on these measures were consistent with those found for *d’* and *c*_YN_ (**Supplementary Figure 6, Supplementary Tables 9-12**).

### Inattentional inflation of stimulus visibility

After characterizing the effects of attention on objective and subjective measures separately, we turned to our main question: under what conditions, if any, does inattention lead to subjective inflation? To test for inattentional inflation, we determined whether subjective reports were higher for unattended stimuli when matched in performance to attended ones. For each attention condition, we plotted subjective reports as a function not of stimulus intensity but of discrimination performance. Leveraging the psychometric functions fitted separately to objective and subjective measures, we then constructed a “relative psychometric function” to describe their relation^46,49^ (Methods, Relative psychometric function) **(Figure 2c)**.

Across all attention conditions, subjective reports increased monotonically and in most cases nonlinearly with increasing performance (**Figures 2c, 3c**), as has been noted in other studies collecting joint measures^46,57,58,63^. So although increases in performance with stimulus strength were accompanied by increases in visibility, the two changed at different rates as stimulus strength increased across its full range.

Attention changed the relation between objective and subjective reports, as shown by the divergence of the relative psychometric functions across validity conditions in most cases. To quantify the effect of attention on the relative psychometric function, we calculated for each validity condition the area under the curve for the subjective measure (on the *y*-axis) across a range of performance common to all conditions (on the x-axis) (“AUC,” see Methods, Area under the relative psychometric function). A higher AUC indicates subjective reports in one condition were “inflated” over another, when the range of performance across conditions was matched (shaded gray regions in **Figures 2c, 3c**).

Across all experiments, the AUC for stimulus visibility **(Figure 2d)** was strongly and significantly modulated by attention: largest for invalid (0.13 [0.11,0.14]), intermediate for neutral (0.10 [0.08,0.11]), and smallest for valid (0.08 [0.06,0.09]) conditions (*F*(2,222)=117.24, *p*<0.001, *η*^2^=0.14, ε=0.91), revealing robust inattentional inflation across the full relative psychometric function. Inattentional inflation was significant for each experiment individually (all *p*<0.001, **Supplementary Table 13**) and found for most participants **(Supplementary Figure 7)**.

### Inattentional inflation of task-relevant feature visibility

To evaluate the possibility that subjective inflation arises from objective and subjective reports accessing different stimulus information, we asked participants to report whether or not they saw the feature relevant for the objective task (the orientation of the grating or oval) in three experiments (1-3). For near-threshold coarse discrimination tasks (as in Experiments 1 and 3), it is usually assumed that discriminating between very different stimulus features (e.g., +45° vs. -45° tilted gratings) relies on the same information as detecting the stimulus as a whole^64^. Perhaps as a result, studies of subjective inflation have never separately assessed inflation for the stimulus (e.g., “Did you see the stimulus?”) vs. the task-relevant feature (e.g., “Did you see the stimulus orientation?”).

Across all experiments, the AUC for feature visibility was significantly modulated by attention **(Figure 3d)**: largest for invalid (0.10 [0.08,0.11]), intermediate for neutral (0.09 [0.08,0.10]), and smallest for valid (0.08 [0.07,0.10]) conditions (*F*(2,168)=6.54, *p*=0.004, *η*^2^<0.01, ε=0.78, **Supplementary Table 14**). This pattern indicates that even when subjective reports stipulated the task-relevant feature, the relationship to feature discriminability could nonetheless decouple with attention.

Following a significant effect of experiment (*F*(2,84)=7.26, *p*<0.001, *η*^2^=0.13) and an interaction of experiment and validity (*F*(4,168)=7.67, *p*<0.001, *η*^2^=0.02, ε=0.78) on the “feature-visibility” AUC, we tested inflation of the task-relevant feature within each experiment. Inattention significantly inflated the task-relevant feature in threshold regimes, for both grating (Experiment 1: *F*(2,56)=14.63, *p*<0.001, *η*^2^=0.10, ε=0.62) and texture (Experiment 3: *F*(2,56)=5.27, *p*=0.011, *η*^2^=0.02, ε=0.89) stimuli (**Figure 3d**, top and bottom rows). But we found no evidence for inflation of the task-relevant feature in suprathreshold regimes for grating stimuli (Experiment 2: *F*(2,56)=2.22, *p*=0.118, *η*^2^<0.01, ε=0.64); if anything the pattern went slightly in the opposite direction (**Figure 3d**, middle row). We could not assess feature-level inflation for suprathreshold textures as feature-specific subjective reports were not collected in Experiment 4. Thus, we only found evidence for inattentional inflation of the task-relevant feature in threshold regimes.

All effects of attention on the stimulus- and feature-level AUC were simultaneously replicated across two experimental sites (no significant interactions of site and validity, all *p*>0.071; **Supplementary Tables 13-14**) **(Figure 5)**.

**Figure 5.**
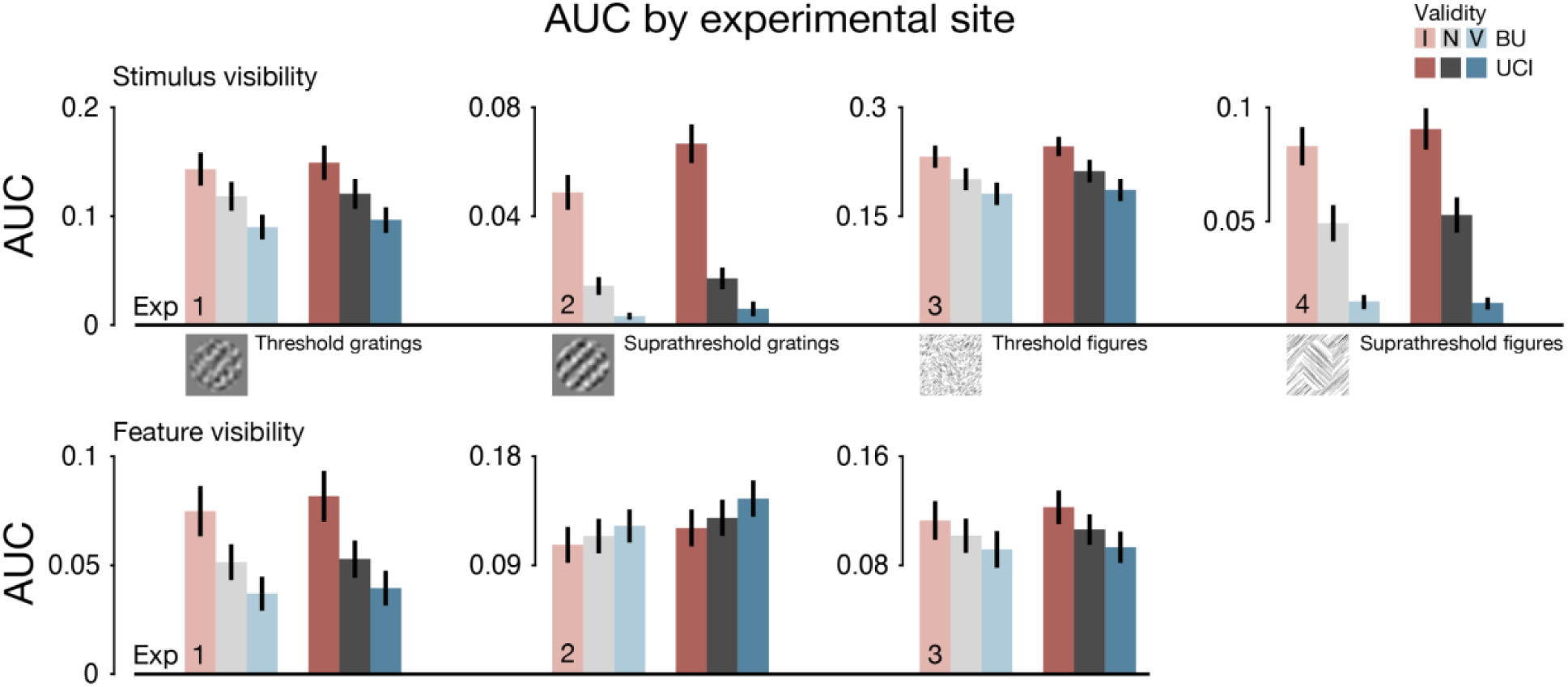
Inflation results replicated across experimental sites. The relative psychometric function AUC, quantifying subjective reports over a matched range of objective performance, at each experimental site (Boston University [BU] vs. University of California Irvine [UCI]). A larger AUC indicates one attention condition (blue = valid, gray = neutral, red = invalid) yielded “inflated” subjective reports over another, for a shared range of performance. Inattention inflated subjective reports of the overall stimulus (top) in all four experiments. Inattention inflated reports of the task-relevant feature (orientation, bottom) in threshold regimes (Experiments 1 and 3) but not suprathreshold regimes (Experiment 2). All effects of attention were simultaneously replicated across the two experimental sites (no significant interaction of site and validity for any experiment and visibility measure, all *p*>0.071). Data (n=15 per site and experiment, except Experiment 4 UCI n=13) are presented as mean values ±1 SEM.

### Comparing inattentional inflation across stimulus types, stimulus strength regimes, and visibility measures

To compare inattentional inflation across different stimulus types, stimulus strength regimes, and visibility measures, we calculated an attentional modulation index (AMI) as the AUC on valid trials subtracted from that on invalid trials, divided by their sum (see Methods, Attentional modulation index). Positive AMI values indicate inattentional inflation, negative values indicate inattentional deflation, and an AMI of zero indicates no dissociation of objective and subjective reports with attention. In all experiments, the AMI for stimulus visibility was significantly above zero **(Figure 6)** (*F*(1,110)=517.17, *p*<0.001, *η*^2^=0.83; mean AMI=0.51 [0.44, 0.58]), confirming strong and consistent inattentional inflation of the overall stimulus.

**Figure 6.**
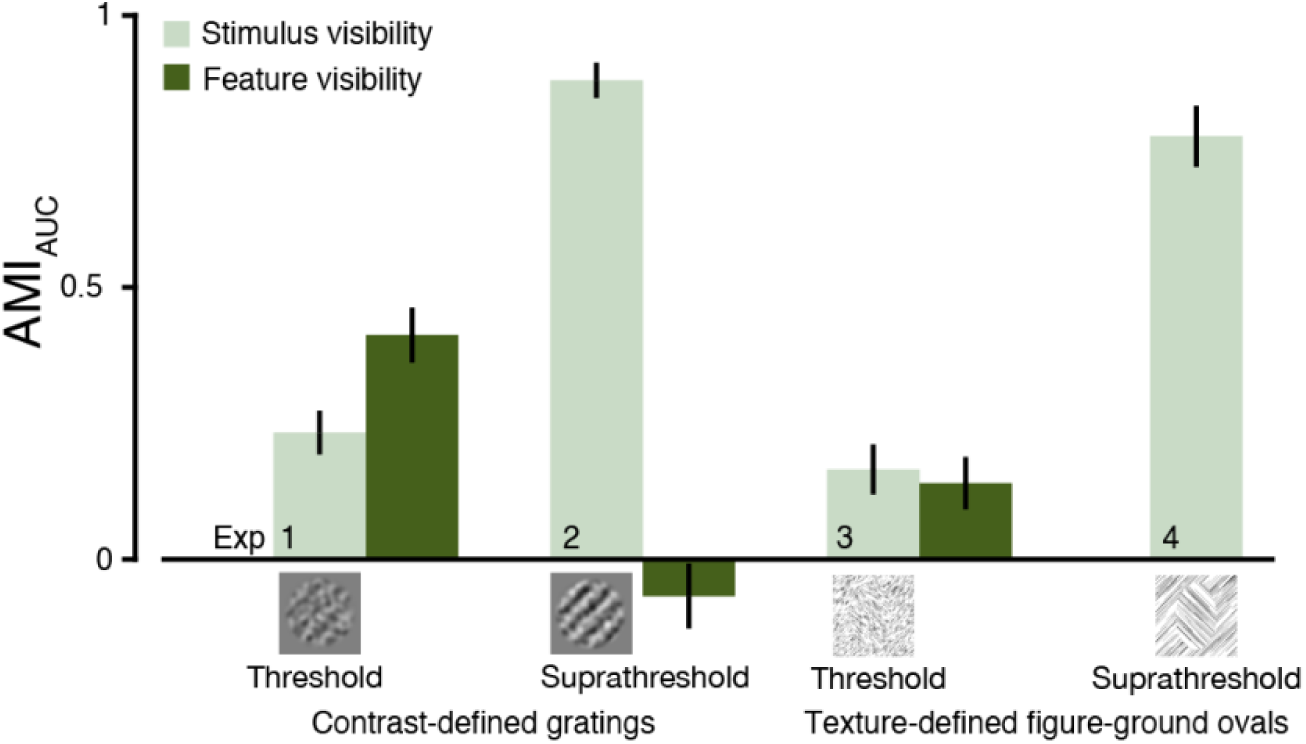
Comparing inattentional inflation across experiments. The AMI, quantifying the degree and direction with which objective and subjective reports dissociate with attention, was significantly greater than zero for stimulus visibility (light green) in all experiments (*p*<0.001), indicating robust and widespread inflation under inattention. Inflation of the overall stimulus visibility was more pronounced in suprathreshold (Experiments 2 and 4) than threshold-strength (Experiments 1 and 3) regimes (*p*<0.001). Inattention also inflated visibility of the task-relevant feature (dark green) but only in threshold regimes. Data (total n=118; Experiments 1-3 each n=30, Experiment 4 n=28) are presented as mean values ±1 SEM.

We first compared AMIs across strength regimes. Stimulus-level inattentional inflation not only occurred beyond threshold vision but was significantly more pronounced in suprathreshold regimes (main effect of strength regime: *F*(1,110)=197.65, *p*<0.001, *η*^2^=0.64; suprathreshold AMI 0.83 [0.77, 0.90] > threshold AMI 0.20 [0.14, 0.26], **Supplementary Table 15**). This pattern was similar for grating and figure-ground stimuli (no significant effect of stimulus type: *F*(1,110)=3.58, *p*=0.061, *η*^2^=0.03). Although the AMI was statistically larger for the suprathreshold experiments, this does not necessarily imply that attention had a larger effect on performance-matched visual experience, given that participants were asked to make qualitatively different kinds of subjective judgments about their visual experience in the tasks at threshold vs. suprathreshold.

We next compared AMIs across visibility measures. For near-threshold figure-ground stimuli (Experiment 3), the magnitude of inflation was similar for the overall stimulus and the task-relevant feature (no effect of visibility measure: *p*=0.707). But interestingly, for near-threshold grating stimuli (Experiment 1), inflation of feature visibility was stronger than inflation of stimulus visibility (main effect of visibility measure: *F*(1,56)=7.51, *p*=0.008, *η*^2^=0.12; AMI feature=0.41 [0.31, 0.51], AMI stimulus=0.23 [0.15, 0.31]).

All measures of AMI were simultaneously replicated at two experimental sites (no significant effect of site, all *p*>0.459, **Supplementary Table 15**). Moreover, to ensure that AMI analysis results were replicable, separate analysis pipelines that computed the AMI from raw data were developed independently at each site. We communicated across sites to align our analytic approaches conceptually but did not share code. There was no effect of analytic pipeline on AMI measures (all *p*>0.657, **Supplementary Table 16**) **(Supplementary Figure 8).** Therefore, the results were robust to differences in experimental setups and samples across sites, as well as to any idiosyncrasies in analytic choices across pipelines.

### Inattentional inflation occurs if and only if attention reduces performance thresholds more than visibility thresholds

Inflation suggests some asymmetry in how attention changes performance vs. visibility. To understand the nature of this asymmetry, we took advantage of having full psychometric functions to quantify attentional effects using a metric that generalizes across scales: psychophysical thresholds, which are expressed in units of stimulus strength. Attentional effects on thresholds are defined as the changes in stimulus strength required to preserve a certain level of performance or visibility as attention varies. Perceptual enhancement due to attention manifests as a reduction in thresholds, such that the same level of performance or visibility can be achieved at weaker stimulus strengths.

In **Supplementary Note 4**, we prove that under minimal assumptions, inattentional inflation in the relative psychometric function is equivalent to attention enhancing thresholds for performance more than thresholds for visibility in the corresponding conventional psychometric functions. Specifically, suppose that some unattended stimulus 𝑥_𝖴_ yields performance *P’* and visibility *V’*.

If attention reduces the stimulus strength needed to achieve *P’* more than it reduces the stimulus strength needed to achieve *V’*, then at the level of the relative psychometric function, visibility at *P’* is higher under inattention than under attention, i.e., there is inattentional inflation at *P’* **(Figure 8a,b)**. The converse is also true: inattentional inflation at *P’* implies that attention reduces the performance threshold more than the visibility threshold for the unattended stimulus 𝑥_𝖴_ that yields *P’*.

**Figure 8.**
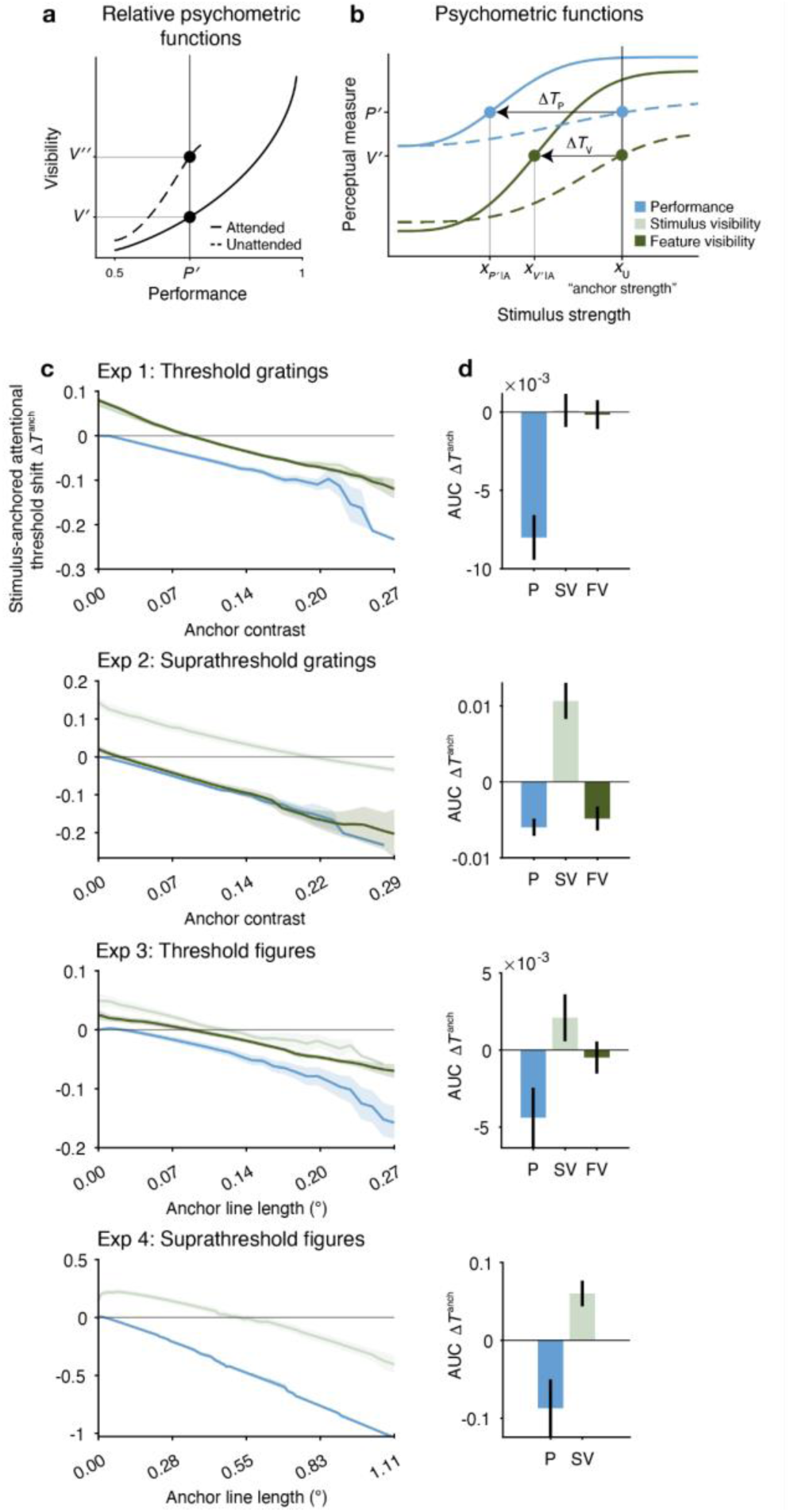
Inattentional inflation occurs if and only if attention reduces stimulus-anchored thresholds for performance more than for visibility. **a)** Example relative psychometric function showing subjective inflation: higher visibility when the stimulus is unattended 𝑉_𝖴_ vs. attended 𝑉_𝖠_ for matched performance level *P’*. **b)** The conventional psychometric functions for performance and visibility vs. stimulus strength yielding the relative psychometric functions shown in panel (a). When inattentional inflation occurs, attention reduces the thresholds at the corresponding unattended “anchor” stimulus 𝑥_𝖴_ more for performance (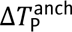) than for visibility (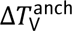) (see Supplementary Note 4). **c)** Stimulus-anchored threshold shifts due to attention for a full range of anchor strengths, Experiments 1-4. More negative shifts correspond to larger leftward shifts of the conventional psychometric function thresholds with attention. **d)** Area under the threshold shift vs. anchor strength curve (Δ𝑇^anch^ AUC), calculated for each perceptual measure from the data in panel (c). More negative AUCs correspond to larger leftward threshold shifts with attention, across all possible anchor strengths. Larger attentional reductions in performance thresholds compared to visibility thresholds correspond to inattentional inflation (cf. Figures 2-3). P = performance, SV = stimulus visibility, FV = feature visibility. Data are shown as means of individual estimates, and error bars indicate ±1 SEM.

To empirically demonstrate the threshold shifts corresponding to inattentional inflation effects (**Figures 2-3**), we used fitted psychometric functions to find the performance *P’* and visibility *V’* associated with a given “anchor stimulus” 𝑥_𝖴_, and measured the change in the thresholds for *P’* and *V’* due to attention for all values of 𝑥_𝖴_ in a usable range (see Methods, Attentional effects on performance and visibility thresholds). We refer to these threshold changes as “stimulus-anchored attentional threshold shifts” for performance and visibility, 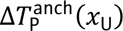 and 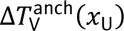, to emphasize that changes due to attention in both thresholds are yoked to a common unattended stimulus strength 𝑥_𝖴_.

For all perceptual measures, we found that the stimulus-anchored attentional threshold shift became more negative with increasing anchor strength **(Figure 8c)**, consistent with attention-induced gain increases in the underlying psychometric functions. Critically, attention reduced thresholds more for performance than for visibility 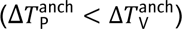 (**Figure 8c)**. This pattern generalized across the full range of anchor strengths, quantified as the area under the threshold shift vs. anchor strength curve (Δ𝑇^anch^ AUC; see Methods, Attentional effects on performance and visibility thresholds). In all experiments, the Δ𝑇^anch^ AUC was lower for performance than for visibility (all *F*>9.64, *p*<0.004; **Supplementary Tables 17-18**), with the exception of feature visibility in Experiment 2 (*F*(1,26)=2.68, *p*=0.114), which did not show inflation **(Figure 8d)**. These findings mirror the inattentional inflation effects observed in **Figures 2** and **3**. In sum, the prominent inattentional inflation effects in these data can also be understood as a general finding that attention enhances performance thresholds more than visibility thresholds.

## Discussion

In four experiments manipulating voluntary spatial attention, we tested claims that the unattended visual periphery can appear “subjectively inflated” relative to objective performance. We put the phenomenon to several key tests. Inattentional inflation was examined in threshold to suprathreshold regimes, using different stimulus types, and over wide ranges of performance. When the subjective report concerned the overall stimulus, inattentional inflation withstood all tests: covertly unattended items were reported as appearing stronger and more visible than attended ones, over matched ranges of performance. However, when the subjective report specified the visibility of the feature relevant for the objective task, we found that inattentional inflation only survived tests in threshold regimes. The results indicate attention regularly, though not invariably, dissociates objective and subjective aspects of perception, thus preserving inattentional inflation as a possible explanation of the apparent richness of the unattended periphery and as a motivating observation for higher-order theories of consciousness.

Using a high-powered experimental design and improved methodology, we overcame limitations of previous studies to provide strong evidence for both the existence and extent of inattentional inflation. While subjective inflation has been an influential concept in the field^24,25,27–30,33,39,65–75^ it has been based on relatively limited empirical evidence, even in a threshold regime^21,23,43^. Beyond threshold, tests of subjective inflation have not been reported, leaving it unknown whether inflation could affect the perception of clearly visible stimuli typical of normal viewing conditions.

We tested the robustness and generality of inattentional inflation in several ways. First, we used an analytic approach recently developed by our group^46^ to relate objective and subjective reports over full psychometric functions. This strategy allowed us to measure subjective reports over large matched ranges of performance across conditions, which was not possible using previous “performance matching” approaches^21,22^ that restricted tests of inflation to single performance levels. Second, we developed an approach to test inflation beyond threshold regimes, by having participants compare the subjective strength of a clearly visible target to a reference, while discriminating a target feature. Together, these methodological advances enabled tests of inattentional inflation across a full range of performance levels and stimulus strengths, revealing that inflation is strong, widespread, and replicable, though not without limits.

Our findings demonstrate that inattentional inflation generalizes to stimuli beyond those dependent on low-level visual properties, like contrast^21,23^ and color^22^ sensitivity. When the peripheral targets were texture-defined figure-ground stimuli, inattention inflated how strongly the figures appeared to “pop-out” from the background, showing that inflation can occur for stimuli defined by the mid-level visual property of texture segmentation. Inflation of these figures behaved similarly to that of simple gratings, indicating the phenomenon can operate at multiple levels of processing and may be commonplace in everyday vision. Moreover, figure-ground stimuli generate visual cortical recurrent processing^54,76–79^, a candidate first-order substrate of conscious vision^40,41^. Assessing subjective experience using these particular stimuli can thus help arbitrate and refine theories of conscious perception.

Stimulus-level inflation also generalized to suprathreshold regimes. Indeed, of the conditions we tested, subjective reports of suprathreshold stimulus visibility exhibited the largest magnitude of inflation. One author’s experience was that inflation effects are obvious at the single-trial level in the suprathreshold task, prompting an alternative framing of inflation that may inform future research (**Supplementary Note 3**). The striking suprathreshold inflation of perceived stimulus strength may contribute to the impression of a rich, intact visual world that extends across the visual field and beyond the focus of attention^28,29,33,66,80^. If the subjective strength of unattended items can far exceed what might be predicted from feature discriminability in suprathreshold scenarios, this discrepancy may help explain the sense of surprise in inattentional^81^ and change^82^ blindness demonstrations, in which people fail to notice a salient stimulus or stimulus alteration but feel certain they should have, given their subjective impressions. It may suggest that unless explicitly instructed, people make such metacognitive assessments based on overall stimulus visibility rather than the visibility of a particular stimulus feature.

If objective and subjective reports derive from different stimulus features^83,84^, their dissociation can be straightforwardly explained. For example, motion in the periphery could signal someone’s approach without indicating their identity, and such different information sources could be differentially affected by attention^53^. A strong test of inflation should therefore constrain subjective reports to the stimulus feature relevant for the objective task. But previous studies have not attempted to do so. When we asked participants to report the visibility of the feature discriminated in the objective task, subjective reports were still inflated under inattention in threshold regimes. However, in the suprathreshold task where these reports were available (Experiment 2 with grating stimuli), we did not find inflation of the task-relevant feature; instead, the relative psychometric functions were statistically indistinguishable across attention conditions. We do note that these were the only reports to be collected with a second keypress, so it is possible that post-decisional effects or extra cognitive load uniquely affected these subjective judgments in a way that obscured potential inflation effects. Thus, while we found that inattention does not inflate feature visibility in some suprathreshold scenarios, feature-level inflation was robust and consistent near the detection threshold.

A prominent model of inattentional inflation at threshold is based on signal detection theory^21^. According to this model, separable components of the internal signal govern subjective reports and objective performance, allowing their dissociation. A stimulus is reported to be visible when its internal signal magnitude exceeds a threshold, whereas its discriminability depends on the signal-to-noise ratio. Inattention is hypothesized to decrease the response magnitude but increase the variability of the internal signal^85^, which can increase the likelihood of crossing a fixed detection threshold. Studies have shown dissociable effects of neural variability^86,87^ on objective and subjective aspects of perception, lending empirical support to the model. Its dissociable computational mechanisms may map onto dissociable aspects of subjective perceptual experience (**Supplementary Note 3**).

This signal detection model^21^ can reproduce several key phenomena in our data regarding how perceptual metrics change for unattended stimuli relative to attended stimuli in detection tasks: false alarm rates increase; both subjective reports and objective performance become less sensitive to stimulus strength; and at matched performance, stimuli are reported as more visible. Additionally, the model can account for the yes-no criterion being more conservative under attention at low stimulus strengths, and it predicts that the criterion becomes more liberal under attention at high stimulus strengths. (See **Supplementary Note 2** for further elaboration of this model prediction, and for an argument that this yes-no criterion effect may be better understood not as an effect on criterion setting *per se*, but rather as an indirect reflection of more primary sensitivity effects.) Here, by measuring the full psychometric function, we not only confirm this prediction, but also provide a unifying explanation for apparent contradictions previously reported in the literature, wherein attention to peripheral stimuli has been variously found to correspond to conservative^21,22,26^, neutral^20^, or liberal^44^ shifts in the apparent detection criterion, depending on the task design and stimuli chosen. Although the model of Rahnev et al.^21^ qualitatively accords with several of our findings in the threshold regime, further work is required to extend this model and other contending models to determine their ability to capture feature-specific reports and suprathreshold regimes, and to quantitatively fit the entire dataset.

According to higher-order theories of conscious perception, subjective inflation arises from an overestimation of the strength of the sensory signals governing performance, driven by a putative implicit metacognitive mechanism^30^. By proposing two levels of representation, higher-order views can account for the decoupling between objective and subjective aspects of perception. But these views do not immediately explain why or under what circumstances they will come apart. In the case of attention, higher-order views do not in general explain why withdrawing attention should inflate subjective strength, when the first-order states are presumably comparable, yielding matched performance. That said, one proposal posits a higher-order Bayesian observer that misrepresents the distribution of sensory noise in the periphery or under inattention, leading to subjective inflation^67,73^.

Some proponents of higher-order theories have interpreted the fixed threshold of visibility in the signal detection theory model of Rahnev et al.^21^ as a higher-order mechanism, likely carried out in prefrontal areas^88^. In this view, the higher-order areas gate the entry into phenomenal consciousness, and the fixed criterion reflects a stable higher-order threshold for conscious perception^32,33^. However, the signal detection model is also compatible with some versions of a first-order view holding that the sensory signal is sufficient to generate conscious perception. The criterion could then be interpreted in different ways consistent with some versions of a first-order view^71^. For example, the criterion could reflect the sensory signals themselves exceeding some threshold that is not set by any higher-order mechanism. Inflation could then arise from attention changing the sensory signals, as in the Rahnev et al.^21^ model. Or, the criterion could be decisional rather than perceptual, governing whether the participant reports seeing the stimulus rather than whether they actually see it^69,89,90^, although the resistance of subjective inflation to feedback and reward gives some reason to think the phenomenon is perceptual^21,23^. Overall, while failure to find inattentional inflation would have challenged a motivation of higher-order theories—and the predictions made by higher-order theorists in the current adversarial collaboration—the behavioral demonstration of inflation is not on its own necessarily incompatible with first-order theories (**Supplementary Note 1**).

By characterizing inflation over full psychometric functions, we identified a simple principle: inattentional inflation occurs if and only if attention reduces performance thresholds more than visibility thresholds for a given stimulus **(Supplementary Note 4)**. We show empirically that this principle holds across all stimulus strengths and all experimental conditions that showed inflation. Because this principle emerges from the mathematical relation between relative and conventional psychometric functions, it is independent of any theory or model of inflation. Any model that can account for performance-matched inflation will also be able to account for the corresponding threshold shifts, and vice versa. Making this mathematical relation explicit may inform future models, whether they invoke first-order or higher-order mechanisms to produce inflation.

Importantly, given the clear data obtained here, any theory of subjective awareness must now take seriously the phenomenon of inattentional inflation. First-order theories must account for the widespread occurrence of inflation, while higher-order theories must account for the conditions under which it is absent. The current dataset, testing large numbers of experimental conditions with high trial counts, provides by far the most comprehensive extant data on subjective inflation, and theories of awareness should seek to explain the detailed pattern of these data. The measurement of full psychometric functions is likely to strongly constrain model fits, allowing model identification that would not be possible with single pairs of performance-matched stimuli. The current dataset, which we have documented and made publicly available to invite model fitting by the wider community, can serve as a benchmark dataset for the field.

Going forward, inattentional inflation presents an opportunity to identify the neural processes specific to subjective perception. Typically, the neural signals that track subjective strength tightly correlate with stimulus processing that supports objective performance, confounding their interpretation^45,91,92^. But by reliably dissociating objective and subjective aspects of perception, inattentional inflation may be a valuable approach to isolate the neural signals that uniquely covary with subjective strength while objective performance is controlled. Different theories of conscious perception make different predictions about which neural processes will correlate specifically with subjective strength (e.g., first-order processes in sensory areas^42,90^, higher-order processes in prefrontal areas^30^, global ignition^93^; for reviews^38,39,94^), making inattentional inflation a promising tool—along with other methods for decoupling objective and subjective aspects of perception^58,86,87,95–104^ to adjudicate among these theories.

The idea that our introspection can be but a dubious authority on our own visual performance has long been recognized. Clinical cases in which visual cortical damage leads to ignorance of residual capacity (blindsight)^105^ or incapacity (Anton’s syndrome)^106^ have shown powerful, sometimes permanent dissociations between subjective reports of visual awareness and the objective ability to discriminate visual features. Our findings show that even in healthy observers, following temporary, voluntary fluctuations of visuospatial attention, objective and subjective aspects of perception routinely come apart. What we think we can see therefore may not accurately reflect how well we can distinguish visual features, particularly at the threshold of vision.

## Methods

### Participants

One hundred twenty healthy adult humans (88 females and 32 males, ages 19-36, based on self-report) participated across four experiments, which was the preregistered target sample size. Fifteen participants per experiment participated at each of two research sites: Boston University (BU) and the University of California, Irvine (UCI). Each participant completed an average of ∼3400 trials (range of 2288-4771 trials) across four to six 1.5-hour-long visits on separate days, for both a highly powered sample size and reliable measurements for each participant. Authors AS, TK, JAM, EO, EER, MEW, and JW participated in the experiments. All other participants were naive to the study design. All participants provided informed consent, and the University Committee on Activities Involving Human Subjects at BU and the Institutional Review Board at UCI approved the experimental protocols. All participants had normal or corrected-to-normal vision and were monetarily compensated for their time. No sex or gender-based analyses were performed, and we did not consider sex or gender in the study design, as neither sex nor gender played a role in our research questions.

### Procedure

Participants were seated in a dark room 75 cm from a computer monitor. Stimuli were generated on Linux at BU and Windows at UCI using MATLAB and Psychophysics Toolbox^107–109^. Stimuli were displayed on a VIEWPixx LCD monitor (VPixx Technologies Inc., QC, Canada) with a resolution of 1920 x 1080 pixels and a refresh rate of 120 Hz at BU and a CRT monitor (NEC MultiSync FE2111SB) with a resolution of 1280 x 1024 pixels and a refresh rate of 60 Hz at UCI. To linearize the contrasts, the displays were calibrated using a Konica Minolta LS-100 Luminance Meter (Konica Minolta, Tokyo, Japan). Participants had their head position stabilized in a chin - and-head rest. Gaze position was continuously recorded at a sampling frequency of 1000 Hz using an EyeLink 1000 (SR Research Ltd., ON, Canada) at BU and 500 Hz using a LiveTrack Lightning eye-tracker (Cambridge Research Systems, Ltd., UK) at UCI.

### Task

In all four experiments, participants performed a spatial attentional cueing task while fixating on a central cross **(Figure 1, Supplementary Figure 1)**. The screen was divided into four quadrants. On each trial, participants viewed up to four peripheral targets, which varied independently across seven strengths. Only one quadrant was relevant for the report, as indicated by a post-stimulus response cue. Before the targets, one or all four arms of a black central precue, each 1° long by 0.1° wide, flashed white (50 ms) to direct covert attention focally to one quadrant (80% of trials) or in a distributed fashion across all quadrants (20% of trials; neutral attention condition). The focal precue matched the response cue with 75% validity, to incentivize using the spatial information provided by the precue, leading to 60% valid and 20% invalid attention trials overall. The precue appeared 300 ms before the targets, to allow for the deployment of covert attention to the cued location^6,12^. The targets were presented for 250 ms. A response cue 500 ms after target onset instructed participants to make both a forced-choice objective orientation report and a subjective visibility report about the response-cued quadrant **(Figure 1c)**.

The subjective report was either to indicate visibility of a near-threshold target (Experiments 1 and 3) or to compare the strength of a suprathreshold target to that of a learned reference (Experiments 2 and 4). In experiments 1-3, participants also reported whether or not they saw the feature relevant for the objective task: the stimulus orientation. All objective and subjective reports were made using a single keypress, except the feature visibility report in Experiment 2, which was made using a second keypress. For the subjective reports, participants were instructed to report how the stimuli appeared to them and not, for example, their confidence in the orientation judgment or what they considered the stimulus contingencies to be. See Supplementary Methods for task instruction excerpts.

When a response was registered, the fixation cross lightened to gray for 500 ms before the next trial was initiated. Otherwise, no feedback was provided. If no valid keypress was registered within 5 s following the response cue, the trial was aborted and not repeated. Participants rarely failed to make a valid keypress within the response window (on average 2.9 trials, SD=5.9). Between trials, a gray screen containing only the fixation cross appeared for 500 ms. The gray was a mid-gray (45.6 cd/m^2^ at BU, 60.7 cd/m^2^ at UCI) in the experiments with grating stimuli. In Experiment 3 with texture stimuli, 7 participants had the same mid-gray blank screen luminance. For the remaining participants in Experiments 3 and 4, the gray was lightened (78.7 cd/m^2^ at BU, 93.2 cd/m^2^ at UCI) to match the average texture luminance, with the goal of preventing eye-tracking issues due to luminance changes between trial stages.

Trials were grouped into consecutive runs of 560 trials. Within each run, precue validity and the location of the response-cued quadrant were counterbalanced. For each permutation of precue validity and response-cue location, stimulus strength and identity at the response-cued location were also counterbalanced. The presentation order of these counterbalanced conditions was pseudo-randomized within each run. On invalid trials, the location of the precued quadrant was randomly selected as one of the three locations not probed by the response cue. The stimulus properties at the non-response-cued quadrants were pseudo-counterbalanced, so that their marginal probabilities were controlled for within a run. Breaks were offered after every block of 112 trials, for 5 blocks per run. The attention manipulation and target timings were identical across experiments. However, the stimuli themselves and the nature of the task report differed from experiment to experiment.

### Stimuli

Visual targets were contrast-defined gratings embedded in noise (Experiments 1 and 2) or texture-defined figure-ground ovals (Experiments 3 and 4). Targets were present either half the time at threshold strengths (Experiments 1 and 3) or all the time at suprathreshold strengths (Experiments 2 and 4). The experiments are reported in a different order than they were collected; the order of data collection adhered to the order in the preregistration: 1) threshold detection of figure-ground ovals (Experiment 3), 2) suprathreshold comparison of figure-ground ovals (Experiment 4), 3) threshold detection of gratings (Experiment 1), and 4) suprathreshold comparison of gratings (Experiment 2).

#### Contrast-defined gratings

Gratings were luminance-modulated sinusoids with a spatial frequency of 1 cycle per degree. Gratings were centered at 5° eccentricity. In Experiment 1, the gratings were presented at low contrasts, calibrated per participant to be near threshold visibility when added to noise pedestals, and oriented ±45° from vertical. In Experiment 2, the gratings were presented at suprathreshold contrasts—fixed across participants from 5% to 50% in 7 log steps—and oriented about vertical at individual tilt thresholds (see Methods, Thresholding).

##### Noise pedestals

Gratings were added pixelwise to noise pedestals, then placed in a circular aperture 5° in diameter with a cosine edge subtending 0.5° that gradually faded to the background gray. To generate each noise patch, Gaussian noise was bandpass filtered around the grating spatial frequency ±1 octave. The filtered noise was then centered at mid-gray (matching the background luminance) and scaled to the desired contrast, which was 50% in the Experiment 1 and lowered to 20% in Experiment 2 to allow the superimposed gratings to have higher signal to noise ratio, thus increasing their visibility. Noise patches that by chance deviated from the desired mean contrast by more than 2% were regenerated.

#### Texture-defined figure-ground ovals

The texture “background” was made of parallel lines, oriented either ±45°, on which an oval “figure” delineated by orthogonal lines could appear (**Supplementary Figure 1**). Texture lines were black on a white background. The entire screen (21.6° by 38.4° at BU, 23.8° by 29.8° at UCI) was filled with textures, with the exception of a gray circle (2° radius) at the center of the screen containing the fixation cross and cues (see Methods, Online fixation monitoring). Each quadrant of the screen (5.4° by 9.6° at BU, 6.0° by 7.4° at UCI) was filled with a unique background texture, made of many randomly placed lines of uniform length and orientation. On a given trial, the background texture lines across quadrants were of the same orientation but could vary in length. Texture lines were truncated at the quadrant boundaries and the perimeter of the central circle so that no lines crossed these boundaries. The number of lines drawn scaled with line length, so that the proportion of black pixels (which controlled the mean luminance of the display) was fixed to 0.15 across all textures (see Methods, Luminance calibration).

The oval “figure” was centered at 8° eccentricity within a quadrant and oriented either vertically or horizontally along the major axis. Each oval was filled with a unique texture of many randomly placed lines, which were of the same length as the background texture lines but orthogonal in orientation. We ensured the ovals had buffers of at least 3.9° (at BU) or 1.7° (at UCI) between their perimeter and the edge of the screen, to minimize edge interactions from providing information about the oval’s orientation. The buffer values correspond to the smallest distance between the edge of a vertically oriented oval and the top or bottom of the screen. Texture lines of the background and figure were truncated at the oval perimeter to provide a clean figure-ground signal (see Figure 1b for examples).

In the threshold detection experiment (Experiment 3), when the oval figures appeared, they were clearly elongated (5° by 3° in aspect ratio) but made difficult to segment from the ground by presenting the texture lines at short lengths, calibrated per participant to be near texture segmentation thresholds. In the suprathreshold experiment (Experiment 4), the texture lines were longer in length—fixed across participants from 0.18° to 3° in 7 common log steps—so that the oval figure was more clearly segmented from the ground, but the oval aspect ratio was calibrated per participant (i.e., made more circular) so that the orientation was difficult to discern.

##### Random textures

Random textures consisted of scattered lines, each randomly oriented and drawn at any of the 7 possible line lengths. The mean luminance of the random textures (controlled by the proportion of black pixels) was fixed to 0.15, matching the luminance of the figure-ground textures (see Methods, Luminance calibration). Random textures were presented before the targets, starting from fixation (see **Supplementary Figure 1** for timing), with the intent of reducing target-evoked contrast transients in future neuroimaging versions of the study and to reduce pupil-dilation effects at target onset.

##### Luminance calibration

To control the mean luminance of the textures, we performed a calibration procedure prior to running the experiments. The calibration procedure was run separately on each stimulus presentation computer at each site to ensure consistency regardless of differences in display properties. The purpose of the calibration procedure was to determine, for a given line length, how many randomly placed texture lines should be drawn to a quadrant of the screen to ensure that the average proportion of black pixels in the quadrant achieved a target value. (It is not straightforward to derive an analytical solution for this problem, given that randomly placed lines may frequently overlap and be truncated at the quadrant boundaries.) The procedure ensured that all textures had equal luminance regardless of differences in line length and computer display properties. We chose a target proportion of 0.15 black pixels to achieve qualitative similarity of appearance to texture stimuli previously used in the literature.

The calibration procedure proceeded as follows. For a given line length 𝐿 and number of lines 𝑁, many background textures were drawn. For each texture generated in this way, the proportion 𝑃 of black pixels drawn to the quadrant was computed. The expected value of the proportion of black pixels <u>𝑃</u> was then computed as the average of 𝑃 across texture instances. For a given value of 𝐿, the procedure was repeated for many values of 𝑁, generating samples of a function <u>𝑃</u> = 𝐹(𝑁|𝐿) describing how the expected proportion of black pixels <u>𝑃</u> depends on number of lines drawn 𝑁 for a given value of line length 𝐿. Interpolation of these samples was used to find the value 𝑁_𝑇_ achieving the desired target value of <u>𝑃</u> = 0.15 for the specified value of 𝐿.

This procedure was then repeated for many values of 𝐿, generating samples of a function 𝑁_𝑇_ = 𝐺(𝐿) describing how 𝑁_𝑇_, the number of lines drawn to achieve <u>𝑃</u> = 0.15, depends on 𝐿. Interpolation of these samples was used to find the value 𝑁_𝑇_ achieving the desired target value of <u>𝑃</u> = 0.15 for any arbitrary value of 𝐿. This second interpolation was necessary to determine in real time how many lines to draw to a quadrant for the many possible values of 𝐿 that could be probed during the thresholding procedure of Experiment 3.

Since the structure of the random textures differed considerably from that of the oriented background textures, a similar calibration procedure was separately performed for the random textures to ensure proper luminance calibration at <u>𝑃</u> = 0.15. The line lengths of random textures were fixed values determined by the thresholding results for each participant, so this procedure only needed to interpolate over samples of a function <u>𝑃</u> = 𝐻(𝑁) and therefore could be performed quickly following the thresholding block to properly calibrate random textures for each participant before the main experiment.

### Thresholding

To calibrate stimulus properties so that the objective task was appropriately challenging, each participant completed a thresholding procedure prior to the main experiment. The thresholding task was identical to the main task, except a stimulus feature was continuously adjusted between trials using the QUEST Bayesian adaptive staircasing procedure^110^ and all precues were neutral. The adjusted feature in the detection experiments was stimulus strength (i.e., grating contrast or texture line length) and in the suprathreshold experiments was the grating tilt or oval aspect ratio. Three independent thresholding tracks, each adaptively approaching a discrimination accuracy of 75%, were interleaved for a total of 240 trials (80 per track) contributing to threshold estimates. On each trial, one of the three tracks was selected pseudo-randomly to set the strength of the response-cued stimulus to the current threshold estimate of that track, and subsequently that track had its threshold estimate updated using the participant’s accuracy on that trial. If the individual tracks did not qualitatively converge, participants repeated the thresholding procedure. There was no statistically significant difference in the median thresholded values by experimental site (factors of experiment and site, no effects of site, all *p*>0.144, **Supplementary Table 17**), suggesting the two experimental setups and samples were reasonably similar.

During the thresholding procedure for the detection experiments (Experiments 1 and 3), while target-absent trials were included to mimic the structure of the main task, only target-present trials contributed to QUEST estimates. Likewise, although participants made a subjective visibility report simultaneously with their orientation report, as in the main experiment, only the orientation report contributed to threshold estimates. Using the thresholding track with the median threshold estimate, 5 middle stimulus strengths were selected to yield orientation discrimination accuracies of 60%, 67.5%, 75%, 82.5%, and 90%. The lowest and highest stimulus strengths were selected to yield near-floor and -ceiling performance. To select comparable values at each extreme, the lowest stimulus strength was chosen to be midway between the minimum possible strength (i.e., 0% contrast or a line length of 3 pixels) and the strength yielding near-chance (51%) accuracy on a common logarithm scale. The highest stimulus strength was chosen to be the line length yielding 90% accuracy plus 𝑑, where 𝑑 was defined as the distance on a common log scale between the lowest stimulus strength and the strength yielding 60% accuracy. However, if in Experiment 1 this procedure resulted in a maximum grating contrast that exceeded 1 when added to noise, the thresholding procedure was repeated. In Experiment 3, if the procedure resulted in a maximum line length that exceeded the length of the oval minor axis (3°), the maximum line length was instead set to 3°.

During the thresholding procedure for the suprathreshold experiments (Experiments 2 and 4), stimuli were fixed to the middle (16% contrast or 0.41° line length) of the 7 possible target strengths (5% to 50% contrast or 0.18° to 3° line length in log steps). Because the targets were always presented at a fixed, easily visible stimulus strength, no subjective report was required. Instead, the task was made challenging by adjusting the grating orientation or oval aspect ratio to yield discrimination accuracy of 75%. The mean thresholded grating tilt was 8.9° (SD=6.6°). The mean thresholded oval aspect ratio as a proportion of the maximum aspect ratio was 0.67 (SD=0.31), with 1 corresponding to a maximally elongated oval (5:3° aspect ratio) and 0 a perfect circle. While the aspect ratio was adjusted, the area of the oval was held constant.

### Online fixation monitoring

To ensure that effects of attention could not be attributed to saccades towards the precued location, participants were instructed to maintain fixation on a small black cross (0.35° wide) displayed within a gray aperture (2° radius) in the center of the screen. Before each session, a calibration sequence converted raw gaze position into degrees of visual angle. The start of every trial was contingent upon fixation for 500 ms within a 2° allowance. If fixation was not acquired within 10 s, participants were shown a message to keep their gaze locked on the center of the screen. If fixation was unable to be acquired within this time limit on two consecutive trials, the calibration sequence was repeated. After acquisition, fixation was enforced until the onset of the response cue. If fixation was lost before then, due either to a saccade or a blink, the trial was stopped and repeated at the end of the block. Any trial interrupted by a fixation break was not completed and therefore not part of the behavioral data used for analysis. On average, participants broke fixation on about 5% of trials.

### Task training

To learn the task, participants first completed a self-guided instructions walk-through, followed by practice trials. Participants completed a sequence of practice trials with increasing difficulty: 1) slowed-down trials, 2) full-speed trials with auditory feedback, and 3) full-speed trials with no feedback. 1) The slowed-down trials were identical to trials in the main task, except stimulus timings were slowed down by a factor of 4. Participants had unlimited time to respond and received trial-by-trial feedback about both the objective and subjective reports in text to check their target and response mappings (e.g., “At the cued location, the stimulus was: counterclockwise (−45°). Your response was: counterclockwise (−45°), saw a grating but not its orientation”). 2) On trials with auditory feedback, the pitch of a 250 ms tone indicated orientation discrimination accuracy (correct=784 Hz, G5; incorrect=523 Hz, C5). Tones were enveloped by cosine ramps 10 ms wide. When the response-cued target was absent, the orientation judgment was irrelevant, so the auditory feedback played a high tone (G5) with 75% chance and a low tone (C5) otherwise, matching the expected average accuracy after thresholding. 3) The full-speed trials with no feedback mimicked trials in the main task.

For all practice stages, cueing validity, the response-cued quadrant, and stimulus properties were randomly chosen on each trial. Each practice block was 10 trials. The experimenter observed the participant during the practice trials to verify comprehension of the task and to answer any questions between blocks. Participants were allowed to repeat any of these blocks of practice trials until they felt comfortable with the task.

### Reference training

In the suprathreshold experiments (Experiments 2 and 4), the subjective report was to judge the visibility of the target relative to a reference. To learn the reference strength, participants completed 120 trials identical to the main task, except all precues were neutral and the response-cued target was fixed to a “reference” strength, which was set to the middle value (i.e., 0.17 grating contrast or 0.41° line length) of the 7 pre-defined stimulus strengths in the main task (see Methods, Stimuli). Rather than make a subjective strength report, participants were instructed to notice and internalize the reference strength. Participants then completed 112 trials incorporating the attentional precue, individually thresholded grating tilt or oval aspect ratio, and subjective report comparing the visibility of the target to the newly learned reference, mimicking the main task.

### Retention and exclusion

Datasets from participants who completed fewer than 2240 trials (i.e., 4 runs) were considered incomplete and not included in the analyses reported here. Across the 4 experiments at both sites, 21 participants with partial datasets were excluded. All partial datasets were due to participant dropout. We recruited additional participants until we reached the preregistered target sample size with complete datasets. To encourage retention, a monetary completion bonus was introduced during the third experiment at BU and the fourth experiment at UCI.

We excluded trials from analysis if no valid keypress was made before the response window timed out. One participant’s data from Experiment 4 was excluded from all analyses due to chance performance on the main task despite sensible behavior during the thresholding procedure. Another participant’s data from Experiment 4 was excluded from the AMI analyses because of chance performance on invalid trials regardless of stimulus strength, resulting in AUCs of 0 across all attention conditions, for which the AMI could not be computed. Although AUCs of 0 across attention conditions do not meaningfully contribute to the AUC analysis, the effect of attention on AUC was consistent whether or not this subject was included, so we included their data in the AUC analysis presented here.

### Data analysis

#### Psychometric function fits

For each participant, we fit psychometric functions (PFs) 𝜓 with parameters 𝜃 for objective task performance *P* and subjective reports of visibility *V* as a function of stimulus strength *x,* separately for attended (A), neutral (N), and unattended (U) conditions:

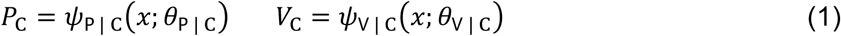

where *C* is a variable for attentional condition that can take on values A, N, or U. Thus e.g., for *C* = A, we have that 𝑃_𝖠_ is task performance under attention, 𝜓_𝖯 | 𝖠_ is the PF for task performance under attention, and 𝜃_𝖯 | 𝖠_ is the set of parameters for 𝜓_𝖯 | 𝖠_. Task performance pertains to the probability of correct orientation discrimination responses. Stimulus visibility pertains to the probability of reporting that the stimulus was seen in the threshold experiments (p(“saw stimulus”)) or the probability of reporting that the stimulus was stronger than the reference in the suprathreshold experiments (p(“test stronger”)). Task-relevant feature visibility pertains to the probability of reporting that the task-relevant stimulus feature was seen (p(“saw feature”)). We fit separate PFs for performance, stimulus visibility, and task-relevant feature visibility under each attentional condition in all experiments except Experiment 4, for which no data on task-relevant feature visibility was available.

We used the Weibull function for 𝜓, defined as

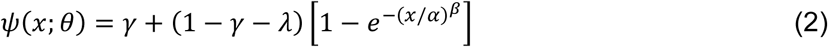

where 𝜃 is a set containing the parameters 𝛼 (threshold), 𝛽 (slope), 𝛾 (guess rate), and 𝜆 (lapse rate). We conducted maximum likelihood estimation (MLE) of 𝜃 for each dataset using the Palamedes toolbox^111^, treating trial-level responses as outcomes of a Bernoulli process. All parameters were free, except the guess rate 𝛾 for orientation discrimination performance, which we fixed to 0.5 (corresponding to chance discriminability). Parameters were optimized in a two-stage process: first, the best parameter estimates over the following predefined grid were identified, which then served as the starting point for Nelder-Mead optimization.

𝛼: [0.05, 3] in increments of 0.05

𝛽: [10^−1^, 10^1^] in increments of 10^0.1^

𝜆: [0, 0.1] in increments of 0.01

𝛾 ∶ [0, 1] in increments of 0.1

#### Relative psychometric function

To relate objective and subjective reports over their full psychometric functions, we created a general *relative psychometric function (RPF) analysis framework*^46^. An RPF is a function that characterizes how the values of one traditional psychometric function are related to the values of another when both depend on a common independent variable (such as stimulus strength). Applying the RPF framework to the current analysis, we expressed visibility as a function of performance using RPFs of the form

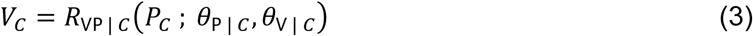

where e.g., for *C* = A, 𝑅_𝖵𝖯 | 𝖠_ is the RPF relating visibility to task performance under attention. This RPF can be constructed from Eq. 1 by solving for 𝑥 = 𝜓^−1^ (𝑃_𝖠_; 𝜃_𝖯 | 𝖠_) and substituting the result into 𝑉_𝖠_ = 𝜓_𝖵 | 𝖠_(𝑥; 𝜃_𝖵 | 𝖠_). Because 𝑅_𝖵𝖯 | 𝖠_ is not fitted directly, but rather is constructed from the separate fitted functions 𝜓_𝖯 | 𝖠_ and 𝜓_𝖵 | 𝖠_, its parameters correspond to the combined parameters of both functions, i.e., 𝜃_𝖯 | 𝖠_ and 𝜃_𝖵 | 𝖠_. Similar considerations apply for 𝑅_𝖵𝖯 | 𝖭_ and 𝑅_𝖵𝖯 | 𝖴_.

Given the expression for 𝜓 in Eq. 2, the full form of the corresponding RPF is

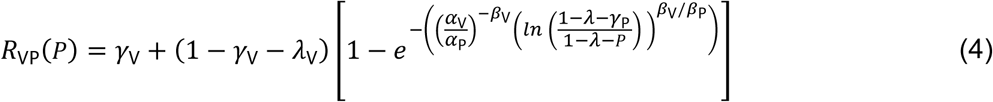

where subscripts for attentional condition have been suppressed for concision, and parameters subscripted with P and V correspond to parameters from 𝜃_𝖯 | 𝐶_ and 𝜃_𝖵 | 𝐶_, respectively.

Full details of the relative psychometric function analysis framework can be found in Maniscalco et al.^46^

#### Area under the relative psychometric function

To quantify subjective reports over a given range of objective performance, following procedures established by Maniscalco and colleagues^46^, we calculated the area under the relative psychometric function (“area under the curve”, AUC). For each attention condition, we approximated the cumulative integral of 𝑅_𝖵𝖯_ using the trapezoidal method, partitioning *P* into segments of 0.01 and computing *V* at each value using Eq. 4. We constrained *P* to a matched range across attention conditions (shaded gray regions in **Figures 2c, 3c**) per participant. For all attention conditions of each participant, we set the lower bound of *P* to 0.5 (i.e., chance performance), capitalizing on the fact that psychometric function fits ensured defined values for *V* at *P* = 0.5. Separately for each participant, we set the upper bound of *P* to the maximum fitted *P* value *within* each attention condition that was minimal *across* conditions. This maximized the *P* interval used to compute AUC for each participant, subject to the constraint that AUC be computed using the same *P* interval for each within-participant condition. This constraint ensured that any across-condition modulation in AUC was due only to differences in *V* over a fixed *P* interval. To measure subjective inflation, we compared the AUC across varying levels of attention. A larger AUC indicates higher reported visibility for a given condition over a common range of objective performance.

#### Attentional modulation index

To compare the degree and direction with which subjective reports dissociate from objective performance with attention across different experiments, we calculated an attentional modulation index (AMI) for each participant as the difference in AUC between the attended (A) and unattended (U) conditions as a proportion of their summed AUC.

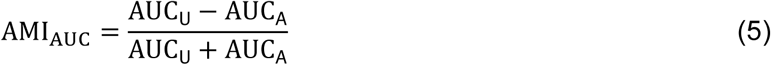

#### Signal detection theory

We calculated signal detection theory measures^61,112^ of sensitivity (*d’*) and criterion (*c*_YN_) using Eq. 6-7 for each participant and attention condition, where 𝑧 is the inverse of the normal cumulative distribution function and *c*_YN_ denotes the criterion for the yes-no discrimination task (see **Supplementary Note 2**). To handle conditions with hit (𝐻) or false alarm (𝐹) rates of 0 or 1, for which the signal detection measures are indeterminate, we applied a log-linear correction^113^, adding 0.5 to all response condition counts and 1 to all stimulus condition counts. These measures assume equal variance of the internal signal and noise distributions.

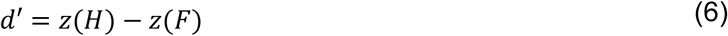

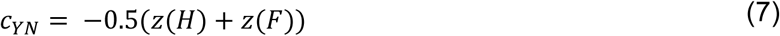

We also calculated the signal detection variants *d_a_*and *c_a_* using Eq. 8-9. These measures relax the equal variance assumption, but require estimating 𝑠, a slope parameter of the 𝑧-transformed receiver operator characteristic (ROC) curve (hit vs. false alarm rates). We leveraged our two subjective measures (of seeing the stimulus and of seeing the task-relevant feature) to estimate 𝑠 **(Supplementary Figure 6c)**, taking the slope of the line between the two points in ROC space.

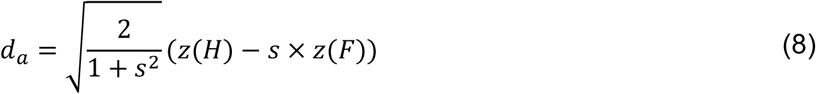

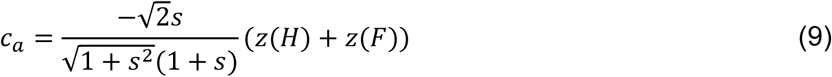

#### Attentional effects on performance and visibility thresholds

For a given unattended anchor stimulus 𝑥_𝖴_ yielding performance *P’* and visibility *V’,* we define the stimulus-anchored attentional threshold shifts for performance and visibility as

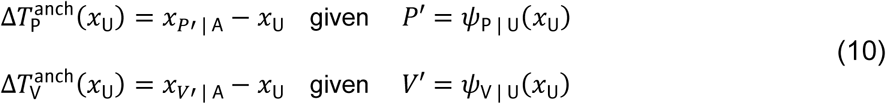

where 𝑥_𝑃′ | 𝖠_ and 𝑥_𝑉′ | 𝖠_ are threshold stimulus strengths yielding *P’* and *V’* under attention. Since 𝑥_𝖴_ is (by definition) the threshold stimulus strength yielding both *P’* and *V’* under inattention, 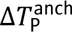 and 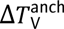 measure how attention shifts thresholds for *P’* and *V’* occurring for a common unattended anchor stimulus 𝑥_𝖴_. Comparing these measures allows us to assess how attention differentially affects thresholds for performance and visibility relative to the fixed reference of the anchor stimulus. For further details, see **Supplementary Note 4**.

For each participant, we used psychometric function fits for performance and visibility in attended (validly cued) and unattended (invalidly cued) conditions (see Methods, Psychometric function fits) to calculate 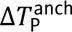 and 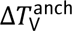 over a wide range of values for 𝑥_𝖴_. Because threshold shifts are not informative in the asymptotic regime of the psychometric function, we excluded data points beyond which the psychometric function approached asymptote, defined as the stimulus strength 𝑥_𝗉98_ yielding 0.98 ⋅ (1 − 𝜆), where 𝜆 is the fitted lapse rate. We calculated the stimulus-anchored threshold shift for each value within a usable range of anchor strengths between 0 and 𝑥_𝗉98_ in steps of 0.01.

To quantify the attentional threshold shifts across the full range of anchor strengths, we computed the signed ‘area’ under the threshold shift vs. anchor strength curve (Δ𝑇^𝘢𝘯𝘤𝘩^ AUC) using the trapezoidal method with segment sizes of 0.01. Because the usable range of anchor strengths differed across participants and measures, we restricted the AUC computation to a common range of anchor strengths across measures per participant. The Δ𝑇^anch^ AUC can take on either positive or negative values, corresponding to whether Δ𝑇^anch^ is entirely or largely above or below zero over the range of anchor strengths used. A negative Δ𝑇^anch^ AUC indicates that across all probed anchor strengths, attention tends to decrease thresholds (i.e., enhance performance or visibility); a positive Δ𝑇^anch^ AUC indicates the opposite.

One participant’s data in Experiment 3 was excluded from these analyses because of unusually shallow psychometric curves, which resulted in AUCs that were more than 3 SDs from the group mean. Four other participants’ data (one in each of Experiments 1, 2, and 4 for stimulus visibility, and one in Experiment 2 for feature visibility) were excluded because of non-overlapping psychometric functions on the *y*-axis between attention conditions, for which the stimulus-anchored threshold shift is not calculable.

#### Statistical analysis

Mixed ANOVAs were performed in R (v4.3.1; R Core Team 2023) and MATLAB to evaluate: 1) the effects of stimulus strength and attention on objective and subjective reports, including signal detection theory measures, 2) the effects of attention on the AUC, and 3) the effects of stimulus and task conditions on the AMI of the AUC. The within-subject factors included attention (valid, neutral, or invalid, with respect to the match between the precue and response cue), stimulus strength level, and visibility report type (of the overall stimulus vs. the task-relevant feature). The between-subjects factors included experimental site (BU vs. UCI) and experiment (1-4). When applicable, experiment was factorized into stimulus type (gratings vs. texture-defined figures) and task type (threshold detection vs. suprathreshold visibility comparison). All statistical tests were two-sided. Effect sizes are reported as generalized eta-squared. When Mauchly’s test indicated violations of sphericity assumptions^114^, we confirmed that all significant *F* tests remained significant after Greenhouse-Geisser correction^115^, with the estimated degree of sphericity violation reported as epsilon (𝜀). Planned analyses were conducted per experiment. Planned comparisons were made between stimulus types, task types, visibility report types, and all pairs of cueing validity and stimulus strengths, when applicable.

## Acknowledgments

This research was supported by the Templeton World Charity Foundation Accelerating Research on Consciousness Initiative TWCF 0567 to BJH, JWB, NB, DC, MAKP, and RND, National Defense Science and Engineering Graduate Fellowship to KJT, startup funding from the University of California Irvine and support from the Canadian Institute of Advanced Research Brain, Mind, & Consciousness program to MAKP, and startup funding from Boston University to RND. The funders had no role in the study design, data collection and analysis, decision to publish or preparation of the manuscript. We thank Sarah Cormiea, Angus Chapman, Jeffrey Nestor, and

Brian Odegaard for helpful discussions at different stages of the project and Giancarlo Arzu and Lizbeth Romero for assistance with data collection.

## Data availability

The data will be made publicly available upon publication.

## Code availability

The code to run the experiments is available at https://github.com/denisonlab/twcf_expt1_code and the presented analyses is available at https://github.com/denisonlab/twcf_expt1_analysis.

The toolbox for fitting the relative psychometric function is available at https://github.com/CNClaboratory/RPF, with additional documentation provided in the accompanying paper by Maniscalco et al.^46^

## Author contributions

KJT, BM, MLE, AS, MAKP, and RND conceptualized and designed the experiments with feedback from RB, VAFL, HL, BJH, JWB, NB, and DC. Theory predictions as to the experimental outcomes were supplied by VAFL, RB, and HL. KJT and BM programmed the experiments. KJT, MLE, AS, OGC, TK, JAM, EO, EER, MEW, JW, TBAZ, and MAKP collected the data. KJT and BM analyzed and visualized the data with input and supervision from MAKP and RND. KJT, BM, MAKP, and RND wrote the manuscript. MLE, AS, JAM, HL, RB, BJH, JWB, NB, and DC provided feedback on the manuscript draft. All authors approved the final version.

