## Supplementary Information for "When awareness outstrips performance: critical tests of subjective inflation under inattention"

Department of Psychological and Brain Sciences
Boston University
111 Cummington Mall
Boston, MA 02215
United States



Keywords: spatial attention, awareness, subjective inflation, visual perception, consciousness

### Supplementary Information

#### Supplementary Figures

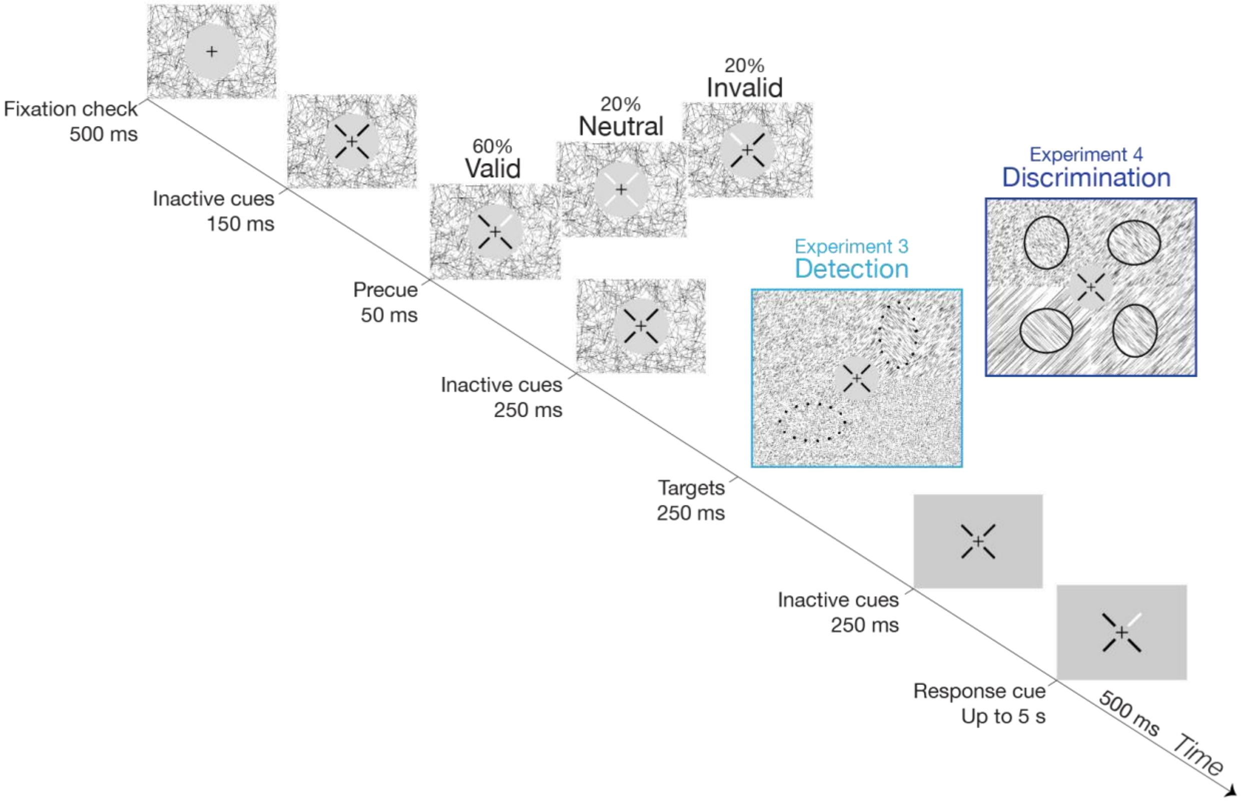


**Supplementary Figure 1. Detailed task timeline for figure-ground experiments.** Expansion of Figure 1. Each trial was initiated by a fixation check. From fixation until target offset, the central fixation cross, attentional precues, and inactive cues were presented in a circular gray aperture (4° diameter). The aperture was placed atop a texture composed of lines of random orientations and lengths. The target display was divided into quadrants, each filled with a unique "background" texture composed of lines of a single orientation (-45° or +45°) and length. An oval "figure" delineated by orthogonal lines appeared in each quadrant with 50% (Experiment 3) or 100% (Experiment 4) probability.

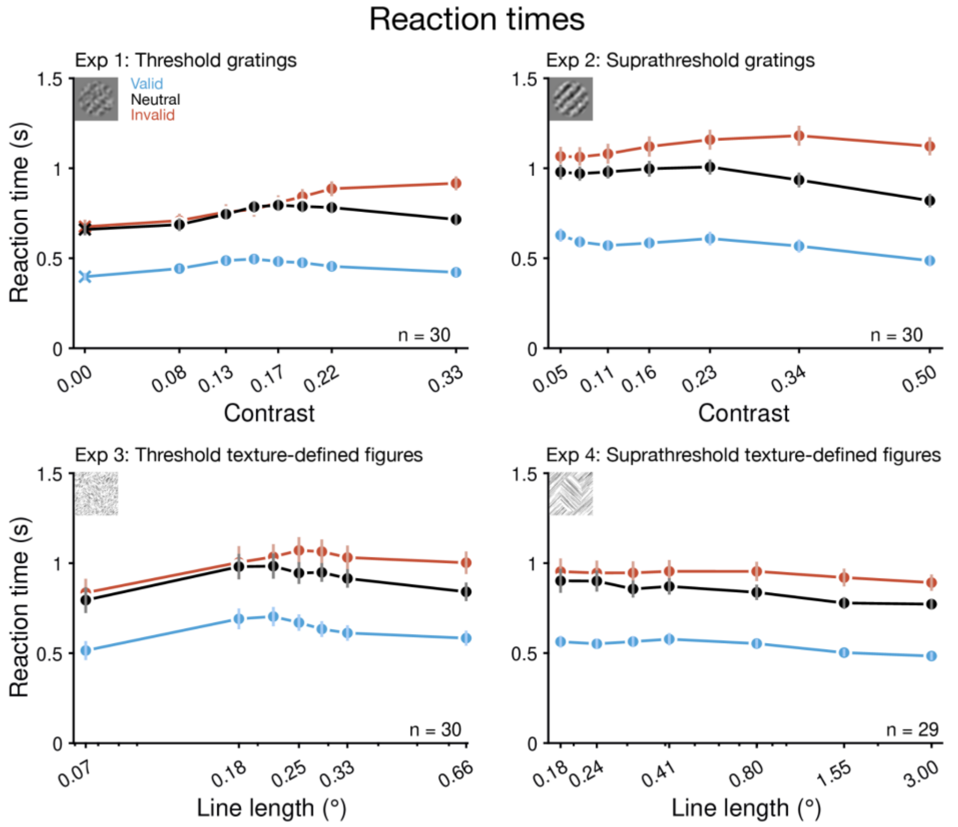


**Supplementary Figure 2. Reaction times.** Reaction times were fastest on valid trials (blue), intermediate on neutral trials (gray), and slowest on invalid trials (red). Data (total n=120; Experiments 1-3 each n=30, Experiment 4 n=29) are presented as mean values ±1 SEM.

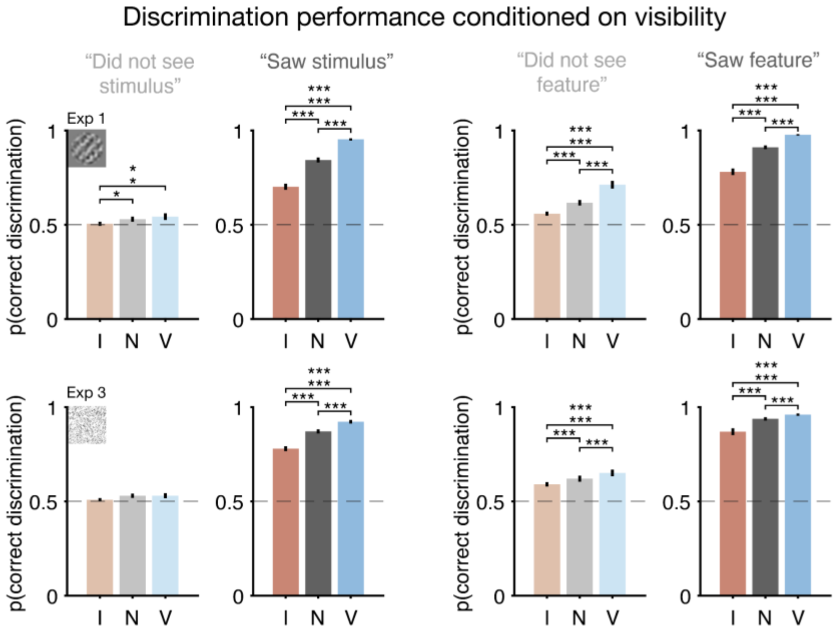


**Supplementary Figure 3. Discrimination performance conditioned on subjective visibility in the detection experiments.** Experiment 1 (top) and 3 (bottom). The attentional cue modulated performance even when the stimuli were reported as unseen. *$p$<0.05, **$p$<0.01, ***$p$<0.001. Data (total n=60; Experiments 1 and 3 each n=30) are presented as mean values ±1 SEM.

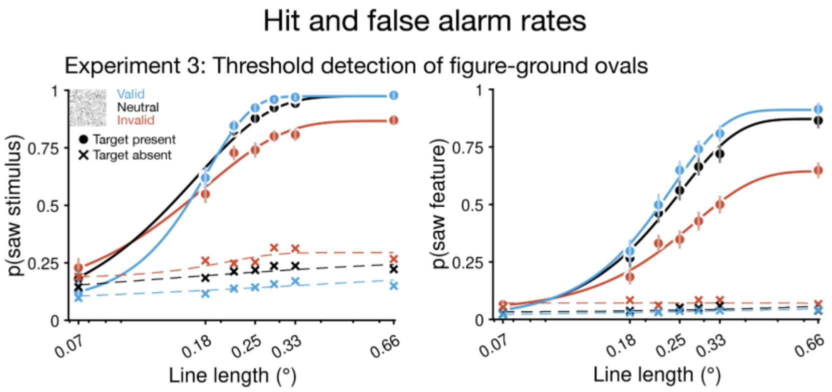


**Supplementary Figure 4. Hit and false alarm rates for figure-ground detection.** Each texture line length corresponded to unique target-present (solid lines) and target-absent (dashed lines) trials, allowing hit and false alarm rates to be estimated separately at each stimulus strength. Stimulus visibility (left) and feature visibility (right). Inattention (red) decreased hit rates, except at the lowest line length, and increased false alarm rates, for both subjective report types. Group means are fit with Weibull functions for visualization. Data (Experiment 3 n=30) are presented as mean values ±1 SEM.

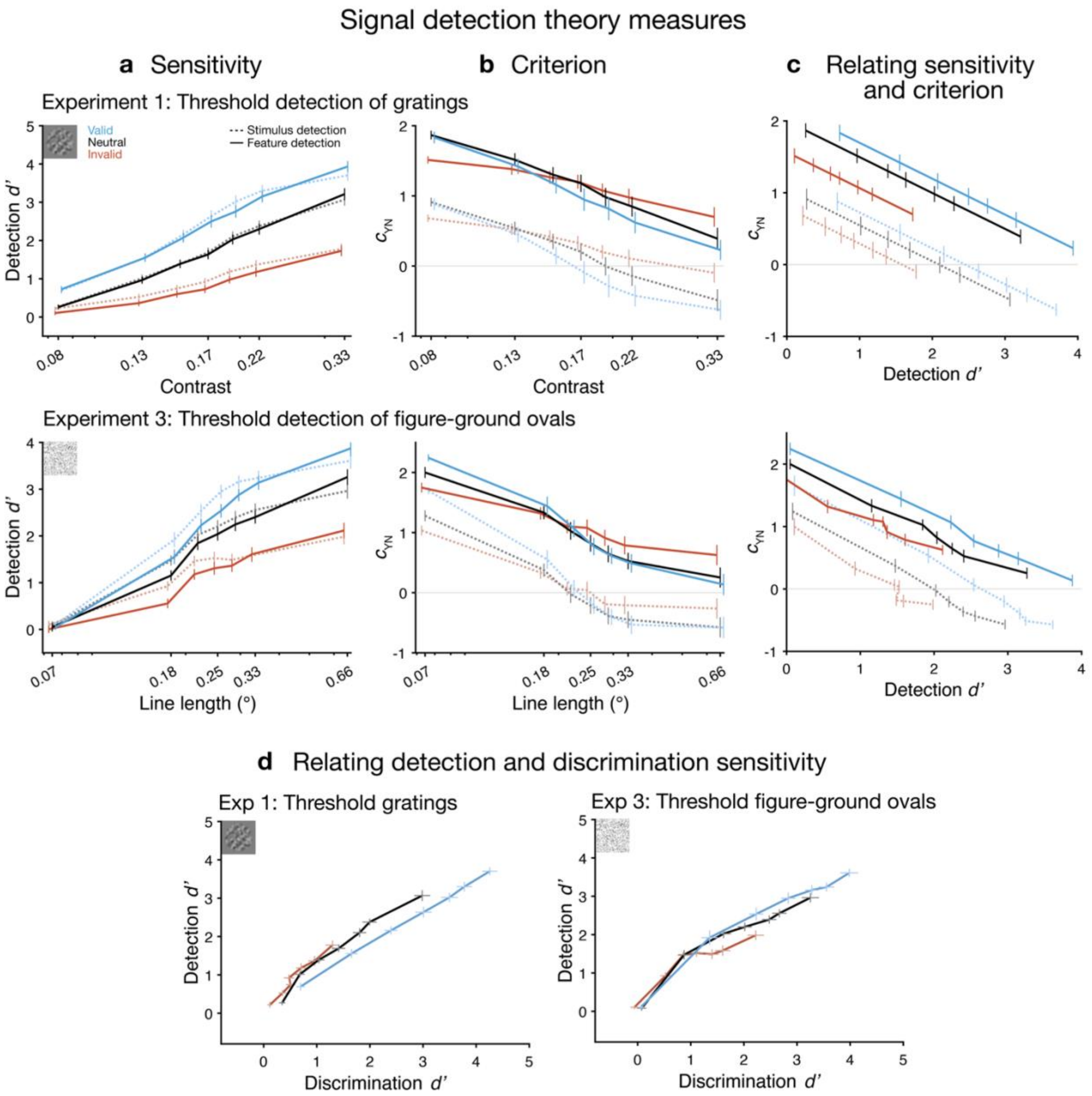


**Supplementary Figure 5. Signal detection theory measures. a)** Detection sensitivity (*d'*) increased with stimulus strength and attention (valid = blue, neutral = gray, invalid = red). **b)** Criterion ($c_{\mathrm{YN}}$) was more liberal for stimulus visibility (dashed lines) than feature visibility (solid lines), whereas detection sensitivity was similar for both report types. Attention induced a more conservative criterion at lower stimulus strengths but a more liberal criterion at higher stimulus strengths (see **Supplementary Note 2** for an argument that these changes in criterion are driven by changes in *d'*). **c)** At matched detection sensitivity, attention consistently induced a more conservative criterion. **d)** Detection vs. discrimination sensitivity. Detection sensitivity was not systematically higher for unattended trials at matched discriminability. Data (total n=60; Experiments 1 and 3 each n=30) are presented as mean values ±1 SEM.

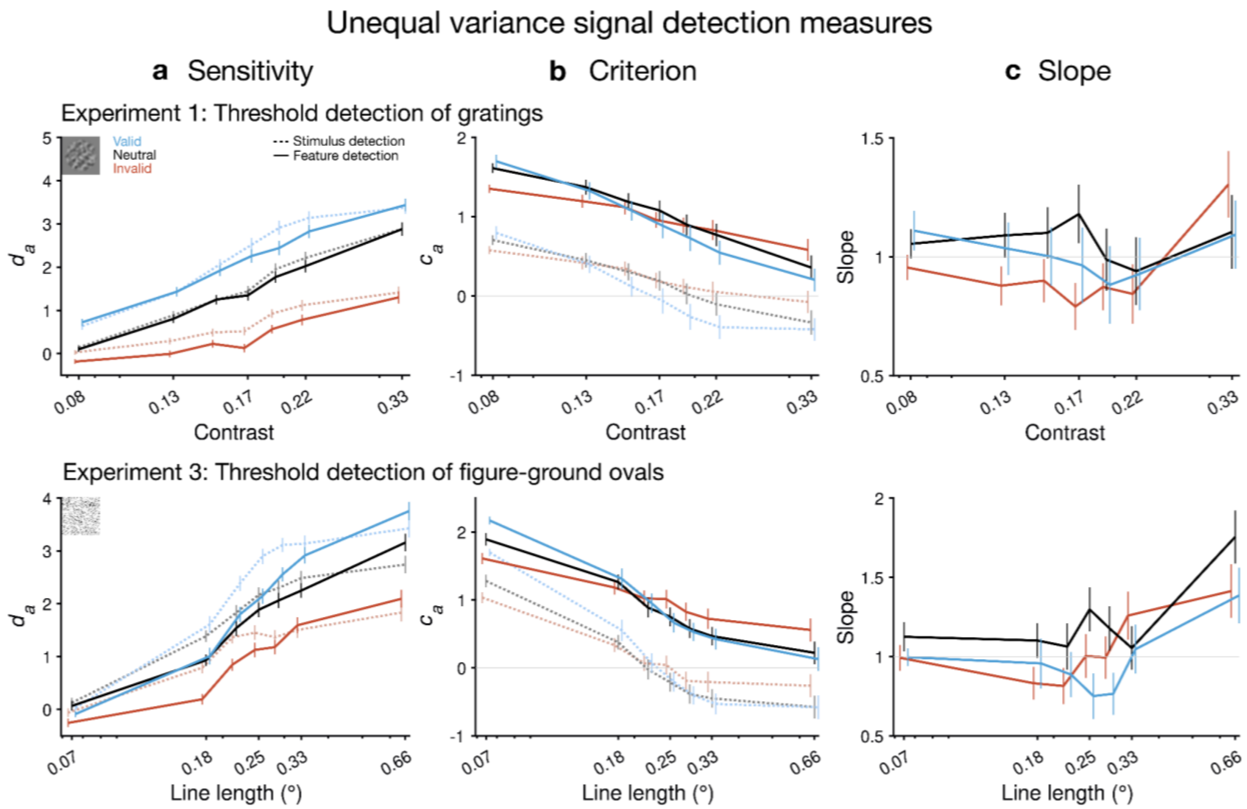


**Supplementary Figure 6. Unequal variance signal detection theory measures.** Effects of stimulus strength and attention on **a)** detection sensitivity $d_a$ and **b)** criterion $c_a$ were consistent with their effects on the equal variance SDT measures (Supplementary Figure 5). **c)** The $z$ROC slope did not deviate substantially from 1, indicating that equal variance assumptions were not violated in most cases, apart from at the highest stimulus strengths. Data (total n=60; Experiments 1 and 3 each n=30) are presented as mean values ±1 SEM.

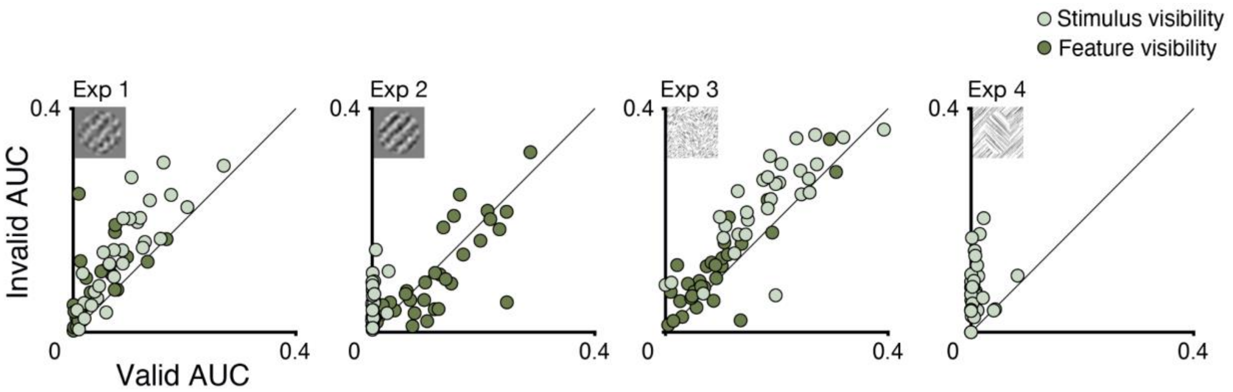


**Supplementary Figure 7. Consistency of inattentional inflation across participants.** Each dot shows an individual participant's AUC for invalid (*y*-axis) vs. valid (*x*-axis) trials. For most participants, the AUC for stimulus visibility (light green) was larger on invalid than valid trials (above the unity line), indicating inattentional inflation. The AUC for feature visibility (dark green) was also larger on invalid trials for most participants in Experiments 1 and 3.

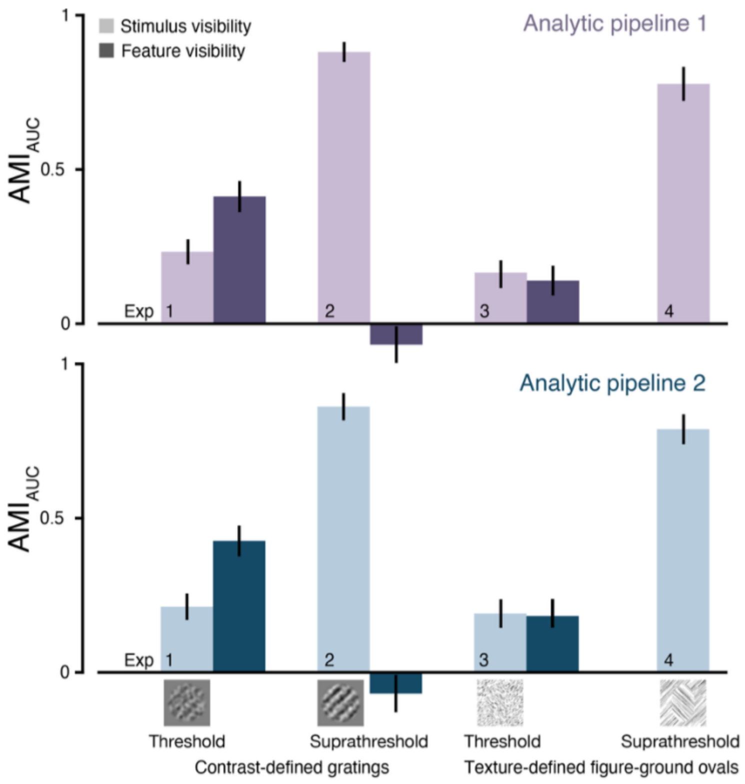


**Supplementary Figure 8. AMI replicated across analytic pipelines.** The attentional modulation index (AMI), calculated from raw data using independent analytic pipelines developed at two study sites: BU (top) and UCI (bottom). AMIs did not differ by pipeline (all $p$>0.657, Supplementary Table 16). Data (total n=118; Experiments 1-3 each n=30 per experiment, Experiment 4 n=28) are presented as mean values ±1 SEM.

#### Supplementary Notes

##### Supplementary Note 1

This experiment was conceived as part of an adversarial collaboration between first-order and higher-order theories of consciousness, as described in the preregistration[1]. The specific theories under consideration were Recurrent Processing Theory[2] (RPT), a first-order theory, and two higher-order theories: Perceptual Reality Monitoring theory[3] (PRM) and Higher-Order Representation of a Representation theory[4,5] (HOROR).

The full adversarial collaboration includes two suites of experiments, involving subjective inflation and change blindness, each with two phases: psychophysics and fMRI. The current manuscript describes the outcomes of the psychophysics phase of the subjective inflation experiments. As described in the preregistered predictions table[1], only the fMRI phase has the potential to pose a serious challenge to one theory that would require a revision of the theory—denoted as a "fail" outcome in the predictions table. However, the theorists representing each theory also made predictions for the behavioral phase. Here, predictions unsupported by experimental outcomes would prompt reconsideration of some aspect of the theory but would not invalidate a core theoretical component—denoted as a "challenge" outcome in the predictions table.

The theorists representing the higher-order theories tested here (PRM and HOROR) predicted that the behavioral experiments would show inattentional subjective inflation. An outcome in which no behavioral experiment showed inflation would have challenged the higher-order theories by invalidating a motivating pillar of these theories. If inflation had not been supported in the current rigorous test, it would have undermined the idea that subjective experience regularly exceeds objective performance, reducing the empirical basis for the theoretical separation of sensory processing and conscious experience inherent to higher-order theories.

The theorist representing the first-order theory tested here (RPT) predicted in contrast that the behavioral experiments would not show inattentional subjective inflation. This prediction stemmed from the absence of a mechanism within RPT to generate inflation: in RPT, recurrent sensory processing is proposed to generate both conscious experience and performance in sensory tasks. However, given that inflation can in principle be generated by first-order signal detection models (see Discussion), a behavioral finding of inflation would not invalidate a core theoretical component of RPT. Rather, the current findings of robust inflation prompt RPT and other first-order theories to develop or incorporate more explicit mechanisms to account for decouplings of objective and subjective reports.

#### Supplementary Note 2

*Overview of the criterion analysis of Figure 4*

In **Figure 4a**, we plot the difference between detection criteria for attended and unattended trials as a function of stimulus strength, following the similar analysis in Figure 2b of Rahnev et al.[6] for their Experiment 2. Rahnev et al.[6] probed detection of peripherally presented gratings at four levels of near-threshold grating contrast when these stimuli were either validly or invalidly cued, making their experimental design similar to that of our Experiment 1. The results in our **Figure 4a** replicate those of Rahnev et al.'s[6] Figure 2b in showing that 1) detection criteria for peripheral gratings are higher (more conservative) for attended trials than for unattended trials at low grating contrasts, and 2) the difference between criteria for attended vs. unattended trials decreases as grating contrast increases. Moreover, our results confirm a prediction made by Rahnev et al.'s[6] computational model of their data by showing that if high enough contrasts are probed, the criterion difference reverses, such that detection criteria are increasingly lower (more liberal) for attended trials as contrast increases—though that study did not note that prediction and instead focused on the finding that attention induces conservative detection criteria. We also extend the results of Rahnev et al.[6] by 1) showing these criterion patterns hold not just for stimulus detection (i.e., reporting that a grating was visible), but also for feature detection (i.e., reporting that the grating's tilt was visible); and 2) showing that the same patterns for stimulus and feature detection criteria hold for texture-defined ovals (Experiment 3).

*Distinguishing the criterion in yes-no discrimination tasks vs. two-response classification tasks*

The analysis of **Figure 4a** requires important conceptual context. Our Experiment 1 uses a detection task featuring a single set of target-absent trials (corresponding to grating contrast = 0) and multiple sets of target-present trials (corresponding to contrasts > 0). The observer must provide two binary classifications for each stimulus, corresponding to stimulus detection ("saw grating" vs. "did not see grating") and feature detection ("saw grating tilt" vs. "did not see grating tilt"). For simplicity, in the following discussion we will consider only the stimulus detection task, but all considerations similarly apply to modeling the feature detection task. In signal detection theory (SDT), the stimulus detection task of Experiment 1 is treated as a *two-response classification task*[7] in which the observer sets a single criterion on a decision axis to classify more than two stimulus categories (here, target-absent trials and target-present trials at multiple contrasts) into two classes.

Importantly, the criterion in two-response classification tasks is computed and interpreted differently from the criterion used to model the more common *yes-no discrimination task*[7]. Here we clarify this distinction to prevent potential confusions that might arise from failing to do so, and to facilitate correct interpretation of our analysis in **Figure 4a**.

In the yes-no discrimination task, the observer is presented with a stimulus from one of two stimulus categories (e.g., target-absent and target-present) and must provide a binary classification (e.g., "yes, saw target" or "no, did not see target"). In the yes-no SDT model, the two stimulus categories generate normal distributions of evidence along some decision axis. The observer sets a criterion on the decision axis such that they respond "yes" for any trial yielding evidence that surpasses the criterion, and "no" otherwise. Let us call the criterion for the yes-no discrimination task the *yes-no criterion*, or $c_{\mathrm{YN}}$. Its formula is given by

$$c_{\mathrm{YN}} = -0.5\,(z(H) + z(F)) \tag{S1}$$

where $H$ and $F$ correspond to hit rate and false alarm rate, respectively, and $z$ is the inverse of the normal CDF. $c_{\mathrm{YN}}$ is measured relative to a coordinate system whose zero occurs at the location on the decision axis where the two stimulus distributions intersect (i.e., have equal likelihood). This is a point of zero response bias in the sense that setting the criterion here yields an equal error rate for "yes" and "no" responses (i.e., equal false alarm rate and miss rate). It follows that the sign of the yes-no criterion can be interpreted in terms of bias in the decision-making strategy. Setting the criterion above the zero-bias point ($c_{\mathrm{YN}} > 0$) is a conservative strategy that prioritizes decreasing false alarm rate at the expense of increasing miss rate, whereas setting it below the zero-bias point ($c_{\mathrm{YN}} < 0$) is a liberal strategy that prioritizes decreasing miss rate at the expense of increasing false alarm rate.

The SDT model of the two-response classification task is identical to that of the yes-no task, except that there are more than two stimulus distributions. It follows that for this model, there is not a unique zero-bias point. Every possible pairing of stimulus distributions yields a different location at which the paired distributions intersect, and these correspond to different zero-bias points that are specific to each pairing. It follows that the criterion used to model such tasks, the *two-response classification criterion* or $c_{\mathrm{2RC}}$, cannot be computed and interpreted relative to a zero-bias point in a way analogous to the yes-no criterion; instead, a new choice must be made for the zero of the measurement scale. One reasonable approach is to compute the location of $c_{\mathrm{2RC}}$ relative to the mean of the noise distribution[a]:

$$c_{2RC} = -z(F) \tag{S2}$$

Whatever convention for the zero point is chosen, the sign of $c_{\mathrm{2RC}}$ cannot be interpreted as reflecting bias in the decision-making strategy in the same way as $c_{\mathrm{YN}}$.

*Interpreting yes-no criterion effects in Experiment 1*

Bearing these distinctions between $c_{\mathrm{YN}}$ and $c_{\mathrm{2RC}}$ in mind, we nonetheless chose to analyze the criterion for the two-response classification task of Experiment 1 using a yes-no analysis framework. We explain the rationale for this choice in the following section; here, we examine why this choice leads to results for Experiment 1 that are trivial and potentially misleading if not properly understood.

Applying a two-response classification SDT model to Experiment 1 would involve computing a single $c_{\mathrm{2RC}}$ value for each attention condition of each participant, where this value corresponds to the single criterion that determines stimulus detection responses for all stimuli. Instead, for each attention condition of each participant, we applied the yes-no discrimination SDT model

[a] Technically, a two-response classification task does not necessarily have to include target-absent or "noise" trials. In this case, the criterion could be computed relative to the mean of the weakest stimulus distribution, with p("yes") for these stimuli being the analogue of false alarm rate.

separately for all possible pairings of the target-present trials at each level of contrast with the common set of target-absent trials. This yielded separate values of $c_{\mathrm{YN}}$ at every contrast. We observed that $c_{\mathrm{YN}}$ decreased with contrast (**Supplementary Figure 5b**), and that the rate of this decrease depended on attention (**Figure 4a**).

Taken at face value, this would seem to suggest that participants employed an increasingly liberal criterion-setting strategy as contrast increased, and that this criterion setting effect was modulated by attention. However, this cannot possibly be the case, since each $c_{\mathrm{YN}}$ within an attention condition is computed from the same false alarm rate arising from the same set of target-absent trials, and therefore must correspond to the *same criterion*, i.e., the fixed two-response classification criterion. This single underlying criterion is assigned different values in the pairwise yes-no analyses due to the fact that the zero-bias point for a given pairing of noise and signal distributions (i.e., the location where they intersect) differs for each level of contrast (**Supplementary Figure 9**). As contrast increases, so does the mean of the corresponding target-present distribution, causing a rightward shift in the zero-bias point and a corresponding decrease in the computed value for $c_{\mathrm{YN}}$. Thus, within each attention condition, the observer sets only one criterion which applies across all contrasts (fixed $c_{\mathrm{2RC}}$), and this single criterion exhibits different relationships to the zero-bias point for each pairing of noise and signal distributions (yielding different $c_{YN}$ values)—but these do not reflect substantive changes in criterion setting *per se*.

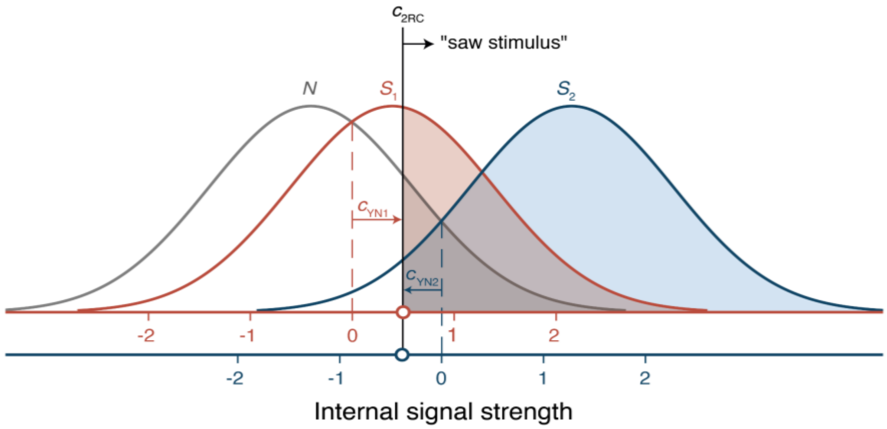


**Supplementary Figure 9. Signal detection theory (SDT) analysis schematic.** Schematic for SDT analysis of a two-response classification task with one set of target-absent trials (corresponding to the noise distribution $N$) and two sets of target-present trials with different stimulus strengths (corresponding to the signal distributions $S_1$ and $S_2$). A single two-response classification criterion $c_{\mathrm{2RC}}$ determines one false alarm rate from $N$ and two hit rates from $S_1$ and $S_2$, respectively. However, when SDT analysis for a yes-no discrimination task is conducted separately for each stimulus strength, it yields two different values ($c_{\mathrm{YN1}}$ and $c_{\mathrm{YN2}}$) for the location of the same underlying criterion $c_{\mathrm{2RC}}$. These values differ because the yes-no criterion is measured relative to the location where the noise and signal distributions intersect, and this zero-bias point depends on the mean of the signal distribution, which changes with signal strength (compare the red and blue axis coordinates derived from $S_1$ and $S_2$, respectively). Here, these different coordinate systems render $c_{\mathrm{2RC}}$ as positive (i.e., conservative) in $c_{\mathrm{YN1}}$ and as negative (i.e., liberal) in $c_{\mathrm{YN2}}$. As the signal distribution mean increases, the zero-bias point shifts rightwards, leading to increasingly negative values for the yes-no criterion $c_{\mathrm{YN}}$ despite the underlying two-response classification criterion $c_{\mathrm{2RC}}$ being fixed. Thus, although measurements for $c_{\mathrm{YN}}$ become more negative as stimulus strength increases, this is best understood not as an actual shift in criterion setting *per se*, but rather as a systematic change in how the fixed criterion $c_{\mathrm{2RC}}$ relates to the changing zero-bias point as sensitivity increases.

Because these changes in $c_{\mathrm{YN}}$ are driven entirely by changes in signal distribution mean, there is a simple relationship between $c_{\mathrm{YN}}$ and sensitivity (*d'*). With every unit increase in *d'*, the zero-bias point shifts rightwards by 0.5 units (since it is located at the mean of the noise and signal distribution means). This entails that $c_{\mathrm{YN}}$ is linearly related to *d'* with a slope of -0.5 (**Supplementary Figure 5c**, top row). As a consequence, changes in $c_{\mathrm{YN}}$ with contrast trivially reflect changes in *d'* with contrast (compare blue plots in **Figure 4** panels **a** and **b**; also top row of **Supplementary Figure 5** panels **a** and **b**). Thus, in two-response classification tasks, not only do changes of $c_{\mathrm{YN}}$ with contrast not reflect changes in criterion setting, in fact they are best understood as sensitivity effects insofar as they indirectly reflect changes in *d'*. Likewise, the decreasing difference between $c_{\mathrm{YN}}$ for attended and unattended trials with increasing contrast (**Figure 4a**, blue plots) does not reflect an attentional modulation of criterion setting across contrasts, but rather reflects that *d'* increases more rapidly with contrast for attended stimuli (**Figure 4b**, blue plots).

*Interpreting yes-no criterion effects in Experiment 3 and Rahnev et al. (2011)*

If the $c_{\mathrm{YN}}$ results for Experiment 1 do not inform us about criterion setting but rather are trivial reflections of the effects of contrast and attention on *d'*, what motivates using the yes-no analysis framework here? The answer is that this analysis as applied to Experiment 1 provides a benchmark for comparison to the $c_{\mathrm{YN}}$ results of our Experiment 3 and Rahnev et al.'s[6] Experiment 2. For reasons discussed below, the $c_{YN}$ analysis is not trivial *a priori* for either of these experiments. However, if these experiments nonetheless exhibit patterns in the $c_{\mathrm{YN}}$ results similar to Experiment 1, this would suggest that the findings of these experiments might best be interpreted in a similar way—i.e., as reflecting not criterion-setting effects *per se*, but rather sensitivity effects.

Although the stimulus detection task for texture-defined oval stimuli in Experiment 3 is structurally similar to the task of Experiment 1 in many ways, for the purposes of the present discussion there is an important difference. In Experiment 3, the presence or absence of the oval is determined by line orientation, which is independent of the stimulus strength manipulation of line length. It follows that each level of line length has its own set of target-absent trials (all lines parallel) and target-present trials (figure lines orthogonal to ground lines). These data are most naturally analyzed not with a two-response classification SDT model as in Experiment 1, but rather with separate yes-no SDT models applied to each line length. Because the yes-no SDT analyses conducted at each line length do not use a common set of target-absent trials, it is not necessarily the case that the $c_{\mathrm{YN}}$ computed at each line length merely reflects a single underlying criterion measured relative to different coordinate systems, as is the case in Experiment 1. Rather, they may reflect real changes in criterion setting induced by perceptible changes in line length.

However, false alarm rates in Experiment 3 exhibited only slight changes as a function of line length (**Supplementary Figure 4**), approximating the constant false alarm rate from the common set of stimulus-absent trials in Experiment 1. This implies that changes in $c_{\mathrm{YN}}$ as a function of line length were driven primarily by changes in hit rate, i.e., by changes in the mean of the signal distribution with stimulus strength—again approximating the sensitivity-driven effects in Experiment 1. These patterns in the Experiment 3 data resulted in criteria being very nearly linear with sensitivity and with a slope of approximately -0.5 (**Supplementary Figure 5c**, bottom row), in close approximation to the (trivially) perfectly linear psychometric functions of slope -0.5 in Experiment 1 (**Supplementary Figure 5c**, top row).

These data suggest that although the $c_{\mathrm{YN}}$ patterns in Experiment 3 are not trivial *a priori* as they are in Experiment 1, nonetheless they closely approximate the patterns in Experiment 1 and therefore may be best understood in the same way—i.e., as reflecting effects of attention on sensitivity, not criterion setting. On this interpretation, the observer sets a fixed criterion for all line lengths within an attention condition. Provided that the means and variances of the noise distributions are roughly constant across line lengths, this fixed criterion would yield approximately constant false alarm rates[b] and increasing hit rates across line lengths. This would create a situation similar to that of the two-response classification task, where $c_{\mathrm{YN}}$ values decrease with stimulus strength as a side-effect of the zero-bias point increasing with *d'* despite the underlying criterion being fixed (**Supplementary Figure 9**).

Experiment 2 of Rahnev et al.[6] used a grating detection task similar to our Experiment 1, which we argued above is best modeled as a two-response classification task. However, there is an important difference in how these two experiments were conducted. In our Experiment 1, target-absent trials and target-present trials of all contrasts were interleaved randomly across trials. In Rahnev et al.'s[6] Experiment 2, each block of trials contained a mix of target-absent trials and target-present trials of a fixed contrast; grating contrast varied across but not within different blocks. This design provides a natural way to pair separate sets of target-absent trials with target-present trials at each contrast, which justifies treating the data as a series of independent yes-no tasks at each contrast rather than as an omnibus two-response classification task using a common set of target-absent trials. In turn, this structure allows for the possibility that changes in $c_{\mathrm{YN}}$ with contrast reflect real changes in criterion setting. For instance, observers might adjust their detection criterion in each block due to perceptible differences in grating contrast. Nonetheless, Rahnev et al.[6] observed that false alarm rates for attended and unattended stimuli were roughly constant across contrasts (their Supplementary Figure 4), suggesting that the effects of contrast and attention on $c_{\mathrm{YN}}$ they observed (their Figure 2b) may be better understood as an indirect reflection of effects of contrast and attention on *d'*, per the above considerations.

In fact, this idea is well accommodated by the analysis of Rahnev et al.[6] Although they analyzed their empirical data in terms of $c_{\mathrm{YN}}$ effects, they proposed a deeper computational model that explains the data in terms of how attention influences sensitivity, not criterion. According to their model, attention both boosts signal magnitude and decreases the variance of perceptual evidence. Under the assumption that the same criterion is used for attended and unattended stimuli, this model predicts that false alarm rates are higher under inattention[c]. Provided that the model parameters are tuned in the appropriate way, the model can yield a higher $c_{\mathrm{YN}}$ value for attended stimuli at low stimulus strengths, in agreement with empirical data. Furthermore, given the model assumptions that 1) the distribution means for unattended stimuli increase more slowly with stimulus strength than those for attended stimuli and 2) the distribution variances for unattended stimuli are always higher than variances for attended ones at each stimulus strength, it follows that sensitivity (*d'*) increases with stimulus strength more slowly for unattended stimuli. The model's assumption of a fixed criterion entails that $c_{\mathrm{YN}}$ decreases with stimulus strength for all attention conditions, and its prediction that *d'* increases with stimulus strength more slowly for unattended stimuli entails that the decrease of $c_{\mathrm{YN}}$ with stimulus strength will correspondingly be

[b] The slight increase in stimulus detection false alarm rates with line length (**Supplementary Figure 4**, left panel) could arise from a fixed criterion if the mean and/or variance of the noise distributions increases slightly with line length.

[c] Provided that the criterion location exceeds the mean of the noise distribution, i.e., provided that false alarm rates are less than 0.5.

slower for unattended stimuli. The slower rate of decrease for $c_{\mathrm{YN}}$ under inattention entails that the difference between criteria for attended and unattended stimuli decreases as stimulus strength increases from low levels, eventually yielding a crossover such that $c_{\mathrm{YN}}$ is lower for attended stimuli at high enough stimulus strengths. This is consistent with the pattern in our data (**Figure 4a**) as well as Rahnev et al.'s[6] data (their Figure 2b) and model fits (their Supplementary Figure 6b, where the model's predicted crossover begins to emerge at the highest plotted stimulus strength, in a straightforward extrapolation of the patterns observed at lower stimulus strengths.)

Since Rahnev et al.'s[6] experiment used low-contrast gratings, their empirical results stopped just short of exhibiting this crossover effect and their interpretation emphasized the effect of attention on making $c_{\mathrm{YN}}$ conservative at low stimulus strengths. However, importantly, their computational model explained these patterns in $c_{\mathrm{YN}}$ as arising from processes best captured by a more complex SDT-based model in which criterion is fixed and attention modulates sensitivity via effects on signal strength and variance. Thus, their modeling approach aligns with the above considerations that these patterns are best understood as effects of stimulus strength and attention on sensitivity rather than criterion-setting.

Although Rahnev et al.[6] modeled a single fixed criterion across all stimulus strengths and attention conditions, their data and ours could also be consistent with a criterion that is fixed across stimulus strengths within each attention condition, but varies with attention[8]. The present dataset will provide an opportunity for further testing and development of SDT models to better understand how attention interacts with perception and perceptual decision making.

**Supplementary Note 3**

*Here, we include discussion of one author's (B.M.'s) experiences of this task that may suggest an alternative framing for subjective inflation. We include these observations as they may be helpful in informing future research—both theoretical and empirical.*

Subjective inflation is typically framed as the surprising and unintuitive finding that, for matched levels of feature discrimination performance, subjective visibility is higher when not attending to the stimulus. However, turning this framing on its head yields the potentially more intuitive idea that, at matched levels of subjective visibility, feature discrimination performance is higher when attending to the stimulus. The general idea that strong overall visibility does not necessarily entail fine perceptual discrimination is familiar from the everyday observation that stimuli perceived in the visual periphery can feel highly prominent and yet elude attempts to discern fine-grained stimulus features. This suggests a more general principle that in impoverished viewing conditions, the subjective experience of perceptual prominence does not necessarily guarantee the subjective experience of perceptual sharpness[d]. It is perhaps not so counterintuitive to suppose that this principle might apply to cases of inattention, making fine discrimination of a strongly visible unattended stimulus more difficult than it would be for a similarly visible attended stimulus.

This framing accords well with the subjective experience of one of the authors (B.M.) when testing the task for suprathreshold texture-defined ovals (Experiment 4). Considering trials where attended and unattended stimuli appeared similar to the reference strength and thus had similarly strong levels of overall “pop-out” from the background (i.e., similar levels of "perceptual prominence"), this matched level of stimulus visibility did not feel nearly as useful for making fine discrimination judgments about the orientation of its nearly circular shape in the absence of attention (i.e., different levels of "perceptual sharpness"). This difference stood out as being strikingly obvious at the single trial level in a way that was not the case for the threshold stimuli. This is likely due to the fact that, to achieve threshold discrimination performance for clearly visible suprathreshold stimuli, the differences in stimulus features to be discriminated must be far more subtle than they are for stimuli near the detection threshold, which has the effect of making discrimination for suprathreshold stimuli feel far more challenging. This configuration maximizes the contrast between these dissociable aspects of perception, with detection being very easy and discrimination being very hard, which in turn may be a formula for making inflation effects stand out as much as possible. The net effect of all this was to make inflation effects (framed as inattentional deficits in feature discrimination for matched-visibility stimuli) seem obvious to the author at the single-trial level in a way that felt analogous to how one can clearly detect an object in the visual periphery and yet struggle to discern its fine-grained features.

Of course, this anecdotal experience is entirely consistent with the data analyzed in the main manuscript, taking "perceptual prominence" and "perceptual sharpness" to correspond to overall

[d] “Perceptual prominence” and “perceptual sharpness” are introduced here as terms to refer to phenomenological aspects of visual experience that are familiar from everyday life. The subjective reports participants made about seeing the stimulus (Experiments 1 and 3) or judging stimulus strength relative to a reference (Experiments 2 and 4) can be seen as operationalizations of the participants' experiences of the perceptual prominence of the response-cued stimuli in these experiments, and similarly reports about seeing stimulus features (Experiments 1, 2, and 3) can be seen as operationalizations of their experiences of the perceptual sharpness of these stimuli. These phenomenological and operational concepts are distinct from, but related to, the computational concepts of signal strength and signal-to-noise ratio in the model discussed below.

stimulus visibility and feature visibility, respectively. Consider the plot of subjective stimulus visibility vs. objective feature discrimination performance for Experiment 4 (**Figure 2c**, bottom panel). Taking any horizontal slice through this plot shows that at matched levels of subjective visibility, discrimination performance is considerably higher when the stimulus is attended than when unattended. Although feature visibility data were not collected for Experiment 4, feature visibility data for Experiment 2 (which used suprathreshold gratings) were closely related to discrimination accuracy in a way that did not depend on attention (**Figure 3c**, middle panel). Provided that feature visibility data in Experiment 4 would have been similar, these results jointly demonstrate that for matched levels of overall stimulus visibility for suprathreshold stimuli, subjective feature visibility (and objective feature discrimination) is considerably higher under attention.

The motivation for including this anecdotal report is not the experienced phenomena per se, as these are entirely consistent with effects already demonstrated or suggested in the data. Rather, what is notable is the force with which single trial observations of suprathreshold inflation stimuli suggested to the author how this alternative framing of inflation effects naturally and intuitively accords with similar experiences familiar from everyday, suprathreshold vision—contra the typical framing of inflation effects as unintuitive and surprising. It is possible that deeper consideration of this alternative framing could lead to insightful new research angles and perhaps stimulate new paradigms for understanding inflation effects and their relationship to similar phenomena in everyday suprathreshold vision.

This framing accords well with the signal detection model of Rahnev et al.[6] As explained in greater detail in the Discussion and in **Supplementary Note 2**, this model explains inflation effects as resulting (in part) from higher levels of noise in perceptual representations of unattended stimuli. At matched levels of signal-to-noise ratio (and so, matched levels of objective performance), the noisier perceptual evidence for unattended stimuli has higher overall magnitude (and so higher subjective visibility). But turning this framing on its head per the above, the model also predicts that at matched levels of subjective visibility, perceptual evidence for unattended stimuli has similar overall magnitude to those of attended stimuli but is noisier, leading to lower signal-to-noise ratio for feature discrimination and thus poorer discrimination performance.

The dissociable mechanisms of overall evidence magnitude and signal-to-noise ratio in the model suggest an analogy with the dissociable subjective experiences of perceptual prominence and perceptual sharpness discussed above. Perceptual prominence maps naturally onto evidence magnitude, as both are magnitude-based concepts. Perceptual sharpness might also seem to map naturally onto signal-to-noise ratio as both are precision-based concepts, but further consideration raises some complexities. The signal detection model attributes feature detection reports to perceptual evidence exceeding a feature detection criterion, and so would seem to associate perceptual sharpness with evidence magnitude rather than signal-to-noise ratio per se. Additionally, signal-to-noise ratio in signal detection theory (SDT) pertains to signal precision across trials, whereas perceptual sharpness pertains to the precision of subjective experience for a single percept (a within-trial phenomenon), and SDT does not model within-trial noise.

Despite these conceptual and computational distinctions, it may be the case that within-trial perceptual sharpness and across-trial signal-to-noise ratio are closely related for suprathreshold stimuli, as we observed that in this regime reports of feature detection were closely correlated with objective feature discrimination performance in a way that was not modulated by attention (**Figure 3c**, middle panel). Ideally, a model of these phenomena would capture this empirical relationship while simultaneously resolving the associated conceptual and computational tensions noted above. For instance, such a model might link perceptual sharpness to within-trial uncertainty in perceptual processing, and in turn link this within-trial uncertainty to across-trial signal-to-noise ratio and thus objective discrimination performance. Alternatively, if one accepts that perceptual sharpness can be adequately characterized as a magnitude-based concept, or provides an account of how evidence magnitude and within-trial uncertainty are linked, then principles from the signal detection model of Rahnev et al.[6] might be sufficient to characterize our findings in the suprathreshold stimulus experiments without raising these conceptual tensions. One natural way to link evidence magnitude to within-trial uncertainty in SDT is via the likelihood ratio between the signal and noise distributions occurring at a given value of evidence magnitude, as evidence associated with a higher likelihood ratio entails better differentiation between signal and noise. However, the Rahnev et al.[6] model posits that decision criteria apply to raw evidence magnitude despite the likelihood ratio of these magnitudes differing across attention conditions, which raises complications for linking the two.

Although the modeling ideas discussed above may have intuitive appeal, a more definitive account would require extending any candidate model to simultaneously account for stimulus detection (related to “perceptual prominence”), feature detection (related to “perceptual sharpness”), and objective discrimination performance across all levels of attention and stimulus strength, and demonstrating that the model in question can account for the data well while also outperforming competing models.

**Supplementary Note 4**

*Inattentional inflation as asymmetry in stimulus-anchored attentional threshold shifts for performance vs. visibility*

#### Introduction

Previous treatments of inattentional inflation have focused on finding cases where point estimates of objective task performance[e] under attention and inattention are equal (e.g., by increasing stimulus strength for the unattended stimulus to compensate for the performance deficit induced by inattention), yet corresponding point estimates for subjective reports of visibility[f] are higher under inattention. More formally, inflation occurs when

$$\exists\, x_{\mathrm{A}}, x_{\mathrm{U}} \left[\, \left(P_{\mathrm{A}}(x_{\mathrm{A}}) = P_{\mathrm{U}}(x_{\mathrm{U}})\right) \wedge \left(V_{\mathrm{U}}(x_{\mathrm{U}}) > V_{\mathrm{A}}(x_{\mathrm{A}})\right) \,\right] \tag{S3}$$

for point estimates of task performance and visibility under attention and inattention $P_{\mathrm{A}}$, $V_{\mathrm{A}}$, $P_{\mathrm{U}}$, and $V_{\mathrm{U}}$ occurring at attended and unattended stimulus strengths $x_{\mathrm{A}}$ and $x_{\mathrm{U}}$.

Here we give broader theoretical consideration to the phenomenon of inattentional inflation in the context of a psychometric function analysis framework. This broader framing yields a new insight: given certain minimal assumptions, inattentional inflation occurs if and only if the effect of attention on psychometric function thresholds occurring at a particular unattended stimulus strength $x_{\mathrm{U}}$ is stronger for task performance than for visibility. This observation deepens our understanding of inattentional inflation—a phenomenon occurring at the level of the relative psychometric function relating visibility and performance—by showing how it manifests at the level of the conventional psychometric functions for visibility and performance.

#### Inattentional inflation in psychometric functions

Consider psychometric functions (PFs) $\psi$ with parameters $\theta$ for task performance *P* and visibility *V*, with separate functions for each depending on whether the stimulus is attended (A) or unattended (U):

$$\begin{aligned} P_{\mathrm{A}} &= \psi_{\mathrm{P\,|\,A}}(x; \theta_{\mathrm{P\,|\,A}}) \qquad P_{\mathrm{U}} = \psi_{\mathrm{P\,|\,U}}(x; \theta_{\mathrm{P\,|\,U}}) \\ V_{\mathrm{A}} &= \psi_{\mathrm{V\,|\,A}}(x; \theta_{\mathrm{V\,|\,A}}) \qquad V_{\mathrm{U}} = \psi_{\mathrm{V\,|\,U}}(x; \theta_{\mathrm{V\,|\,U}}) \end{aligned} \tag{S4}$$

where e.g., $P_{\mathrm{A}}$ is task performance under attention, $\psi_{\mathrm{P\,|\,A}}$ is the PF for task performance under attention, $\theta_{\mathrm{P\,|\,A}}$ is the set of parameters for $\psi_{\mathrm{P\,|\,A}}$, and *x* is stimulus strength.

[e] E.g., accuracy in a feature discrimination task.

[f] Inattentional inflation occurs for both subjective reports of visibility and estimates of confidence in one's perceptual decision. Here we discuss visibility for simplicity of exposition, but all arguments apply equally well for confidence.

From these conventional PFs, we may construct relative psychometric functions (RPFs) *R* to characterize visibility as a function of performance separately for each attention condition[9]:

$$V_{\mathrm{A}} = R_{\mathrm{VP}\,|\,\mathrm{A}}\left(P_{\mathrm{A}}\,;\,\theta_{\mathrm{P}\,|\,\mathrm{A}}, \theta_{\mathrm{V}\,|\,\mathrm{A}}\right)$$
$$V_{\mathrm{U}} = R_{\mathrm{VP}\,|\,\mathrm{U}}\left(P_{\mathrm{U}}\,;\,\theta_{\mathrm{P}\,|\,\mathrm{U}}, \theta_{\mathrm{V}\,|\,\mathrm{U}}\right) \tag{S5}$$

where e.g., $R_{\mathrm{VP}\,|\,\mathrm{A}}$ is the RPF relating visibility to task performance under attention. For brevity, we will not include notation for the PF or RPF parameters $\theta$ in the following, but they are implied.

We may formally characterize whether the RPFs in Eq. S5 exhibit inflation at a given performance level *P'* via

$$\text{inflation}(P') = \left[\, R_{\mathrm{VP}\,|\,\mathrm{U}}(P') > R_{\mathrm{VP}\,|\,\mathrm{A}}(P') \,\right] \quad \text{where} \quad P' \in \text{dom}\left(R_{\mathrm{VP}\,|\,\mathrm{A}}\right) \cap \text{dom}\left(R_{\mathrm{VP}\,|\,\mathrm{U}}\right) \tag{S6}$$

where inflation(*P'*) is a logical function defined for any *P'* that lies within the domain of both $R_{\mathrm{VP}\,|\,\mathrm{A}}$ and $R_{\mathrm{VP}\,|\,\mathrm{U}}$ that returns "true" if the condition for inattentional inflation is met at *P'* and "false" otherwise.[g]

Given that inattentional inflation is by definition an effect of attention on the visibility-performance RPF (Eqs. S5, S6), and that this RPF is constructed from conventional PFs for task performance and visibility (Eq. S4), it is natural to ask how inattentional inflation in the RPF manifests at the level of its constituent psychometric functions. A first move in this direction is to note that equating performance at *P'* across attentional conditions requires us to find the threshold stimulus strengths at which *P'* is achieved for both attentional conditions. Thresholds for performance *P'* and visibility *V'* for all PFs in Eq. S4 are given by

$$x_{P'\,|\,\mathrm{A}} = \psi^{-1}_{\mathrm{P}\,|\,\mathrm{A}}(P') \qquad x_{P'\,|\,\mathrm{U}} = \psi^{-1}_{\mathrm{P}\,|\,\mathrm{U}}(P')$$
$$x_{V'\,|\,\mathrm{A}} = \psi^{-1}_{\mathrm{V}\,|\,\mathrm{A}}(V') \qquad x_{V'\,|\,\mathrm{U}} = \psi^{-1}_{\mathrm{V}\,|\,\mathrm{U}}(V') \tag{S7}$$

where e.g., $x_{P'\,|\,\mathrm{A}}$ is the threshold value of *x* needed to achieve task performance *P'* under attention.

We may then re-express inattentional inflation (Eq. S6) at the level of PFs for visibility via

$$\text{inflation}(P') = \left[\psi_{\mathrm{V}\,|\,\mathrm{U}}\left(x_{P'\,|\,\mathrm{U}}\right) > \psi_{\mathrm{V}\,|\,\mathrm{A}}\left(x_{P'\,|\,\mathrm{A}}\right)\right] \quad \text{where} \quad P' \in \text{ran}\left(\psi_{\mathrm{P}\,|\,\mathrm{A}}\right) \cap \text{ran}\left(\psi_{\mathrm{P}\,|\,\mathrm{U}}\right) \tag{S8}$$

---

[g] Note that one way this condition can be undefined is if the asymptotic value of the PF for task performance is smaller under inattention than attention. This situation would yield a smaller domain of performance levels for the RPF under inattention, entailing that there are some *P'* values for which the RPF is defined under attention but not under inattention.

In other words, inflation occurs if the visibility of unattended stimuli at the performance threshold for *P′* (i.e., $x_{P' \mid \mathrm{U}}$) is greater than the visibility of attended stimuli at the same performance threshold ($x_{P' \mid \mathrm{A}}$). Note that the performance matching aspect of inflation is captured implicitly here by specifying stimulus strength *x* in terms of the threshold for achieving the same performance *P′* for each attentional condition.

Consideration of Eq. S8 suggests a connection between inattentional inflation and differential effects of attention on psychometric function thresholds for performance and visibility. The potential effect of attention on performance thresholds is implicit in Eq. S8 via the potential difference between the values for the threshold stimulus strengths at which performance *P′* is achieved under attention ($x_{P' \mid \mathrm{A}}$) and inattention ($x_{P' \mid \mathrm{U}}$). Furthermore, it is easy to see that if attention equally shifted the thresholds for the performance and visibility achieved at $x_{P' \mid \mathrm{U}}$, then visibility at $x_{P' \mid \mathrm{A}}$ and $x_{P' \mid \mathrm{U}}$ would be identical, and so inflation would not occur at *P′*. Therefore, for inflation to occur, there must be some asymmetry in how attention affects performance and visibility thresholds.

The main phenomenon of interest for characterizing inattentional inflation at the level of conventional psychometric functions is thus the *attentional threshold shift* occurring for some performance threshold for *P′* or visibility threshold for *V′*, given by

$$\Delta T_{\mathrm{P}}(P') = x_{P' \mid \mathrm{A}} - x_{P' \mid \mathrm{U}}$$
$$\Delta T_{\mathrm{V}}(V') = x_{V' \mid \mathrm{A}} - x_{V' \mid \mathrm{U}} \qquad \text{(S9)}$$

Negative attentional threshold shifts reflect improved performance or visibility with attention, because they indicate that the stimulus strength needed to achieve a given level of performance or visibility is lower for attended compared to unattended stimuli.

To facilitate direct comparison of the magnitudes of attentional threshold shifts for performance and visibility, it will be most useful to measure these shifts relative to the thresholds occurring for an unattended stimulus[h] of some given strength $x_{\mathrm{U}}$ (the *anchor stimulus*), via

$$\Delta T_{\mathrm{P}}^{\mathrm{anch}}(x_{\mathrm{U}}) = \psi_{\mathrm{P} \mid \mathrm{A}}^{-1}\left(\psi_{\mathrm{P} \mid \mathrm{U}}(x_{\mathrm{U}})\right) - x_{\mathrm{U}}$$
$$\Delta T_{\mathrm{V}}^{\mathrm{anch}}(x_{\mathrm{U}}) = \psi_{\mathrm{V} \mid \mathrm{A}}^{-1}\left(\psi_{\mathrm{V} \mid \mathrm{U}}(x_{\mathrm{U}})\right) - x_{\mathrm{U}} \qquad \text{(S10)}$$

[h] The order of subtraction in Eq. S9 is arbitrary, as is the choice in Eq. S10 of fixing comparison at attended or unattended stimulus strength. The choices we make here are intended to provide conceptual cohesion by always considering unattended stimulus strength as the implicit reference point in all equations such that these measures reflect attentional effects, i.e., effects due to the presence of attention relative to the absence of attention.

where $\Delta T_{\mathrm{P}}^{\mathrm{anch}}(x_{\mathrm{U}})$ and $\Delta T_{\mathrm{V}}^{\mathrm{anch}}(x_{\mathrm{U}})$ are the *stimulus-anchored attentional threshold shifts* for performance and visibility, respectively. Eq. S10 is equivalent to Eq. S9 when the *P′* and *V′* terms in Eq. S9 are defined to be the performance and visibility for unattended stimuli with strength $x_{\mathrm{U}}$ as specified in Eq. S10, i.e.

$$\Delta T_{\mathrm{P}}^{\mathrm{anch}}(x_{\mathrm{U}}) = \Delta T_{\mathrm{P}}(P') = x_{P'\,|\,\mathrm{A}} - x_{P'\,|\,\mathrm{U}} \quad \text{given} \quad P' = \psi_{\mathrm{P}\,|\,\mathrm{U}}(x_{\mathrm{U}})$$
$$\Delta T_{\mathrm{V}}^{\mathrm{anch}}(x_{\mathrm{U}}) = \Delta T_{\mathrm{V}}(V') = x_{V'\,|\,\mathrm{A}} - x_{V'\,|\,\mathrm{U}} \quad \text{given} \quad V' = \psi_{\mathrm{V}\,|\,\mathrm{U}}(x_{\mathrm{U}}) \tag{S11}$$

When evaluated at the same $x_{\mathrm{U}}$, comparing $\Delta T_{\mathrm{P}}^{\mathrm{anch}}(x_{\mathrm{U}})$ and $\Delta T_{\mathrm{V}}^{\mathrm{anch}}(x_{\mathrm{U}})$ can reveal differences in how attention affects the threshold stimulus strengths needed to maintain the same performance and visibility as are achieved for the unattended stimulus $x_{\mathrm{U}}$. Thus e.g., if $\Delta T_{\mathrm{P}}^{\mathrm{anch}}(x_{\mathrm{U}}) < \Delta T_{\mathrm{V}}^{\mathrm{anch}}(x_{\mathrm{U}})$, this implies that attention decreases the threshold for performance more than it does the threshold for visibility relative to the fixed unattended stimulus $x_{\mathrm{U}}$.[i]

Below, we prove that

$$\mathrm{inflation}(P') \leftrightarrow \left[\Delta T_{\mathrm{P}}^{\mathrm{anch}}\left(x_{P'\,|\,\mathrm{U}}\right) < \Delta T_{\mathrm{V}}^{\mathrm{anch}}\left(x_{P'\,|\,\mathrm{U}}\right)\right] \tag{S12}$$

provided some minimal assumptions hold. In other words, inattentional inflation occurs at performance level *P′* if and only if the stimulus-anchored attentional threshold shift $\Delta T^{\mathrm{anch}}$ is stronger (i.e., more negative) for task performance than for visibility at the value of $x_{\mathrm{U}}$ yielding *P′* for unattended stimuli. More loosely stated, inflation occurs at the level of the relative psychometric function when the effect of attention on conventional psychometric function thresholds is stronger for performance than for visibility.

In brief, the proof proceeds as follows:

1. Define *P′* and *V′* to be the performance and visibility occurring for some unattended "anchor" stimulus $x_{\mathrm{U}}$.
2. Assume that conditions hold which allow thresholds for *P′* and *V′* under attention and inattention (Eq. S7) to be well-defined.
3. Assume that the psychometric function for visibility under attention $\psi_{\mathrm{V}\,|\,\mathrm{A}}$ is a strictly increasing monotonic function over the interval of stimulus strengths containing the thresholds for *P′* and *V′*, i.e., $x_{P'\,|\,\mathrm{A}}$ and $x_{V'\,|\,\mathrm{A}}$.
4. Per Eq. S8, inflation occurs when visibility for the unattended stimulus yielding *P′* (i.e., $x_{P'\,|\,\mathrm{U}}$) is greater than the visibility for the attended stimulus yielding *P′* (i.e., $x_{P'\,|\,\mathrm{A}}$). Given the definition of *V′* in step 1, this is equivalent to stating that inflation occurs when visibility at $x_{V'\,|\,\mathrm{A}}$ exceeds visibility at $x_{P'\,|\,\mathrm{A}}$. Given the monotonicity assumption in step 3, this entails that $x_{P'\,|\,\mathrm{A}} < x_{V'\,|\,\mathrm{A}}$, i.e., that attention reduces the threshold for *P′* more than for *V′* relative

[i] We use language like "attention *decreases* the threshold for performance *more* than it does the threshold for visibility" for simplicity since we generally expect attention to decrease psychometric function thresholds, corresponding to negative values for both threshold shifts. However, technically speaking these threshold shifts need not be negative for the relationship in Eq. S12 to hold.

to the unattended anchor stimulus $x_{\mathrm{U}}$ defined in step 1. Thus inflation implies the presence of this asymmetric threshold shift.

5. Similarly, if we begin by assuming the presence of the asymmetric threshold shift relative to the anchor stimulus $x_{\mathrm{U}}$ such that $x_{P'|\mathrm{A}} < x_{V'|\mathrm{A}}$, the monotonicity assumption in step 3 entails that visibility is greater at $x_{V'|\mathrm{A}}$ than $x_{P'|\mathrm{A}}$, and the definitional equality of visibility at $x_{V'|\mathrm{A}}$ and $x_{P'|\mathrm{U}}$ entails greater visibility at $x_{P'|\mathrm{U}}$ than at $x_{P'|\mathrm{A}}$, i.e., inattentional inflation.

Essentially, the monotonicity assumption in step 3 is the bridge whereby the differences in visibility along the *y*-axis constitutive of inattentional inflation (Eq. S8) can be translated to the differences in threshold along the *x*-axis constitutive of asymmetry in stimulus-anchored attentional threshold shifts (Eq. S11), and vice versa. Via this link we can demonstrate the equivalence of the two phenomena (Eq. S12).

In **Figure S10** we provide a visual intuition for this connection. First we plot example RPFs exhibiting inattentional inflation at performance *P′* (**Figure S10a**; Eq. S6), and the corresponding conventional psychometric functions for visibility (**Figure S10b**; Eq. S8). In **Figure S10c** we highlight the effects of attention on psychometric function thresholds for performance and visibility. The crucial link between inflation and threshold shifts arises from how attention maps the visibility *V′* of the unattended anchor stimulus $x_{P'|\mathrm{U}}$ onto the attended psychometric function at $x_{V'|\mathrm{A}}$ via $\Delta T_{\mathrm{V}}^{\mathrm{anch}}$ (indicated in green). This converts the comparison of visibilities *across* attentional curves at $x_{P'|\mathrm{A}}$ and $x_{P'|\mathrm{U}}$ involved in assessing inattentional inflation (**Figure S10b**; Eq. S8) to a comparison of visibilities *within* the attended curve at $x_{P'|\mathrm{A}}$ and $x_{V'|\mathrm{A}}$ (**Figure S10c**). Given the assumption that attended visibility increases with stimulus strength, the only way for inflation to hold such that visibility at $x_{V'|\mathrm{A}}$ exceeds visibility at $x_{P'|\mathrm{A}}$ is if $x_{V'|\mathrm{A}} > x_{P'|\mathrm{A}}$. But in turn, this implies that $\Delta T_{\mathrm{V}}^{\mathrm{anch}} < \Delta T_{\mathrm{P}}^{\mathrm{anch}}$ (Eq. S11). In **Figure S10d** we illustrate that when $\Delta T_{\mathrm{V}}^{\mathrm{anch}} = \Delta T_{\mathrm{P}}^{\mathrm{anch}}$, visibility at *P′* must be equal across attentional conditions and so inflation at *P′* cannot occur.

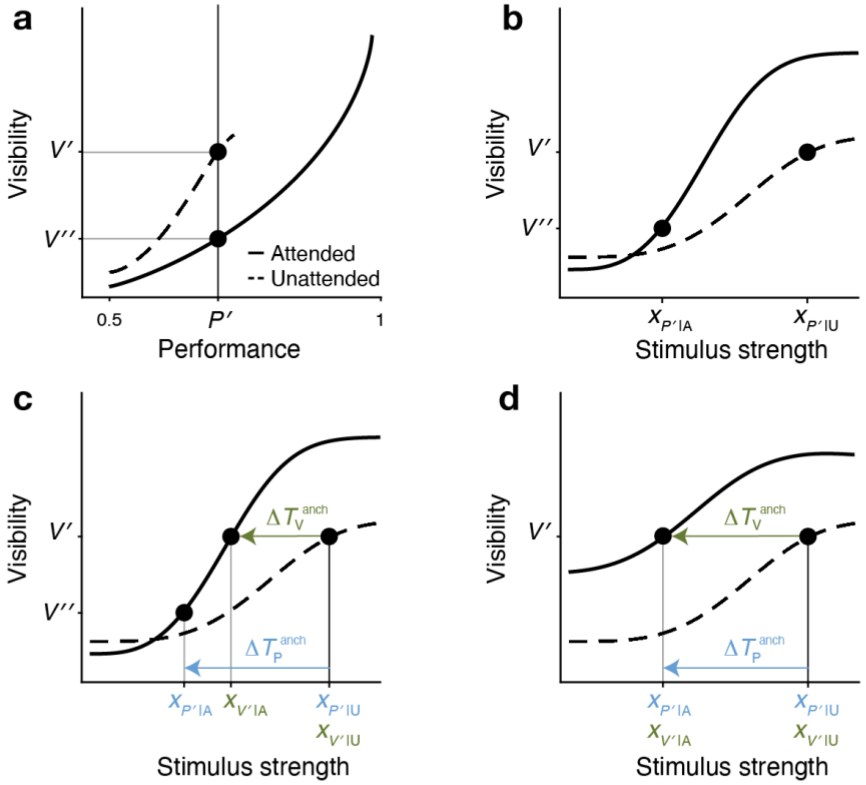


**Supplementary Figure 10. Visual intuition for the relationship between inattentional inflation and asymmetric threshold shifts for performance and visibility. a)** Inflation as expressed in relative psychometric functions, with example RPFs for visibility as a function of performance under attention and inattention (corresponding to $R_{\mathrm{VP\,|\,A}}$ and $R_{\mathrm{VP\,|\,U}}$ in Eqs. S5 and S6). Since visibility at performance *P′* is greater under inattention (*V′*) than under attention (*V″*), inattentional inflation occurs at *P′* (Eq. S6). **b)** Inflation as expressed in conventional psychometric functions. Here we plot the same visibility data as in (a), but plotted as a function of stimulus strength *x* rather than performance *P* (yielding $\psi_{\mathrm{V\,|\,A}}$ and $\psi_{\mathrm{V\,|\,U}}$ as in Eqs. S4 and S8). By definition, performance *P′* is achieved at threshold stimulus strengths $x_{P'\,|\,\mathrm{A}}$ under attention and $x_{P'\,|\,\mathrm{U}}$ under inattention (Eq. S7). Since inflation occurs at *P′*, visibility is greater at $x_{P'\,|\,\mathrm{U}}$ than at $x_{P'\,|\,\mathrm{A}}$ (Eq. S8). **c)** Inflation implies that attention reduces the visibility threshold less than the performance threshold. Here we plot the same data as in (b), now focusing on how attention influences psychophysical thresholds. In this example, attention reduces the threshold stimulus strengths needed to achieve the same performance *P′* and visibility *V′* as are achieved under inattention at $x_{P'\,|\,\mathrm{U}}$. The reduction in performance threshold $\Delta T_{\mathrm{P}}^{\mathrm{anch}}$ occurs as the leftward shift of $x_{P'\,|\,\mathrm{U}}$ to $x_{P'\,|\,\mathrm{A}}$ (Eq. S11), indicated in blue. The reduction in visibility threshold $\Delta T_{\mathrm{V}}^{\mathrm{anch}}$ occurs as the leftward shift of $x_{P'\,|\,\mathrm{U}}$ (equivalently, $x_{V'\,|\,\mathrm{U}}$) to $x_{V'\,|\,\mathrm{A}}$ (Eq. S11), indicated in green. The fact that the threshold shift is smaller for visibility than for performance is directly related to the fact that inflation occurs at *P′*. As shown in (a) and (b), inflation at *P′* means that unattended visibility *V′* at $x_{P'\,|\,\mathrm{U}}$ is greater than attended visibility *V″* at $x_{P'\,|\,\mathrm{A}}$ (Eq. S6). But this is equivalent to saying that *attended* visibility *V′* at $x_{V'\,|\,\mathrm{A}}$ is greater than attended visibility *V″* at $x_{P'\,|\,\mathrm{A}}$. Provided that attended visibility increases with stimulus strength, the only way for this to happen is if $x_{V'\,|\,\mathrm{A}} > x_{P'\,|\,\mathrm{A}}$, which in turn implies $\Delta T_{\mathrm{V}}^{\mathrm{anch}} < \Delta T_{\mathrm{P}}^{\mathrm{anch}}$ (Eq. S11). **d)** Inflation does not occur if attention does not reduce the visibility threshold less than the performance threshold. Here we plot the same unattended visibility data as in (c), but different attended visibility data, such that under attention, *V′* is now achieved at $x_{P'\,|\,\mathrm{A}}$. This entails that $\Delta T_{\mathrm{V}}^{\mathrm{anch}} = \Delta T_{\mathrm{P}}^{\mathrm{anch}}$. But since attended and unattended visibility are equal at $x_{P'\,|\,\mathrm{A}}$ and $x_{P'\,|\,\mathrm{U}}$, it follows that visibility is equal at *P′* and so inflation does not occur at *P′*. Similarly, if $\Delta T_{\mathrm{V}}^{\mathrm{anch}} > \Delta T_{\mathrm{P}}^{\mathrm{anch}}$ (not shown), inflation also does not occur (instead, the opposite happens—visibility at *P′* is greater for attention than for inattention). Thus, inflation at *P′* only occurs if $\Delta T_{\mathrm{V}}^{\mathrm{anch}} < \Delta T_{\mathrm{P}}^{\mathrm{anch}}$.

The equivalence of inflation and asymmetric threshold shifts suggests that at the descriptive level of analysis afforded by psychometric functions, these are two different ways of framing the same underlying phenomenon, with inflation framing things at the level of the relative psychometric functions relating visibility to performance (Eqs. S5 and S6) and the asymmetric threshold shift framing things at the level of the conventional psychometric functions for performance and visibility (Eqs. S4 and S8). Since these phenomena are equivalent at this descriptive level of analysis, neither one can be said to be more primary than the other, such that e.g., one arises from the other or occurs due to the other.

However, different theories may attribute greater primacy to underlying mechanisms more closely associated with one level of analysis than the other. For instance, a first-order theory might view inflation as being derivative upon the influence of attention on lower-level mechanisms determining performance and visibility thresholds. Conversely, a higher-order theory might view such threshold asymmetries as being derivative upon how attention changes the relationship between representations underlying task performance and higher-level representations underlying subjective visibility. These are empirical and theoretical questions beyond the scope of the analysis presented in this supplementary note.

#### Proof

##### Theorem: $\mathbf{inflation}(\boldsymbol{P'}) \leftrightarrow \left[\Delta \boldsymbol{T}_{\mathbf{P}}^{\mathbf{anch}}\left(\boldsymbol{x}_{\boldsymbol{P'}\,|\,\mathbf{U}}\right) < \Delta \boldsymbol{T}_{\mathbf{V}}^{\mathbf{anch}}\left(\boldsymbol{x}_{\boldsymbol{P'}\,|\,\mathbf{U}}\right)\right]$

The proof for Eq. S12 proceeds via separate proofs for each direction of implication. We will first list a set of definitions, assumptions, and corollaries common to both proofs before considering each separate proof in turn.

*Definitions common to both proofs*

**An unattended stimulus $x_{\mathrm{U}}$ yields *P′* and *V′*.** Let *P′* and *V′* be the performance and visibility yielded by a particular unattended "anchor" stimulus $x_{\mathrm{U}}$, i.e.,

$$P' = \psi_{\mathrm{P\,|\,U}}(x_{\mathrm{U}})$$
$$V' = \psi_{\mathrm{V\,|\,U}}(x_{\mathrm{U}}) \qquad \text{(D1)}$$

*Assumptions common to both proofs*

***P′* and *V′* can be achieved under attention.** The performance *P′* and visibility *V′* produced by the unattended stimulus $x_{\mathrm{U}}$ can also be produced by suitable attended stimuli, i.e.,

$$P' \in \mathrm{Range}\left(\psi_{\mathrm{P\,|\,A}}\right)$$
$$V' \in \mathrm{Range}\left(\psi_{\mathrm{V\,|\,A}}\right) \qquad \text{(A1)}$$

***P′* and *V′* have unique threshold stimulus strengths in all attentional conditions*.*** Under each attentional condition, there is a single stimulus strength yielding *P′*, and likewise for *V′*, such that

$$\exists!\, x \left[P' = \psi_{\mathrm{P}\,|\,\mathrm{A}}(x)\right] \qquad \exists!\, x \left[P' = \psi_{\mathrm{P}\,|\,\mathrm{U}}(x)\right]$$
$$\exists!\, x \left[V' = \psi_{\mathrm{V}\,|\,\mathrm{A}}(x)\right] \qquad \exists!\, x \left[V' = \psi_{\mathrm{V}\,|\,\mathrm{U}}(x)\right] \tag{A2}$$

where ∃! is the unique existential quantifier.

**Monotonicity over *P′* and *V′* thresholds under attention.** Over some interval of *x* values $I$ containing $x_{P'\,|\,\mathrm{A}}$ and $x_{V'\,|\,\mathrm{A}}$, $\psi_{\mathrm{V}\,|\,\mathrm{A}}$ is a strictly increasing monotonic function, such that

$$\forall\, x_1, x_2 \in I \left[x_2 > x_1 \rightarrow \psi_{\mathrm{V}\,|\,\mathrm{A}}(x_2) > \psi_{\mathrm{V}\,|\,\mathrm{A}}(x_1)\right] \tag{A3}$$

$$x_{P'\,|\,\mathrm{A}},\ x_{V'\,|\,\mathrm{A}} \in I \tag{A4}$$

**Comments**

Note that definition (D1) and assumptions (A1) and (A2) combine to ensure that the thresholds $x_{P'\,|\,\mathrm{A}}$, $x_{P'\,|\,\mathrm{U}}$, $x_{V'\,|\,\mathrm{A}}$, and $x_{V'\,|\,\mathrm{U}}$ in Eq. S7 all have unique, well-defined values. In order for well-defined thresholds to exist, *P′* or *V′* must lie within the range of the relevant psychometric function, which is ensured under inattention by definition (D1) and under attention by assumption (A1). Additionally, for the inverse of the psychometric function to be defined for *P′* or *V′* as in Eq. S7, it must be the case that *P′* or *V′* can only be produced by a single *x* value within a given attentional condition; this stipulation fails to hold e.g., if the psychometric function has a constant value of *P′* or *V′* over an interval of *x* values, or if the function produces multiple instances of *P′* or *V′* due to non-monotonicity. Uniqueness of thresholds is ensured by assumption (A2).

The simplest way to ensure that (A2) holds is to assume that all psychometric functions under consideration are strictly increasing monotonic functions (and are therefore invertible) over all possible *x* values. This would in turn render assumption (A3) redundant. However, psychometric functions are typically sigmoidal with the possibility of nearly flat asymptotes at very low or high stimulus strengths. Although such asymptotes can be formally characterized as strictly increasing, for practical purposes they are essentially constant, such that it is not possible to compute a unique and stable threshold estimate very close to the asymptotic value. A weaker and more generally applicable assumption is that psychometric functions exhibit non-decreasing monotonicity rather than strictly increasing monotonicity. Thus, we could instead assume that all psychometric functions under consideration exhibit non-decreasing monotonicity, and further assume that these exhibit strictly increasing monotonicity over subintervals of each function in a way that satisfies the conditions stipulated in the current (A2) and (A3), but it becomes more difficult to characterize this set of assumptions clearly and concisely. Therefore we use the current formulation to strike the best balance between generality, practical applicability, and simplicity.

**Corollaries**

From definition (D1) and assumption (A2), we can use the terms defined in Eq. S7 to write that

$$x_{\mathrm{U}} = x_{P' \mid \mathrm{U}} = x_{V' \mid \mathrm{U}} \tag{C1}$$

$$V' = \psi_{\mathrm{V} \mid \mathrm{U}}\left(x_{P' \mid \mathrm{U}}\right) \tag{C2}$$

Given assumptions (A1) and (A2) and the definitions in Eq. S7, we can further express *V'* in terms of the stimulus strength $x_{V' \mid \mathrm{A}}$ needed to achieve this visibility under attention:

$$V' = \psi_{V \mid \mathrm{A}}\left(x_{V' \mid \mathrm{A}}\right) \tag{C3}$$

Combining corollaries (C2) and (C3) gives

$$\psi_{\mathrm{V} \mid \mathrm{U}}\left(x_{P' \mid \mathrm{U}}\right) = \psi_{\mathrm{V} \mid \mathrm{A}}\left(x_{V' \mid \mathrm{A}}\right) \tag{C4}$$

It follows straightforwardly from the monotonicity assumption in (A3) that

$$\forall\, x_1, x_2 \in I\ \left[\psi_{\mathrm{V} \mid \mathrm{A}}(x_2) > \psi_{\mathrm{V} \mid \mathrm{A}}(x_1) \rightarrow x_2 > x_1\right] \tag{C5}$$

#### Theorem 1: $\mathbf{inflation}(\boldsymbol{P'}) \rightarrow \left[\boldsymbol{\Delta T}_{\mathbf{P}}^{\mathbf{anch}}\left(\boldsymbol{x}_{\boldsymbol{P'} \mid \mathbf{U}}\right) < \boldsymbol{\Delta T}_{\mathbf{V}}^{\mathbf{anch}}\left(\boldsymbol{x}_{\boldsymbol{P'} \mid \mathbf{U}}\right)\right]$

*Assumption unique to this proof*

**Inflation at *P'*.** Inattentional inflation occurs at *P'*, such that following Eq. S8, we can write

$$\psi_{\mathrm{V} \mid \mathrm{U}}\left(x_{P' \mid \mathrm{U}}\right) > \psi_{\mathrm{V} \mid \mathrm{A}}\left(x_{P' \mid \mathrm{A}}\right) \tag{P1.A1}$$

**Proof 1**

Combining assumption (P1.A1) on inattentional inflation with corollary (C4) and reversing the order of the inequality gives

$$\psi_{\mathrm{V\,|\,A}}(x_{P'\,|\,\mathrm{A}}) < \psi_{\mathrm{V\,|\,A}}(x_{V'\,|\,\mathrm{A}}) \qquad \text{(P1.1)}$$

Applying monotonicity-related statements (C5) and (A4) to (P1.1) entails that

$$x_{P'\,|\,\mathrm{A}} < x_{V'\,|\,\mathrm{A}} \qquad \text{(P1.2)}$$

From (C1), (P1.2), and Eq. S11, it follows that the stimulus-anchored attentional threshold shift for task performance (i.e., $\Delta T_{\mathrm{P}}^{\mathrm{anch}}$) for the unattended stimulus strength yielding *P′* (i.e., $x_{P'\,|\,\mathrm{U}}$) must be lower than the corresponding threshold shift for visibility (i.e., $\Delta T_{\mathrm{V}}^{\mathrm{anch}}$):

$$\Delta T_{\mathrm{P}}^{\mathrm{anch}}(x_{P'\,|\,\mathrm{U}}) \; < \; \Delta T_{\mathrm{V}}^{\mathrm{anch}}(x_{P'\,|\,\mathrm{U}}) \qquad \text{(P1.3)}$$

■

#### Theorem 2: $\left[\boldsymbol{\Delta T_{\mathrm{P}}^{\mathrm{anch}}(x_{P'\,|\,\mathrm{U}}) \; < \; \Delta T_{\mathrm{V}}^{\mathrm{anch}}(x_{P'\,|\,\mathrm{U}})}\right] \rightarrow \textbf{inflation}(\boldsymbol{P'})$

*Assumption unique to this proof*

**Asymmetric stimulus-anchored attentional threshold shifts at $\boldsymbol{x_{P'\,|\,\mathrm{U}}}$.** The stimulus-anchored attentional threshold shift at $x_{P'\,|\,\mathrm{U}}$ is lower for performance than for visibility, such that

$$\Delta T_{\mathrm{P}}^{\mathrm{anch}}(x_{P'\,|\,\mathrm{U}}) \; < \; \Delta T_{\mathrm{V}}^{\mathrm{anch}}(x_{P'\,|\,\mathrm{U}}) \qquad \text{(P2.A1)}$$

**Proof 2**

From corollary (C1) and Eq. S11, it follows that the only way for the inequality of threshold shifts in assumption (P2.A1) to hold is if

$$x_{P'\,|\,\mathrm{A}} < x_{V'\,|\,\mathrm{A}} \qquad \text{(P2.1)}$$

From (P2.1) and monotonicity-related assumptions (A3) and (A4), it follows that

$$\psi_{\mathrm{V}\,|\,\mathrm{A}}(x_{P'\,|\,\mathrm{A}}) < \psi_{\mathrm{V}\,|\,\mathrm{A}}(x_{V'\,|\,\mathrm{A}}) \qquad \text{(P2.2)}$$

Combining (P2.2) with (C4) and reversing the order of the inequality reveals inattentional inflation at *P′* (see Eq. S8):

$$\psi_{\mathrm{V}\,|\,\mathrm{U}}(x_{P'\,|\,\mathrm{U}}) > \psi_{\mathrm{V}\,|\,\mathrm{A}}(x_{P'\,|\,\mathrm{A}}) \qquad \text{(P2.3)}$$

■

#### Supplementary Methods

#### Task instruction excerpts

##### Experiment 1: Threshold detection of gratings

In this experiment you will be asked to make judgments about visual stimuli. Specifically, you will be pointed or “cued” to a particular location on the screen and will have to respond:

1. Did you see a grating embedded in the noise patch in the cued location?
2. If yes, did you see what direction the grating was oriented?
3. Was the grating oriented counterclockwise (-45 deg) or clockwise (+45 deg) from vertical?

We ask separate questions about whether you saw a grating and whether you saw its orientation because sometimes you may clearly see some kind of grating in the noise, without being able to see clearly the exact orientation of this grating.

About the “saw grating” / “saw orientation” judgments

> When we ask about whether you saw a grating and its orientation, we’re interested to know about what your actual visual experience was like. Did it actually look like there was a grating embedded in the noise patch in the response-cued location? If so, could you actually see whether the grating was oriented counterclockwise or clockwise from vertical?
>
> Don’t try to answer this question based on any information other than what your actual visual experience of the patch was like.

About the “counterclockwise” / “clockwise” judgment:

> Sometimes you may not be very sure whether the grating was oriented counterclockwise or clockwise from vertical. This may be especially the case when you didn’t see a grating to begin with!
>
> On trials where the response-cued quadrant didn’t contain a grating to begin with, obviously the counterclockwise / clockwise judgment has no meaning. However, just because you didn’t see a grating doesn’t mean there wasn’t one there! In cases where a grating was objectively present but you didn’t see it, the orientation judgment is still meaningful and your response may be meaningful too, even if it “feels” like a wild guess.
>
> For this reason, we ask that you take the counterclockwise / clockwise question seriously on each trial and give the best response you can, even when you didn’t see a grating. On such trials, try to make the orientation judgment *as if* a grating had been presented but you just didn't see it. If you’re not sure about the grating’s orientation, just make your best gut instinct guess without spending too much time deliberating.
>
> If you have to guess, it’s important that you don’t guess based on a strategy (like always responding the opposite of what you chose last, or always responding “counterclockwise”). Try to keep your “counterclockwise” and “clockwise” guesses roughly balanced, based on your best hunch. If you have no idea at all, try to pick randomly, as if you were flipping a coin.

Stimulus frequencies and dependencies

> Overall, each quadrant is equally likely to contain a grating. When a grating is present, it is equally likely to be oriented counterclockwise or clockwise from vertical.
>
> It is very important to note that grating presence and orientation in each quadrant is completely independent of grating presence and orientation in the other quadrants. In other words, knowing what was shown in one quadrant gives you no information whatsoever about what was shown in any other quadrant.

As a consequence, your responses should always be based *only* on what you saw at the response-cued quadrant, and should never be influenced by what you saw at any of the quadrants that were not cued.

#### Supplementary Tables

**Supplementary Table 1.** Objective performance (related to Figure 2a).

| | Effect | $DF_n$ | $DF_d$ | *F* | *p* | $\eta_G^2$ | $\varepsilon$ | *p*[GG] |
|---|---|---|---|---|---|---|---|---|
| | *All experiments* | | | | | | | |
| 1 | Site | 1 | 111 | 5.87 | **0.017** | 0.02 | - | - |
| 2 | Exp | 3 | 111 | 2.69 | 0.056 | 0.03 | - | - |
| 3 | Strength | 6 | 666 | 748.05 | **<0.001** | 0.61 | 0.48 | **<0.001** |
| 4 | Att | 2 | 222 | 892.80 | **<0.001** | 0.52 | 0.70 | **<0.001** |
| 5 | Site:Exp | 3 | 111 | 2.67 | 0.051 | 0.03 | - | - |
| 6 | Site:Strength | 6 | 666 | 0.18 | 0.983 | <0.01 | 0.48 | 0.907 |
| 7 | Exp:Strength | 18 | 666 | 13.79 | **<0.001** | 0.08 | 0.48 | **<0.001** |
| 8 | Site:Att | 2 | 222 | 2.78 | 0.064 | <0.01 | 0.70 | 0.084 |
| 9 | Exp:Att | 6 | 222 | 14.78 | **<0.001** | 0.05 | 0.70 | **<0.001** |
| 10 | Strength:Att | 12 | 1332 | 53.93 | **<0.001** | 0.08 | 0.75 | **<0.001** |
| 11 | Site:Exp:Strength | 18 | 666 | 0.93 | 0.538 | 0.01 | 0.48 | 0.495 |
| 12 | Site:Exp:Att | 6 | 222 | 0.48 | 0.820 | <0.01 | 0.70 | 0.756 |
| 13 | Site:Strength:Att | 12 | 1332 | 2.05 | **0.018** | <0.01 | 0.75 | **0.032** |
| 14 | Exp:Strength:Att | 36 | 1332 | 4.45 | **<0.001** | 0.02 | 0.75 | **<0.001** |
| 15 | Site:Exp:Strength:Att | 36 | 1332 | 1.78 | **0.003** | 0.01 | 0.75 | **0.009** |
| | *Experiment 1* | | | | | | | |
| 1 | Site | 1 | 28 | 0.17 | 0.681 | <0.01 | - | - |
| 2 | Strength | 6 | 168 | 158.04 | **<0.001** | 0.62 | 0.56 | **<0.001** |
| 3 | Att | 2 | 56 | 315.97 | **<0.001** | 0.64 | 0.69 | **<0.001** |
| 4 | Site:Strength | 6 | 168 | 1.01 | 0.418 | 0.01 | 0.56 | 0.396 |
| 5 | Site:Att | 2 | 56 | 1.65 | 0.202 | <0.01 | 0.69 | 0.210 |
| 6 | Strength:Att | 12 | 336 | 18.56 | **<0.001** | 0.16 | 0.61 | **<0.001** |
| 7 | Site:Strength:Att | 12 | 336 | 1.54 | 0.109 | 0.02 | 0.61 | 0.154 |
| | *Experiment 2* | | | | | | | |
| 1 | Site | 1 | 28 | 11.39 | **0.002** | 0.11 | - | - |
| 2 | Strength | 6 | 168 | 249.79 | **<0.001** | 0.72 | 0.30 | **<0.001** |
| 3 | Att | 2 | 56 | 356.47 | **<0.001** | 0.70 | 0.63 | **<0.001** |
| 4 | Site:Strength | 6 | 168 | 0.41 | 0.873 | <0.01 | 0.30 | 0.645 |
| 5 | Site:Att | 2 | 56 | 2.03 | 0.142 | 0.01 | 0.69 | 0.158 |
| 6 | Strength:Att | 12 | 336 | 30.10 | **<0.001** | 0.19 | 0.55 | **<0.001** |
| 7 | Site:Strength:Att | 12 | 336 | 3.35 | **<0.001** | 0.03 | 055 | **0.003** |
| | *Experiment 3* | | | | | | | |
| 1 | Site | 1 | 28 | 0.60 | 0.445 | <0.01 | - | - |
| 2 | Strength | 6 | 168 | 219.99 | **<0.001** | 0.74 | 0.38 | **<0.001** |
| 3 | Att | 2 | 56 | 112.19 | **<0.001** | 0.37 | 0.62 | **<0.001** |
| 4 | Site:Strength | 6 | 168 | 0.80 | 0.573 | 0.10 | 0.38 | 0.469 |
| 5 | Site:Att | 2 | 56 | 0.73 | 0.487 | <0.01 | 0.62 | 0.427 |
| 6 | Strength:Att | 12 | 336 | 12.90 | **<0.001** | 0.01 | 0.62 | **<0.001** |
| 7 | Site:Strength:Att | 12 | 336 | 1.48 | 0.131 | 0.01 | 0.62 | 0.173 |
| | *Experiment 4* | | | | | | | |
| 1 | Site | 1 | 28 | 3.30 | 0.080 | 0.08 | - | - |
| 2 | Strength | 6 | 168 | 148.88 | **<0.001** | 0.33 | 0.39 | **<0.001** |
| 3 | Att | 2 | 56 | 173.33 | **<0.001** | 0.38 | 0.75 | **<0.001** |
| 4 | Site:Strength | 6 | 168 | 0.59 | 0.741 | <0.01 | 0.39 | 0.587 |
| 5 | Site:Att | 2 | 56 | 0.05 | 0.948 | <0.01 | 0.75 | 0.904 |
| 6 | Strength:Att | 12 | 336 | 7.80 | **<0.001** | 0.03 | 0.51 | **<0.001** |
| 7 | Site:Strength:Att | 12 | 336 | 1.58 | 0.095 | <0.01 | 0.51 | 0.154 |

Factors of experiment, attention, and stimulus strength are abbreviated as “Exp,” “Att,” and “Strength.” The statistical tests used in this table were repeated measures ANOVAs. For all ANOVA tables, when Mauchly’s test indicated violation of sphericity, Greenhouse-Geisser epsilon and corrected p-values are shown. Significant p-values are bolded.

**Supplementary Table 2.** Objective performance for stimuli reported as "unseen" (related to Supplementary Figure 3).

| | Effect | $DF_n$ | $DF_d$ | *F* | p | $\eta^2_G$ | $\varepsilon$ | p[GG] |
|---|---|---|---|---|---|---|---|---|
| | *Experiments 1 and 3* | | | | | | | |
| 1 | Site | 1 | 56 | 1.67 | 0.201 | 0.02 | - | - |
| 2 | Expt | 1 | 56 | 0.05 | 0.817 | <0.01 | - | - |
| 3 | Att | 2 | 112 | 5.48 | **0.005** | 0.03 | 0.94 | **0.006** |
| 4 | Site:Expt | 1 | 56 | 0.05 | 0.818 | <0.01 | - | - |
| 5 | Site:Att | 2 | 112 | 2.27 | 0.108 | 0.01 | 0.94 | 0.112 |
| 6 | Expt:Att | 2 | 112 | 0.41 | 0.664 | <0.01 | 0.94 | 0.651 |
| 7 | Site:Expt:Att | 2 | 112 | 2.72 | 0.070 | 0.02 | 0.94 | 0.074 |

**Supplementary Table 3.** Subjective reports of stimulus visibility (related to Figure 2b).

| | Effect | $DF_n$ | $DF_d$ | *F* | *p* | $\eta_G^2$ | $\varepsilon$ | *p*[GG] |
|---|---|---|---|---|---|---|---|---|
| | *All experiments* | | | | | | | |
| 1 | Site | 1 | 111 | 0.13 | 0.720 | <0.01 | - | - |
| 2 | Exp | 3 | 111 | 87.35 | **<0.001** | 0.52 | - | - |
| 3 | Strength | 6 | 666 | 544.71 | **<0.001** | 0.65 | 0.34 | **<0.001** |
| 4 | Att | 2 | 222 | 82.33 | **<0.001** | 0.05 | 0.70 | **<0.001** |
| 5 | Site:Exp | 3 | 111 | 0.19 | 0.902 | <0.01 | - | - |
| 6 | Site:Strength | 6 | 666 | 0.57 | 0.753 | <0.01 | 0.34 | 0.570 |
| 7 | Exp:Strength | 18 | 666 | 34.72 | **<0.001** | 0.26 | 0.34 | **<0.001** |
| 8 | Site:Att | 2 | 222 | 2.57 | 0.079 | <0.01 | 0.70 | 0.099 |
| 9 | Exp:Att | 6 | 222 | 25.78 | **<0.001** | 0.05 | 0.70 | **<0.001** |
| 10 | Strength:Att | 12 | 1332 | 67.52 | **<0.001** | 0.05 | 0.53 | **<0.001** |
| 11 | Site:Exp:Strength | 18 | 666 | 1.51 | 0.081 | 0.02 | 0.34 | 0.175 |
| 12 | Site:Exp:Att | 6 | 222 | 3.22 | **0.005** | <0.01 | 0.70 | **0.013** |
| 13 | Site:Strength:Att | 12 | 1332 | 2.07 | 0.016 | <0.01 | 0.53 | 0.050 |
| 14 | Exp:Strength:Att | 36 | 1332 | 13.93 | **<0.001** | 0.03 | 0.53 | **<0.001** |
| 15 | Site:Exp:Strength:Att | 36 | 1332 | 0.75 | 0.859 | <0.01 | 0.53 | 0.769 |
| | *Experiment 1* | | | | | | | |
| 1 | Site | 1 | 28 | 0.02 | 0.889 | <0.01 | - | - |
| 2 | Strength | 6 | 168 | 168.54 | **<0.001** | 0.57 | 0.31 | **<0.001** |
| 3 | Att | 2 | 56 | 98.32 | **<0.001** | 0.23 | 0.61 | **<0.001** |
| 4 | Site:Strength | 6 | 168 | 0.11 | 0.998 | <0.01 | 0.31 | 0.883 |
| 5 | Site:Att | 2 | 56 | 8.13 | **0.001** | 0.02 | 0.61 | **0.005** |
| 6 | Strength:Att | 12 | 336 | 24.87 | **<0.001** | 0.07 | 0.50 | **<0.001** |
| 7 | Site:Strength:Att | 12 | 336 | 1.02 | 0.434 | <0.01 | 0.50 | 0.416 |
| | *Experiment 2* | | | | | | | |
| 1 | Site | 1 | 28 | 0.84 | 0.367 | 0.01 | - | - |
| 2 | Strength | 6 | 168 | 220.87 | **<0.001** | 0.81 | 0.28 | **<0.001** |
| 3 | Att | 2 | 56 | 20.41 | **<0.001** | 0.02 | 0.81 | **<0.001** |
| 4 | Site:Strength | 6 | 168 | 0.20 | 0.977 | <0.01 | 0.28 | 0.783 |
| 5 | Site:Att | 2 | 56 | 2.32 | 0.108 | <0.01 | 0.81 | 0.119 |
| 6 | Strength:Att | 12 | 336 | 22.76 | **<0.001** | 0.08 | 0.26 | **<0.001** |
| 7 | Site:Strength:Att | 12 | 336 | 0.55 | 0.885 | <0.01 | 0.26 | 0.660 |
| | *Experiment 3* | | | | | | | |
| 1 | Site | 1 | 28 | 0.06 | 0.807 | <0.01 | - | - |
| 2 | Strength | 6 | 168 | 171.40 | **<0.001** | 0.73 | 0.37 | **<0.001** |
| 3 | Att | 2 | 56 | 22.98 | **<0.001** | 0.08 | 0.59 | **<0.001** |
| 4 | Site:Strength | 6 | 168 | 1.34 | 0.240 | 0.02 | 0.37 | 0.269 |
| 5 | Site:Att | 2 | 56 | 0.90 | 0.414 | <0.01 | 0.59 | 0.368 |
| 6 | Strength:Att | 12 | 336 | 17.98 | **<0.001** | 0.06 | 0.43 | **<0.001** |
| 7 | Site:Strength:Att | 12 | 336 | 1.52 | 0.115 | <0.01 | 0.43 | 0.185 |
| | *Experiment 4* | | | | | | | |
| 1 | Site | 1 | 28 | 0.13 | 0.721 | <0.01 | - | - |
| 2 | Strength | 6 | 168 | 102.84 | **<0.001** | 0.60 | 0.23 | **<0.001** |
| 3 | Att | 2 | 56 | 2.29 | 0.111 | 0.01 | 0.79 | 0.124 |
| 4 | Site:Strength | 6 | 168 | 2.65 | 0.018 | 0.04 | 0.23 | 0.102 |
| 5 | Site:Att | 2 | 56 | 0.18 | 0.834 | <0.01 | 0.79 | 0.782 |
| 6 | Strength:Att | 12 | 336 | 46.17 | **<0.001** | 0.09 | 0.38 | **<0.001** |
| 7 | Site:Strength:Att | 12 | 336 | 1.33 | 0.199 | <0.01 | 0.38 | 0.259 |

**Supplementary Table 4.** Subjective reports of task-relevant feature visibility (related to Figure 3b).

| | Effect | $DF_n$ | $DF_d$ | *F* | *p* | $\eta_G^2$ | $\varepsilon$ | *p*[GG] |
|---|---|---|---|---|---|---|---|---|
| | *All experiments* | | | | | | | |
| 1 | Site | 1 | 84 | 0.01 | 0.915 | <0.01 | - | - |
| 2 | Exp | 2 | 84 | 11.92 | **<0.001** | 0.15 | - | - |
| 3 | Strength | 6 | 504 | 457.96 | **<0.001** | 0.53 | 0.36 | **<0.001** |
| 4 | Att | 2 | 168 | 327.53 | **<0.001** | 0.27 | 0.63 | **<0.001** |
| 5 | Site:Exp | 2 | 84 | 0.47 | 0.626 | 0.01 | - | - |
| 6 | Site:Strength | 6 | 504 | 0.37 | 0.888 | <0.01 | 0.36 | 0.699 |
| 7 | Exp:Strength | 12 | 504 | 7.15 | **<0.001** | 0.03 | 0.36 | **<0.001** |
| 8 | Site:Att | 2 | 168 | 0.50 | 0.606 | <0.01 | 0.63 | 0.523 |
| 9 | Exp:Att | 4 | 168 | 13.83 | **<0.001** | 0.03 | 0.63 | **<0.001** |
| 10 | Strength:Att | 12 | 1008 | 71.71 | **<0.001** | 0.06 | 0.48 | **<0.001** |
| 11 | Site:Exp:Strength | 12 | 504 | 0.89 | 0.561 | <0.01 | 0.36 | 0.480 |
| 12 | Site:Exp:Att | 4 | 168 | 0.52 | 0.724 | <0.01 | 0.63 | 0.641 |
| 13 | Site:Strength:Att | 12 | 1008 | 0.80 | 0.654 | <0.01 | 0.48 | 0.567 |
| 14 | Exp:Strength:Att | 24 | 1008 | 7.81 | **<0.001** | 0.01 | 0.48 | **<0.001** |
| 15 | Site:Exp:Strength:Att | 24 | 1008 | 1.35 | 0.120 | <0.01 | 0.48 | 0.189 |
| | *Experiment 1* | | | | | | | |
| 1 | Site | 1 | 28 | 0.56 | 0.461 | 0.83 | - | - |
| 2 | Strength | 6 | 168 | 209.62 | **<0.001** | 0.57 | 0.35 | **<0.001** |
| 3 | Att | 2 | 56 | 136.45 | **<0.001** | 0.27 | 0.67 | **<0.001** |
| 4 | Site:Strength | 6 | 168 | 1.07 | 0.385 | <0.01 | 0.35 | 0.353 |
| 5 | Site:Att | 2 | 56 | 2.12 | 0.130 | <0.01 | 0.67 | 0.148 |
| 6 | Strength:Att | 12 | 336 | 37.09 | **<0.001** | 0.01 | 0.45 | **<0.001** |
| 7 | Site:Strength:Att | 12 | 336 | 1.19 | 0.287 | <0.01 | 0.45 | 0.315 |
| | *Experiment 2* | | | | | | | |
| 1 | Site | 1 | 28 | 0.19 | 0.669 | <0.01 | - | - |
| 2 | Strength | 6 | 168 | 116.73 | **<0.001** | 0.41 | 0.25 | **<0.001** |
| 3 | Att | 2 | 56 | 116.52 | **<0.001** | 0.39 | 0.59 | **<0.001** |
| 4 | Site:Strength | 6 | 168 | 0.64 | 0.702 | <0.01 | 0.25 | 0.493 |
| 5 | Site:Att | 2 | 56 | 0.01 | 0.989 | <0.01 | 0.59 | 0.941 |
| 6 | Strength:Att | 12 | 336 | 21.46 | **<0.001** | 0.05 | 0.26 | **<0.001** |
| 7 | Site:Strength:Att | 12 | 336 | 0.98 | 0.468 | <0.01 | 0.26 | 0.408 |
| | *Experiment 3* | | | | | | | |
| 1 | Site | 1 | 28 | 0.23 | 0.637 | <0.01 | - | - |
| 2 | Strength | 6 | 168 | 153.98 | **<0.001** | 0.60 | 0.38 | **<0.001** |
| 3 | Att | 2 | 56 | 101.66 | **<0.001** | 0.16 | 0.67 | **<0.001** |
| 4 | Site:Strength | 6 | 168 | 0.57 | 0.751 | <0.01 | 0.38 | 0.590 |
| 5 | Site:Att | 2 | 56 | 0.03 | 0.972 | <0.01 | 0.67 | 0.922 |
| 6 | Strength:Att | 12 | 336 | 29.59 | **<0.001** | 0.06 | 0.59 | **<0.001** |
| 7 | Site:Strength:Att | 12 | 336 | 1.37 | 0.178 | <0.01 | 0.59 | 0.219 |

**Supplementary Table 5.** Stimulus detection sensitivity (related to Supplementary Figure 5a).

| | Effect | $DF_n$ | $DF_d$ | *F* | *p* | $\eta_G^2$ | $\varepsilon$ | *p*[GG] |
|---|---|---|---|---|---|---|---|---|
| | *All experiments* | | | | | | | |
| 1 | Site | 1 | 56 | 1.40 | 0.241 | 0.01 | - | - |
| 2 | Exp | 1 | 56 | 2.81 | 0.099 | 0.03 | - | - |
| 3 | Strength | 6 | 336 | 350.59 | **<0.001** | 0.59 | 0.42 | **<0.001** |
| 4 | Att | 2 | 112 | 296.54 | **<0.001** | 0.40 | 0.74 | **<0.001** |
| 5 | Site:Exp | 1 | 56 | <0.01 | 0.944 | <0.01 | - | - |
| 6 | Site:Strength | 6 | 336 | 0.44 | 0.854 | <0.01 | 0.42 | 0.693 |
| 7 | Exp:Strength | 6 | 336 | 10.97 | **<0.001** | 0.04 | 0.42 | **<0.001** |
| 8 | Site:Att | 2 | 112 | 0.35 | 0.703 | <0.01 | 0.74 | 0.639 |
| 9 | Exp:Att | 2 | 112 | 3.13 | 0.048 | <0.01 | 0.74 | 0.063 |
| 10 | Strength:Att | 12 | 672 | 56.13 | **<0.001** | 0.10 | - | - |
| 11 | Site:Exp:Strength | 6 | 336 | 0.59 | 0.739 | <0.01 | 0.42 | 0.594 |
| 12 | Site:Exp:Att | 2 | 112 | 0.32 | 0.725 | <0.01 | 0.74 | 0.660 |
| 13 | Site:Strength:Att | 12 | 672 | 1.39 | 0.164 | <0.01 | - | - |
| 14 | Exp:Strength:Att | 12 | 672 | 1.32 | 0.201 | <0.01 | - | - |
| 15 | Site:Exp:Strength:Att | 12 | 672 | 2.30 | **0.007** | <0.01 | - | - |
| | *Experiment 1* | | | | | | | |
| 1 | Site | 1 | 28 | 0.77 | 0.389 | 0.02 | - | - |
| 2 | Strength | 6 | 168 | 243.16 | **<0.001** | 0.60 | 0.38 | **<0.001** |
| 3 | Att | 2 | 56 | 186.97 | **<0.001** | 0.48 | 0.80 | **<0.001** |
| 4 | Site:Strength | 6 | 168 | 0.06 | 0.999 | <0.01 | 0.38 | 0.956 |
| 5 | Site:Att | 2 | 56 | 0.67 | 0.517 | <0.01 | 0.80 | 0.485 |
| 6 | Strength:Att | 12 | 336 | 45.43 | **<0.001** | 0.11 | 0.61 | **<0.001** |
| 7 | Site:Strength:Att | 12 | 336 | 1.15 | 0.319 | <0.01 | 0.61 | 0.333 |
| | *Experiment 3* | | | | | | | |
| 1 | Site | 1 | 28 | 0.64 | 0.431 | 0.01 | - | - |
| 2 | Strength | 6 | 168 | 149.19 | **<0.001** | 0.59 | 0.40 | **<0.001** |
| 3 | Att | 2 | 56 | 115.62 | **<0.001** | 0.33 | 0.66 | **<0.001** |
| 4 | Site:Strength | 6 | 168 | 0.74 | 0.616 | <0.01 | 0.40 | 0.503 |
| 5 | Site:Att | 2 | 56 | 0.03 | 0.966 | <0.01 | 0.66 | 0.910 |
| 6 | Strength:Att | 12 | 336 | 21.49 | **<0.001** | 0.10 | 0.68 | **<0.001** |
| 7 | Site:Strength:Att | 12 | 336 | 2.14 | **0.014** | 0.01 | 0.68 | **0.032** |

**Supplementary Table 6.** Stimulus detection criterion (related to Supplementary Figure 5b).

| | Effect | $DF_n$ | $DF_d$ | *F* | *p* | $\eta^2_G$ | $\varepsilon$ | *p*[GG] |
|---|---|---|---|---|---|---|---|---|
| | *All experiments* | | | | | | | |
| 1 | Site | 1 | 56 | 0.22 | 0.641 | <0.01 | - | - |
| 2 | Exp | 1 | 56 | 0.53 | 0.470 | <0.01 | - | - |
| 3 | Strength | 6 | 336 | 352.02 | **<0.001** | 0.38 | 0.40 | **<0.001** |
| 4 | Att | 2 | 112 | 11.15 | **<0.001** | 0.01 | 0.63 | **0.001** |
| 5 | Site:Exp | 1 | 56 | <0.01 | 0.987 | <0.01 | - | - |
| 6 | Site:Strength | 6 | 336 | 2.36 | 0.030 | <0.01 | 0.40 | 0.088 |
| 7 | Exp:Strength | 6 | 336 | 24.81 | **<0.001** | 0.04 | 0.40 | **<0.001** |
| 8 | Site:Att | 2 | 112 | 7.13 | **0.001** | 0.01 | 0.63 | **0.006** |
| 9 | Exp:Att | 2 | 112 | 9.35 | **<0.001** | 0.01 | 0.63 | **0.002** |
| 10 | Strength:Att | 12 | 672 | 61.69 | **<0.001** | 0.04 | 0.61 | **<0.001** |
| 11 | Site:Exp:Strength | 6 | 336 | 2.56 | 0.019 | <0.01 | 0.40 | 0.071 |
| 12 | Site:Exp:Att | 2 | 112 | 1.90 | 0.155 | <0.01 | 0.63 | 0.171 |
| 13 | Site:Strength:Att | 12 | 672 | 1.19 | 0.288 | <0.01 | 0.61 | 0.308 |
| 14 | Exp:Strength:Att | 12 | 672 | 2.05 | **0.018** | <0.01 | 0.61 | **0.036** |
| 15 | Site:Exp:Strength:Att | 12 | 672 | 0.90 | 0.550 | <0.01 | 0.61 | 0.523 |
| | *Experiment 1* | | | | | | | |
| 1 | Site | 1 | 28 | 0.09 | 0.773 | <0.01 | - | - |
| 2 | Strength | 6 | 168 | 243.16 | **<0.001** | 0.27 | 0.38 | **<0.001** |
| 3 | Att | 2 | 56 | 15.00 | **<0.001** | 0.04 | 0.64 | **<0.001** |
| 4 | Site:Strength | 6 | 168 | 0.06 | 0.999 | <0.01 | 0.38 | 0.956 |
| 5 | Site:Att | 2 | 56 | 6.84 | **0.002** | 0.02 | 0.64 | **0.008** |
| 6 | Strength:Att | 12 | 336 | 45.43 | **<0.001** | 0.03 | 0.61 | **<0.001** |
| 7 | Site:Strength:Att | 12 | 336 | 1.15 | 0.319 | <0.01 | 0.61 | 0.333 |
| | *Experiment 3* | | | | | | | |
| 1 | Site | 1 | 28 | 0.15 | 0.703 | <0.01 | - | - |
| 2 | Strength | 6 | 168 | 171.11 | **<0.001** | 0.51 | 0.37 | **<0.001** |
| 3 | Att | 2 | 56 | 3.18 | 0.049 | <0.01 | 0.61 | 0.076 |
| 4 | Site:Strength | 6 | 168 | 3.22 | **0.005** | 0.02 | 0.37 | **0.041** |
| 5 | Site:Att | 2 | 56 | 1.05 | 0.357 | <0.01 | 0.61 | 0.328 |
| 6 | Strength:Att | 12 | 336 | 26.73 | **<0.001** | 0.06 | 0.48 | **<0.001** |
| 7 | Site:Strength:Att | 12 | 336 | 1.00 | 0.448 | <0.01 | 0.48 | 0.425 |

**Supplementary Table 7.** Feature detection sensitivity (related to Supplementary Figure 5a).

| | Effect | $DF_n$ | $DF_d$ | *F* | *p* | $\eta_G^2$ | $\varepsilon$ | *p*[GG] |
|---|---|---|---|---|---|---|---|---|
| | *All experiments* | | | | | | | |
| 1 | Site | 1 | 56 | <0.01 | 0.950 | <0.01 | - | - |
| 2 | Exp | 1 | 56 | 1.81 | 0.184 | 0.01 | - | - |
| 3 | Strength | 6 | 336 | 397.97 | **<0.001** | 0.64 | 0.37 | **<0.001** |
| 4 | Att | 2 | 112 | 248.90 | **<0.001** | 0.44 | 0.72 | **<0.001** |
| 5 | Site:Exp | 1 | 56 | <0.01 | 0.983 | <0.01 | - | - |
| 6 | Site:Strength | 6 | 336 | 0.18 | 0.982 | <0.01 | 0.37 | 0.856 |
| 7 | Exp:Strength | 6 | 336 | 6.77 | **<0.001** | 0.03 | 0.37 | **0.001** |
| 8 | Site:Att | 2 | 112 | 0.48 | 0.618 | <0.01 | 0.72 | 0.556 |
| 9 | Exp:Att | 2 | 112 | 5.70 | **0.004** | 0.02 | 0.72 | **0.010** |
| 10 | Strength:Att | 12 | 672 | 47.89 | **<0.001** | 0.10 | 0.74 | **<0.001** |
| 11 | Site:Exp:Strength | 6 | 336 | 1.92 | 0.077 | <0.01 | 0.37 | 0.146 |
| 12 | Site:Exp:Att | 2 | 112 | 1.93 | 0.150 | <0.01 | 0.72 | 0.163 |
| 13 | Site:Strength:Att | 12 | 672 | 1.97 | **0.025** | <0.01 | 0.74 | **0.042** |
| 14 | Exp:Strength:Att | 12 | 672 | 1.39 | 0.165 | <0.01 | 0.74 | 0.190 |
| 15 | Site:Exp:Strength:Att | 12 | 672 | 1.91 | **0.031** | <0.01 | 0.74 | **0.049** |
| | *Experiment 1* | | | | | | | |
| 1 | Site | 1 | 28 | <0.01 | 0.976 | <0.01 | - | - |
| 2 | Strength | 6 | 168 | 264.35 | **<0.001** | 0.64 | 0.31 | **<0.001** |
| 3 | Att | 2 | 56 | 146.49 | **<0.001** | 0.54 | 0.77 | **<0.001** |
| 4 | Site:Strength | 6 | 168 | 2.35 | 0.033 | 0.02 | 0.31 | 0.109 |
| 5 | Site:Att | 2 | 56 | 1.74 | 0.185 | 0.01 | 0.77 | 0.193 |
| 6 | Strength:Att | 12 | 336 | 35.14 | **<0.001** | 0.12 | 0.55 | **<0.001** |
| 7 | Site:Strength:Att | 12 | 336 | 0.93 | 0.522 | <0.01 | 0.56 | 0.485 |
| | *Experiment 3* | | | | | | | |
| 1 | Site | 1 | 28 | <0.01 | 0.954 | <0.01 | - | - |
| 2 | Strength | 6 | 168 | 172.17 | **<0.001** | 0.64 | 0.39 | **<0.001** |
| 3 | Att | 2 | 56 | 102.73 | **<0.001** | 0.33 | 0.65 | **<0.001** |
| 4 | Site:Strength | 6 | 168 | 0.42 | 0.867 | <0.01 | 0.39 | 0.693 |
| 5 | Site:Att | 2 | 56 | 0.53 | 0.594 | <0.01 | 0.65 | 0.520 |
| 6 | Strength:Att | 12 | 336 | 19.06 | **<0.001** | 0.09 | 0.70 | **<0.001** |
| 7 | Site:Strength:Att | 12 | 336 | 2.48 | **0.004** | 0.01 | 0.70 | **0.012** |

**Supplementary Table 8.** Feature detection criterion (related to Supplementary Figure 5b).

| | Effect | $DF_n$ | $DF_d$ | *F* | *p* | $\eta^2_G$ | $\varepsilon$ | *p*[GG] |
|---|---|---|---|---|---|---|---|---|
| | *All experiments* | | | | | | | |
| 1 | Site | 1 | 56 | 0.02 | 0.899 | <0.01 | - | - |
| 2 | Exp | 1 | 56 | 0.67 | 0.417 | 0.01 | - | - |
| 3 | Strength | 6 | 336 | 401.02 | **<0.001** | 0.40 | 0.42 | **<0.001** |
| 4 | Att | 2 | 112 | 11.86 | **<0.001** | 0.01 | 0.84 | **<0.001** |
| 5 | Site:Exp | 1 | 56 | 1.12 | 0.294 | 0.02 | - | - |
| 6 | Site:Strength | 6 | 336 | 0.97 | 0.447 | <0.01 | 0.42 | 0.398 |
| 7 | Exp:Strength | 6 | 336 | 12.95 | **<0.001** | 0.02 | 0.42 | **<00.001** |
| 8 | Site:Att | 2 | 112 | 1.82 | 0.167 | <0.01 | 0.84 | 0.174 |
| 9 | Exp:Att | 2 | 112 | 5.21 | **0.007** | <0.01 | 0.84 | **0.010** |
| 10 | Strength:Att | 12 | 672 | 58.27 | **<0.001** | 0.04 | 0.65 | **<0.001** |
| 11 | Site:Exp:Strength | 6 | 336 | 3.41 | **0.003** | <0.01 | 0.42 | **0.026** |
| 12 | Site:Exp:Att | 2 | 112 | 0.05 | 0.948 | <0.01 | 0.84 | 0.924 |
| 13 | Site:Strength:Att | 12 | 672 | 1.05 | 0.403 | <0.01 | 0.65 | 0.399 |
| 14 | Exp:Strength:Att | 12 | 672 | 1.49 | 0.121 | <0.01 | 0.65 | 0.159 |
| 15 | Site:Exp:Strength:Att | 12 | 672 | 2.02 | **0.020** | <0.01 | 0.65 | **0.044** |
| | *Experiment 1* | | | | | | | |
| 1 | Site | 1 | 28 | 0.64 | 0.429 | 0.02 | - | - |
| 2 | Strength | 6 | 168 | 264.35 | **<0.001** | 0.33 | 0.31 | **<0.001** |
| 3 | Att | 2 | 56 | 6.76 | **0.002** | 0.02 | 0.83 | **0.004** |
| 4 | Site:Strength | 6 | 168 | 2.35 | 0.033 | <0.01 | 0.31 | 0.109 |
| 5 | Site:Att | 2 | 56 | 0.54 | 0.588 | <0.01 | 0.83 | 0.556 |
| 6 | Strength:Att | 12 | 336 | 35.14 | **<0.001** | 0.04 | 0.56 | **<0.001** |
| 7 | Site:Strength:Att | 12 | 336 | 0.93 | 0.522 | <0.01 | 0.56 | 0.485 |
| | *Experiment 3* | | | | | | | |
| 1 | Site | 1 | 28 | 0.48 | 0.495 | 0.01 | - | - |
| 2 | Strength | 6 | 168 | 183.35 | **<0.001** | 0.47 | 0.43 | **<0.001** |
| 3 | Att | 2 | 56 | 13.61 | **<0.001** | 0.01 | - | **-** |
| 4 | Site:Strength | 6 | 168 | 2.12 | 0.053 | 0.01 | 0.43 | 0.113 |
| 5 | Site:Att | 2 | 56 | 2.08 | 0.135 | <0.01 | - | - |
| 6 | Strength:Att | 12 | 336 | 27.09 | **<0.001** | 0.05 | - | - |
| 7 | Site:Strength:Att | 12 | 336 | 1.86 | **0.038** | <0.01 | - | - |

**Supplementary Table 9.** Unequal variance stimulus detection sensitivity (related to Supplementary Figure 6a).

| | Effect | $DF_n$ | $DF_d$ | *F* | *p* | $\eta_G^2$ | $\varepsilon$ | *p*[GG] |
|---|---|---|---|---|---|---|---|---|
| | *All experiments* | | | | | | | |
| 1 | Site | 1 | 56 | 2.76 | 0.102 | 0.02 | - | - |
| 2 | Exp | 1 | 56 | 5.50 | 0.023 | 0.04 | - | - |
| 3 | Strength | 6 | 336 | 209.87 | **<0.001** | 0.50 | 0.44 | **<0.001** |
| 4 | Att | 2 | 112 | 201.24 | **<0.001** | 0.36 | 0.82 | **<0.001** |
| 5 | Site:Exp | 1 | 56 | 0.27 | 0.603 | <0.01 | - | - |
| 6 | Site:Strength | 6 | 336 | 0.49 | 0.818 | <0.01 | 0.44 | 0.667 |
| 7 | Exp:Strength | 6 | 336 | 7.29 | **<0.001** | 0.03 | 0.44 | **<0.001** |
| 8 | Site:Att | 2 | 112 | 2.38 | 0.098 | <0.01 | 0.82 | 0.109 |
| 9 | Exp:Att | 2 | 112 | 4.53 | **0.013** | 0.01 | 0.82 | **0.019** |
| 10 | Strength:Att | 12 | 672 | 23.79 | **<0.001** | 0.08 | 0.75 | **<0.001** |
| 11 | Site:Exp:Strength | 6 | 336 | 0.29 | 0.941 | <0.01 | 0.44 | 0.806 |
| 12 | Site:Exp:Att | 2 | 112 | 1.24 | 0.292 | <0.01 | 0.82 | 0.288 |
| 13 | Site:Strength:Att | 12 | 672 | 1.14 | 0.323 | <0.01 | 0.75 | 0.332 |
| 14 | Exp:Strength:Att | 12 | 672 | 1.35 | 0.185 | <0.01 | 0.75 | 0.208 |
| 15 | Site:Exp:Strength:Att | 12 | 672 | 1.58 | 0.094 | <0.01 | 0.75 | 0.120 |
| | *Experiment 1* | | | | | | | |
| 1 | Site | 1 | 28 | 2.76 | 0.108 | 0.03 | - | - |
| 2 | Strength | 6 | 168 | 106.35 | **<0.001** | 0.49 | 0.43 | **<0.001** |
| 3 | Att | 2 | 56 | 107.61 | **<0.001** | 0.44 | 0.82 | **<0.001** |
| 4 | Site:Strength | 6 | 168 | 0.25 | 0.957 | <0.01 | 0.43 | 0.831 |
| 5 | Site:Att | 2 | 56 | 2.84 | 0.07 | 0.02 | 0.82 | 0.079 |
| 6 | Strength:Att | 12 | 336 | 14.21 | **<0.001** | 0.09 | 0.50 | **<0.001** |
| 7 | Site:Strength:Att | 12 | 336 | 0.69 | 0.765 | <0.01 | 050 | 0.660 |
| | *Experiment 3* | | | | | | | |
| 1 | Site | 1 | 28 | 0.57 | 0.456 | <0.01 | - | - |
| 2 | Strength | 6 | 168 | 110.36 | **<0.001** | 0.52 | 0.40 | **<0.001** |
| 3 | Att | 2 | 56 | 95.30 | **<0.001** | 0.28 | 0.80 | **<0.001** |
| 4 | Site:Strength | 6 | 168 | 0.50 | 0.811 | <0.01 | 0.40 | 0.645 |
| 5 | Site:Att | 2 | 56 | 0.15 | 0.861 | <0.01 | 0.80 | 0.815 |
| 6 | Strength:Att | 12 | 336 | 11.27 | **<0.001** | 0.08 | 0.72 | **<0.001** |
| 7 | Site:Strength:Att | 12 | 336 | 1.90 | 0.034 | 0.01 | 0.72 | 0.056 |

**Supplementary Table 10.** Unequal variance stimulus detection criterion (related to Supplementary Figure 6b).

| | Effect | $DF_n$ | $DF_d$ | *F* | *p* | $\eta_G^2$ | $\varepsilon$ | *p*[GG] |
|---|---|---|---|---|---|---|---|---|
| | *All experiments* | | | | | | | |
| 1 | Site | 1 | 56 | 0.30 | 0.586 | <0.01 | - | - |
| 2 | Exp | 1 | 56 | 0.42 | 0.520 | <0.01 | - | - |
| 3 | Strength | 6 | 336 | 250.30 | **<0.001** | 0.38 | 0.42 | **<0.001** |
| 4 | Att | 2 | 112 | 4.65 | **0.012** | <0.01 | 0.70 | **0.023** |
| 5 | Site:Exp | 1 | 56 | 0.03 | 0.864 | <0.01 | - | - |
| 6 | Site:Strength | 6 | 336 | 3.88 | **0.001** | <0.01 | 0.42 | **0.016** |
| 7 | Exp:Strength | 6 | 336 | 24.11 | **<0.001** | 0.06 | 0.42 | **<0.001** |
| 8 | Site:Att | 2 | 112 | 6.20 | **0.003** | 0.01 | 0.70 | **0.008** |
| 9 | Exp:Att | 2 | 112 | 5.57 | **0.005** | <0.01 | 0.70 | **0.012** |
| 10 | Strength:Att | 12 | 672 | 33.19 | **<0.001** | 0.04 | 0.61 | **<0.001** |
| 11 | Site:Exp:Strength | 6 | 336 | 1.36 | 0.231 | <0.01 | 0.42 | 0.261 |
| 12 | Site:Exp:Att | 2 | 112 | 2.08 | 0.129 | <0.01 | 0.70 | 0.146 |
| 13 | Site:Strength:Att | 12 | 672 | 1.40 | 0.162 | <0.01 | 0.61 | 0.202 |
| 14 | Exp:Strength:Att | 12 | 672 | 2.55 | **0.003** | <0.01 | 0.61 | **0.013** |
| 15 | Site:Exp:Strength:Att | 12 | 672 | 1.44 | 0.143 | <0.01 | 0.51 | 0.184 |
| | *Experiment 1* | | | | | | | |
| 1 | Site | 1 | 28 | 0.24 | 0.626 | <0.01 | - | - |
| 2 | Strength | 6 | 168 | 99.71 | **<0.001** | 0.25 | 0.41 | **<0.001** |
| 3 | Att | 2 | 56 | 6.88 | **0.002** | 0.03 | 0.71 | **0.006** |
| 4 | Site:Strength | 6 | 168 | 1.43 | 0.205 | <0.01 | 0.41 | 0.244 |
| 5 | Site:Att | 2 | 56 | 6.20 | **0.004** | 0.02 | 0.71 | **0.010** |
| 6 | Strength:Att | 12 | 336 | 11.09 | **<0.001** | 0.03 | 0.43 | **<0.001** |
| 7 | Site:Strength:Att | 12 | 336 | 1.83 | 0.043 | <0.01 | 0.43 | 0.109 |
| | *Experiment 3* | | | | | | | |
| 1 | Site | 1 | 28 | 0.08 | 0.786 | <0.01 | - | - |
| 2 | Strength | 6 | 168 | 157.43 | **<0.001** | 0.50 | 0.36 | **<0.001** |
| 3 | Att | 2 | 56 | 2.21 | 0.120 | <0.01 | 0.68 | 0.139 |
| 4 | Site:Strength | 6 | 168 | 3.26 | **0.005** | 0.02 | 0.36 | **0.041** |
| 5 | Site:Att | 2 | 56 | 0.78 | 0.466 | <0.01 | 0.68 | 0.421 |
| 6 | Strength:Att | 12 | 336 | 24.84 | **<0.001** | 0.06 | 0.49 | **<0.001** |
| 7 | Site:Strength:Att | 12 | 336 | 1.00 | 0.452 | <0.01 | 0.49 | 0.429 |

**Supplementary Table 11.** Unequal variance feature detection sensitivity (related to Supplementary Figure 6a).

| | Effect | $DF_n$ | $DF_d$ | *F* | *p* | $\eta^2_G$ | $\varepsilon$ | *p*[GG] |
|---|---|---|---|---|---|---|---|---|
| | *All experiments* | | | | | | | |
| 1 | Site | 1 | 56 | 0.15 | 0.703 | <0.01 | - | - |
| 2 | Exp | 1 | 56 | 1.89 | 0.174 | 0.01 | - | - |
| 3 | Strength | 6 | 336 | 169.03 | **<0.001** | 0.41 | 0.46 | **<0.001** |
| 4 | Att | 2 | 112 | 107.66 | **<0.001** | 0.26 | 0.80 | **<0.001** |
| 5 | Site:Exp | 1 | 56 | 0.12 | 0.731 | <0.01 | - | - |
| 6 | Site:Strength | 6 | 336 | 0.25 | 0.960 | <0.01 | 0.46 | 0.847 |
| 7 | Exp:Strength | 6 | 336 | 4.51 | **<0.001** | 0.02 | 0.46 | **0.006** |
| 8 | Site:Att | 2 | 112 | 2.05 | 0.134 | <0.01 | 0.80 | 0.144 |
| 9 | Exp:Att | 2 | 112 | 6.66 | **0.002** | 0.02 | 0.80 | **0.004** |
| 10 | Strength:Att | 12 | 672 | 8.83 | **<0.001** | 0.03 | 0.61 | **<0.001** |
| 11 | Site:Exp:Strength | 6 | 336 | 0.50 | 0.812 | <0.01 | 0.46 | 0.670 |
| 12 | Site:Exp:Att | 2 | 112 | 3.69 | **0.028** | 0.01 | 0.80 | **0.038** |
| 13 | Site:Strength:Att | 12 | 672 | 1.24 | 0.248 | <0.01 | 0.61 | 0.275 |
| 14 | Exp:Strength:Att | 12 | 672 | 1.25 | 0.245 | <0.01 | 0.61 | 0.272 |
| 15 | Site:Exp:Strength:Att | 12 | 672 | 1.15 | 0.316 | <0.01 | 0.61 | 0.330 |
| | *Experiment 1* | | | | | | | |
| 1 | Site | 1 | 28 | <0.01 | 0.981 | <0.01 | - | - |
| 2 | Strength | 6 | 168 | 75.51 | **<0.001** | 0.33 | 0.43 | **<0.001** |
| 3 | Att | 2 | 56 | 57.77 | **<0.001** | 0.34 | 0.80 | **<0.001** |
| 4 | Site:Strength | 6 | 168 | 0.29 | 0.939 | <0.01 | 0.43 | 0.798 |
| 5 | Site:Att | 2 | 56 | 3.90 | **0.026** | 0.03 | 0.80 | **0.036** |
| 6 | Strength:Att | 12 | 336 | 6.27 | **<0.001** | 0.04 | 0.48 | **<0.001** |
| 7 | Site:Strength:Att | 12 | 336 | 0.84 | 0.610 | <0.01 | 0.48 | 0.536 |
| | *Experiment 3* | | | | | | | |
| 1 | Site | 1 | 28 | 0.31 | 0.584 | <0.01 | - | - |
| 2 | Strength | 6 | 168 | 94.67 | **<0.001** | 0.49 | 0.44 | **<0.001** |
| 3 | Att | 2 | 56 | 55.57 | **<0.001** | 0.17 | 0.80 | **<0.001** |
| 4 | Site:Strength | 6 | 168 | 0.43 | 0.862 | <0.01 | 0.44 | 0.712 |
| 5 | Site:Att | 2 | 56 | 0.22 | 0.807 | <0.01 | 0.80 | 0.758 |
| 6 | Strength:Att | 12 | 336 | 4.19 | **<0.001** | 0.04 | 0.59 | **<0.001** |
| 7 | Site:Strength:Att | 12 | 336 | 1.45 | 0.144 | 0.01 | 0.59 | 0.188 |

**Supplementary Table 12.** Unequal variance feature detection criterion (related to Supplementary Figure 6b).

| | Effect | $DF_n$ | $DF_d$ | *F* | *p* | $\eta_G^2$ | $\varepsilon$ | *p*[GG] |
|---|---|---|---|---|---|---|---|---|
| | *All experiments* | | | | | | | |
| 1 | Site | 1 | 56 | <0.01 | 0.973 | <0.01 | - | - |
| 2 | Exp | 1 | 56 | 0.38 | 0.539 | <0.01 | - | - |
| 3 | Strength | 6 | 336 | 322.36 | **<0.001** | 0.40 | 0.47 | **<0.001** |
| 4 | Att | 2 | 112 | 3.42 | **0.036** | <0.01 | 0.84 | **0.045** |
| 5 | Site:Exp | 1 | 56 | 0.63 | 0.430 | <0.01 | - | - |
| 6 | Site:Strength | 6 | 336 | 0.98 | 0.441 | <0.01 | 0.47 | 0.402 |
| 7 | Exp:Strength | 6 | 336 | 13.90 | **<0.001** | 0.03 | 0.47 | **<0.001** |
| 8 | Site:Att | 2 | 112 | 0.30 | 0.741 | <0.01 | 0.84 | 0.702 |
| 9 | Exp:Att | 2 | 112 | 3.65 | **0.029** | <0.01 | 0.84 | **0.037** |
| 10 | Strength:Att | 12 | 672 | 29.21 | **<0.001** | 0.05 | 0.65 | **<0.001** |
| 11 | Site:Exp:Strength | 6 | 336 | 2.04 | 0.059 | <0.01 | 0.47 | 0.114 |
| 12 | Site:Exp:Att | 2 | 112 | 0.53 | 0.588 | <0.01 | 0.84 | 0.557 |
| 13 | Site:Strength:Att | 12 | 672 | 0.82 | 0.633 | <0.01 | 0.65 | 0.584 |
| 14 | Exp:Strength:Att | 12 | 672 | 2.20 | **0.010** | <0.01 | 0.65 | **0.028** |
| 15 | Site:Exp:Strength:Att | 12 | 672 | 1.45 | 0.139 | <0.01 | 0.65 | 0.177 |
| | *Experiment 1* | | | | | | | |
| 1 | Site | 1 | 28 | 0.28 | 0.598 | <0.01 | - | - |
| 2 | Strength | 6 | 168 | 175.96 | **<0.001** | 0.32 | 0.43 | **<0.001** |
| 3 | Att | 2 | 56 | 1.88 | 0.162 | <0.01 | 0.84 | 0.170 |
| 4 | Site:Strength | 6 | 168 | 0.89 | 0.504 | <0.01 | 0.43 | 0.437 |
| 5 | Site:Att | 2 | 56 | 0.03 | 0.972 | <0.01 | 0.84 | 0.954 |
| 6 | Strength:Att | 12 | 336 | 11.90 | **<0.001** | 0.03 | 0.41 | **<0.001** |
| 7 | Site:Strength:Att | 12 | 336 | 0.42 | 0.957 | <0.01 | 0.41 | 0.834 |
| | *Experiment 3* | | | | | | | |
| 1 | Site | 1 | 28 | 0.35 | 0.559 | <0.01 | - | - |
| 2 | Strength | 6 | 168 | 164.18 | **<0.001** | 0.48 | 0.46 | **<0.001** |
| 3 | Att | 2 | 56 | 10.31 | **<0.001** | 0.01 | - | - |
| 4 | Site:Strength | 6 | 168 | 1.82 | 0.097 | 0.01 | 0.46 | 0.154 |
| 5 | Site:Att | 2 | 56 | 2.01 | 0.144 | <0.01 | - | - |
| 6 | Strength:Att | 12 | 336 | 18.81 | **<0.001** | 0.07 | 0.56 | **<0.001** |
| 7 | Site:Strength:Att | 12 | 336 | 1.72 | 0.062 | <0.01 | 0.56 | 0.110 |

**Supplementary Table 13.** AUC of stimulus visibility (related to Figure 2d and Figure 5 top row).

| | | Effect | $DF_r$ | $DF_d$ | *F* | *p* | $\eta_G^2$ | $\varepsilon$ | *p*[GG] |
|---|---|---|---|---|---|---|---|---|---|
| *All experiments* | 1 | Site | 1 | 111 | 0.26 | 0.613 | <0.01 | - | - |
| | 2 | Exp | 3 | 111 | 67.57 | **<0.001** | 0.61 | - | - |
| | 3 | Att | 2 | 222 | 117.24 | **<0.001** | 0.14 | 0.91 | **<0.001** |
| | 4 | Site:Exp | 3 | 111 | 3.52 | **0.017** | 0.08 | - | - |
| | 5 | Site:Att | 2 | 222 | 0.81 | 0.447 | <0.01 | 0.91 | 0.436 |
| | 6 | Exp:Att | 6 | 222 | 1.75 | 0.110 | <0.01 | 0.91 | 0.118 |
| | 7 | Site:Exp:Att | 6 | 222 | 1.46 | 0.193 | <0.01 | 0.91 | 0.200 |
| *Experiment 1* | 1 | Site | 1 | 28 | 4.79 | **0.037** | 0.14 | - | - |
| | 2 | Att | 2 | 56 | 36.06 | **<0.001** | 0.10 | 0.77 | **<0.001** |
| | 3 | Site:Att | 2 | 56 | 3.03 | 0.056 | 0.01 | 0.77 | 0.071 |
| *Experiment 2* | 1 | Site | 1 | 28 | 0.04 | 0.840 | <0.01 | - | - |
| | 2 | Att | 2 | 56 | 49.41 | **<0.001** | 0.42 | 0.61 | **<0.001** |
| | 3 | Site:Att | 2 | 56 | 0.50 | 0.608 | <0.01 | 0.61 | 0.52 |
| *Experiment 3* | 1 | Site | 1 | 28 | 1.77 | 0.194 | 0.05 | - | - |
| | 2 | Att | 2 | 56 | 15.84 | **<0.001** | 0.06 | - | - |
| | 3 | Site:Att | 2 | 56 | 1.03 | 0.362 | <0.01 | - | - |
| *Experiment 4* | 1 | Site | 1 | 27 | 0.34 | 0.568 | <0.01 | - | - |
| | 2 | Att | 2 | 54 | 39.80 | **<0.001** | 0.37 | - | - |
| | 3 | Site:Att | 2 | 54 | 0.84 | 0.438 | 0.01 | - | - |

**Supplementary Table 14.** AUC of feature visibility (related to Figure 3d and Figure 5 bottom row).

| | | Effect | $DF_r$ | $DF_d$ | *F* | *p* | $\eta_G^2$ | $\varepsilon$ | *p*[GG] |
|---|---|---|---|---|---|---|---|---|---|
| *All experiments* | 1 | Site | 1 | 84 | 0.42 | 0.520 | <0.01 | - | - |
| | 2 | Exp | 2 | 84 | 7.26 | **<0.001** | 0.13 | - | - |
| | 3 | Att | 2 | 168 | 6.54 | **0.002** | <0.01 | 0.78 | **0.004** |
| | 4 | Site:Exp | 3 | 84 | 1.80 | 0.172 | 0.04 | - | - |
| | 5 | Site:Att | 2 | 168 | 0.32 | 0.728 | <0.01 | 0.78 | 0.674 |
| | 6 | Exp:Att | 4 | 168 | 7.67 | **<0.001** | 0.02 | 0.78 | **<0.001** |
| | 7 | Site:Exp:Att | 4 | 168 | 1.58 | 0.183 | <0.01 | 0.78 | 0.196 |
| *Experiment 1* | 1 | Site | 1 | 28 | 2.01 | 0.168 | 0.05 | - | - |
| | 2 | Att | 2 | 56 | 14.63 | **<0.001** | 0.09 | 0.62 | **<0.001** |
| | 3 | Site:Att | 2 | 56 | 1.18 | 0.314 | <0.01 | 0.62 | 0.297 |
| *Experiment 2* | 1 | Site | 1 | 28 | 1.27 | 0.269 | 0.04 | - | - |
| | 2 | Att | 2 | 56 | 2.22 | 0.118 | <0.01 | 0.64 | 0.140 |
| | 3 | Site:Att | 2 | 56 | 1.48 | 0.238 | <0.01 | 0.64 | 0.239 |
| *Experiment 3* | 1 | Site | 1 | 28 | 1.14 | 0.294 | 0.04 | - | - |
| | 2 | Att | 2 | 56 | 5.27 | **0.008** | 0.02 | 0.89 | **0.011** |
| | 3 | Site:Att | 2 | 56 | 0.72 | 0.492 | <0.01 | 0.89 | 0.477 |

**Supplementary Table 15.** AMI (related to Figure 6).

| | | Effect | $DF_r$ | $DF_d$ | *F* | *p* | $\eta_G^2$ |
|---|---|---|---|---|---|---|---|
| *Stimulus-level* | 1 | Site | 1 | 110 | 0.064 | 0.800 | <0.01 |
| | 2 | Strength regime | 1 | 110 | 197.65 | **<0.001** | 0.64 |
| | 3 | Stimulus type | 1 | 110 | 3.58 | 0.061 | 0.03 |
| | 4 | Site:Strength regime | 1 | 110 | <0.01 | 0.963 | <0.01 |
| | 5 | Site:Stimulus type | 1 | 110 | 0.03 | 0.876 | <0.01 |
| | 6 | Strength regime:Stimulus type | 1 | 110 | 0.15 | 0.697 | <0.01 |
| | 7 | Site:Strength regime:Stimulus type | 1 | 110 | 1.41 | 0.238 | 0.01 |
| *Feature-level* | 1 | Site | 1 | 84 | 0.55 | 0.459 | <0.01 |
| | 2 | Exp | 2 | 84 | 20.43 | **<0.001** | 0.33 |
| | 3 | Site:Exp | 2 | 84 | 0.73 | 0.485 | 0.02 |

**Supplementary Table 16.** AMI by analytic pipeline (related to Supplementary Figure 8).

| | | Effect | $DF_r$ | $DF_d$ | *F* | *p* | $\eta_G^2$ |
|---|---|---|---|---|---|---|---|
| *Stimulus-level* | 1 | Site | 1 | 220 | 0.74 | 0.390 | <0.01 |
| | 2 | Exp | 3 | 220 | 130.23 | **<0.001** | 0.64 |
| | 3 | Pipe | 1 | 220 | 0.02 | 0.895 | <0.01 |
| | 4 | Site:Exp | 3 | 220 | 0.75 | 0.525 | <0.01 |
| | 5 | Site:Pipe | 1 | 220 | 0.26 | 0.613 | <0.01 |
| | 6 | Exp:Pipe | 3 | 220 | 0.15 | 0.931 | <0.01 |
| | 7 | Site:Exp:Pipe | 3 | 220 | 0.07 | 0.976 | <0.01 |
| *Feature-level* | 1 | Site | 1 | 168 | 2.15 | 0.145 | 0.01 |
| | 2 | Exp | 2 | 168 | 45.11 | **<0.001** | 0.35 |
| | 3 | Pipe | 1 | 168 | 0.20 | 0.657 | <0.01 |
| | 4 | Site:Exp | 2 | 168 | 2.13 | 0.122 | 0.03 |
| | 5 | Site:Pipe | 1 | 168 | 0.14 | 0.706 | <0.01 |
| | 6 | Exp:Pipe | 2 | 168 | 0.09 | 0.911 | <0.01 |
| | 7 | Site:Exp:Pipe | 2 | 168 | 0.06 | 0.943 | <0.01 |

**Supplementary Table 17.** Thresholds.

| | | Effect | $DF_r$ | $DF_d$ | *F* | *p* | $\eta_G^2$ |
|---|---|---|---|---|---|---|---|
| *All experiments* | 1 | Site | 1 | 112 | 0.22 | 0.637 | <0.01 |
| | 2 | Exp | 3 | 112 | 36.31 | **<0.001** | 0.49 |
| | 3 | Site:Exp | 3 | 112 | 2.53 | 0.061 | 0.06 |
| *Experiment 1* | 1 | Site | 1 | 28 | 0.10 | 0.752 | <0.01 |
| *Experiment 2* | 1 | Site | 1 | 28 | 1.51 | 0.229 | 0.05 |
| *Experiment 3* | 1 | Site | 1 | 28 | 2.26 | 0.144 | 0.08 |
| *Experiment 4* | 1 | Site | 1 | 28 | 1.84 | 0.185 | 0.06 |

**Supplementary Table 18.** Stimulus-anchored attentional threshold shifts for performance vs. stimulus visibility (Figure 8).

| | | Effect | $DF_r$ | $DF_d$ | $F$ | $p$ | $\eta_G^2$ | $\varepsilon$ | $p$[GG] |
|---|---|---|---|---|---|---|---|---|---|
| *All experiments* | 1 | Site | 1 | 107 | 0.50 | 0.482 | <0.01 | - | - |
| | 2 | Exp | 3 | 107 | 2.17 | 0.096 | 0.02 | - | - |
| | 3 | Measure | 1 | 107 | 49.10 | **<0.001** | 0.22 | - | - |
| | 4 | Site:Exp | 3 | 107 | 0.44 | 0.723 | <0.01 | - | - |
| | 5 | Site:Measure | 1 | 107 | 0.002 | 0.964 | <0.01 | - | - |
| | 6 | Exp:Measure | 3 | 107 | 29.72 | **<0.001** | 0.34 | - | - |
| | 7 | Site:Exp:Measure | 3 | 107 | 0.006 | 0.999 | <0.01 | - | - |
| *Experiment 1* | 1 | Site | 1 | 26 | 0.068 | 0.797 | <0.01 | - | - |
| | 2 | Measure | 1 | 26 | 85.88 | **<0.001** | 0.48 | - | - |
| | 3 | Site:Measure | 1 | 26 | 1.09 | 0.307 | 0.01 | - | - |
| *Experiment 2* | 1 | Site | 1 | 26 | 0.195 | 0.652 | <0.01 | - | - |
| | 2 | Measure | 1 | 26 | 84.85 | **<0.001** | 0.67 | - | - |
| | 3 | Site:Measure | 1 | 26 | 0.29 | 0.595 | <0.01 | - | - |
| *Experiment 3* | 1 | Site | 1 | 27 | 0.22 | 0.640 | <0.01 | - | - |
| | 2 | Measure | 1 | 27 | 17.57 | **<0.001** | 0.26 | - | - |
| | 3 | Site:Measure | 1 | 27 | 0.17 | 0.682 | <0.01 | - | - |
| *Experiment 4* | 1 | Site | 1 | 25 | 0.72 | 0.403 | 0.01 | - | - |
| | 2 | Measure | 1 | 25 | 31.39 | **<0.001** | 0.44 | - | - |
| | 3 | Site:Measure | 1 | 25 | 0.001 | 0.978 | <0.01 | - | - |

"Measure" indicates the factor of perceptual measure.

**Supplementary Table 19.** Stimulus-anchored attentional threshold shifts for performance vs. feature visibility (Figure 8).

| | | Effect | $DF_r$ | $DF_d$ | $F$ | $p$ | $\eta_G^2$ | $\varepsilon$ | $p$[GG] |
|---|---|---|---|---|---|---|---|---|---|
| *All experiments* | 1 | Site | 1 | 80 | 0.01 | 0.942 | <0.01 | - | - |
| | 2 | Exp | 2 | 80 | 5.15 | **0.008** | 0.08 | - | - |
| | 3 | Measure | 1 | 80 | 51.91 | **<0.001** | 0.19 | - | - |
| | 4 | Site:Exp | 2 | 80 | 0.16 | 0.850 | <0.01 | - | - |
| | 5 | Site:Measure | 1 | 80 | 0.03 | 0.866 | <0.01 | - | - |
| | 6 | Exp:Measure | 2 | 80 | 8.98 | **<0.001** | 0.08 | - | - |
| | 7 | Site:Exp:Measure | 2 | 80 | 1.13 | 0.330 | 0.01 | - | - |
| *Experiment 1* | 1 | Site | 1 | 27 | 0.14 | 0.714 | <0.01 | - | - |
| | 2 | Measure | 1 | 27 | 62.11 | **<0.001** | 0.47 | - | - |
| | 3 | Site:Measure | 1 | 27 | 0.23 | 0.635 | <0.01 | - | - |
| *Experiment 2* | 1 | Site | 1 | 26 | 0.04 | 0.849 | <0.01 | - | - |
| | 2 | Measure | 1 | 26 | 2.68 | 0.114 | 0.03 | - | - |
| | 3 | Site:Measure | 1 | 26 | 2.68 | 0.114 | 0.03 | - | - |
| *Experiment 3* | 1 | Site | 1 | 27 | 0.17 | 0.682 | <0.01 | - | - |
| | 2 | Measure | 1 | 27 | 9.64 | **0.004** | 0.14 | - | - |
| | 3 | Site:Measure | 1 | 27 | 0.22 | 0.645 | <0.01 | - | - |
